# Improved ^1^H NMR and ATR-FTIR approaches for structural characterization of high-G-block and high-M-block alginic acids isolated from sugar kelp by a two-step gelation method

**DOI:** 10.64898/2026.10.21.760866

**Authors:** Xiaohui Xing, Barinder Bajwa, Kristin Elizabeth Low, Jeffrey P. Tingley, Vincent Weiler, Tony Montina, Spencer C. Serin, D. Wade Abbott

## Abstract

Two low-molecular-weight alginic acid fractions, G-block-rich SKP-G1 and M-block-rich SKP-G2, were sequentially isolated from a hot-water extract of sugar kelp (*Saccharina latissima*) alcohol-insoluble residue by acetic acid-induced and subsequent freeze–thaw-induced gelation. High-field (700 MHz) 2D ^1^H–^1^H COSY and TOCSY spectroscopy enabled comprehensive cross-peak assignments for selected alginate diad and triad sequences, supporting interpretation of 1D ^1^H NMR spectra. Sequence frequencies were combined with previously established inequalities [1] to calculate constrained frequency ranges for homoblocks beyond the triad level. For selected homoblock lengths up to a degree of polymerization of 18, comparison of the experimentally derived frequency ranges with frequencies predicted by chain-growth statistical models ruled out the Bernoulli model for both fractions and the first-order Markov model for SKP-G1. Applying the recently developed logarithmic cosine transformation (LCT) algorithm [2] to first- and second-derivative ATR-FTIR spectra enhanced the visualization of subtle spectral features and produced distinct fingerprints for the two fractions. As a proof of concept, LCT-transformed derivative ATR-FTIR spectra yielded distinct score distributions for the fractions in principal component analysis, suggesting their potential for chemometric differentiation of alginates with different block structures. In conclusion, extending ^1^H NMR-based sequence analysis to calculate constrained frequency ranges for homoblock polyads beyond the triad level, together with LCT-transformed derivative ATR-FTIR fingerprinting, expands the analytical toolkit for alginate block structure characterization.

**Highlights:**

- Low-molecular-weight high-G and high-M alginic acids were isolated from sugar kelp by two-step gelation.
- Logarithmic cosine transformation enabled extraction of key features from derivative ATR-FTIR spectra.
- High-field (700 MHz) 2D ^1^H–^1^H NMR facilitated complete assignment of alginate homoblock triad signals.
- Alginate homoblock frequency ranges were calculated from 1D ^1^H NMR data using an improved approach.
- Alginate homoblock frequency ranges were used to evaluate Bernoulli and Markov chain-growth models.

**Graphical abstract:** 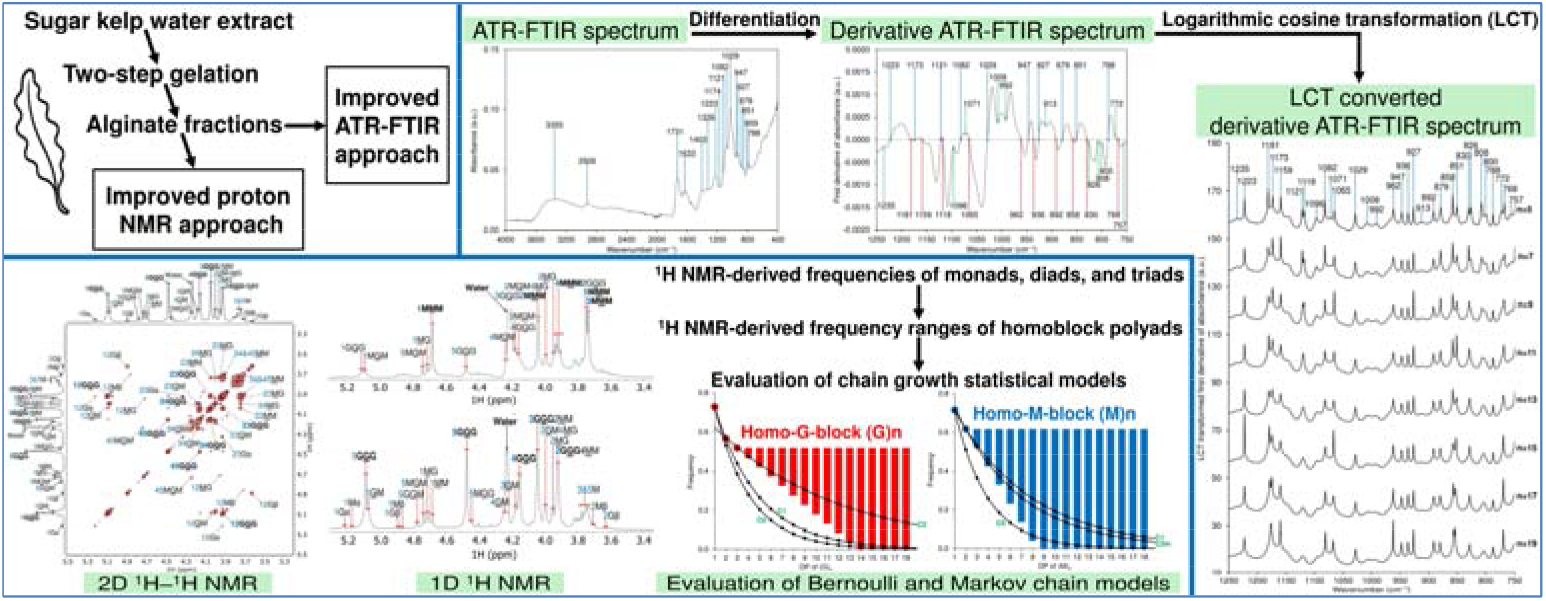

## 1. Introduction

*Saccharina latissima*, commonly known as sugar kelp, is a perennial macroscopic brown alga widely distributed across the Northern Hemisphere, from the high Arctic to cold-temperate regions along the Atlantic and Pacific coasts, with its southernmost range extending to the Iberian Peninsula in Europe, New Jersey on the Atlantic coast of North America, central California on the Pacific coast, and the Korean Peninsula in the Far East [3–5]. As a key primary producer, sugar kelp supports marine ecosystems by providing both food and habitat for diverse organisms [6]. Rich in carbohydrates, proteins, lipids, polyphenols, and other bioactive compounds, it exhibits antioxidant, antitumor, immunomodulatory, antiviral, and anti-inflammatory properties, contributing to its nutritional and functional value [7]. Its major carbohydrate-related constituents include mannitol, alginic acids, fucoidans, laminarins, and cellulose, which can be extracted as value-added products for diverse applications [7–9]. Its ecological and economic significance has led to the expansion of sugar kelp farming in Western countries such as Canada [10], the USA [11], Norway [8], and the UK [12].

Alginic acids are anionic polysaccharides consisting of linear chains of 1,4-linked β-D-mannuronic acid (M) and α-L-guluronic acid (G) residues, arranged in homopolymeric (MM- or GG-rich) or heteropolymeric (MG- or GM-rich) blocks [13]. Both alginic acid and its salt form, alginate, are widely utilized in the food, pharmaceutical, and other industries as bioactive carbohydrates, dietary fibers, functional additives, and biomaterials [13, 14]. First extracted from brown seaweed by E.C.C. Stanford in 1881, alginate has since been conventionally produced via alkaline extraction, with insoluble alginic acid or calcium alginate precipitated to facilitate collection [15]. Calcium alginate is typically converted to alginic acid by treatment with a mineral acid and then transformed into water-soluble sodium alginate via reaction with sodium hydroxide or sodium carbonate [16]. Conventional alkaline extraction is often preceded by pretreatment with solutions of formaldehyde, hydrochloric acid, or alcohol [17]. Microwave-, ultrasound-, and enzyme-assisted extraction techniques have attracted interest for reducing processing time and enhancing yield [17].

Attenuated total reflectance Fourier transform infrared spectroscopy (ATR-FTIR) is a useful tool for the compositional and structural characterization of alginate and alginic acid, enabling rapid, non-destructive identification of functional groups and molecular interactions associated with processes such as gelation, hydration, and crosslinking [18–20]. First- and second-derivative spectra enhance spectral resolution and reduce baseline shifts, facilitating accurate peak assignment and the detection of subtle structural variations [20–23]. Recognized as a powerful fingerprinting technique, ATR-FTIR provides spectral features for chemometric analysis, including principal component analysis (PCA) [24, 25]. These features facilitate precise classification, authentication, and quality control, making ATR-FTIR a valuable tool for both research and industrial applications [24, 25].

Solution-state ^1^H NMR is the gold standard for characterizing purified alginate, enabling determination of the monosaccharide composition (M/G ratio) and the full set of diad and G-centered triad frequencies via integration of specific peaks in the anomeric region of the ^1^H NMR spectrum [24, 26]. Although ^13^C NMR can provide complete diad and triad frequencies, including M-centered triads [27, 28], it is less commonly used because of longer acquisition times and higher spectral noise. NMR-derived diad and triad frequencies also enable analysis of homoblock polyad frequency ranges using recently derived inequality relations [1], facilitating evaluation of chain-growth statistical models, such as Bernoulli processes and Markov chains, for homoblock elongation [1, 28]. Advances in high-field ^1^H NMR have improved spectral resolution, and 2D techniques such as ^1^H–^1^H correlation spectroscopy (COSY) and total correlation spectroscopy (TOCSY) facilitate the assignment of alginate signals [29–32].

Prior to ^1^H NMR analysis, the purity and molecular weight distribution of alginate samples are usually assessed using chromatographic methods. High-performance size-exclusion chromatography (HPSEC), combined with refractive index (RI) and multi-angle light scattering (MALS) detectors, enables direct measurement of molecular weights based on the physical principle of static light scattering, obviating the need for conventional column calibration [33, 34]. Monosaccharide composition is typically determined by depolymerizing alginic acid under acidic conditions (*e.g*., acid methanolysis [34], aqueous acid hydrolysis [35, 36], or a combination thereof [37]), followed by chromatographic analysis using techniques such as high-performance anion-exchange chromatography with pulsed amperometric detection (HPAEC-PAD) [34] or gas chromatography coupled with mass spectrometry and/or flame ionization detection (GC-MS/FID) [38, 39]. Although quantitative depolymerization of alginate remains challenging due to the resistance of its block structures to acid hydrolysis and the partial degradation of liberated uronic acids, these methods are sufficient for assessing purity and estimating M/G ratios [34–36].

This study introduces a two-step gelation method for sequentially isolating two structurally distinct low-molecular-weight alginic acid fractions from a hot-water extract of alcohol-insoluble residue (AIR) prepared from sugar kelp. Purity and molecular weight distribution were assessed using GC- and HPLC-based chromatographic techniques, followed by structural characterization using improved ^1^H NMR and ATR-FTIR approaches.

## 2. Materials and methods

### 2.1. Seaweed material

*Saccharina latissima* was harvested from the Pacific Northwest of Canada on May 1, 2022, by Cascadia Seaweed Corporation (Sidney, British Columbia, Canada). Following harvest, the seaweed was rinsed to remove debris and air-dried at temperatures not exceeding 30 °C for 96 h [10]. The dry samples were then shipped in sealed bags at ambient temperature to the Lethbridge Research and Development Centre for analysis.

### 2.2. Preparation of AIR

The dry seaweed sample was ball-milled into a fine powder using a Mixer Mill MM 400 system (Retsch, Germany). The powder was transferred into nineteen 50 mL tubes and weighed, with each tube containing 3.1–3.5 g of powder (Table S1). Subsequently, 40 mL of 95% (v/v) ethanol was added to each tube. The tubes were capped, vortexed, and incubated at 60 °C for 2 h; this treatment was performed twice. After each incubation, the samples were centrifuged (3000 × *g*, 30 min) at room temperature, and the pellets were collected for the subsequent incubation. A similar procedure was performed using 80% (v/v) ethanol at 40 °C for three cycles (8 h, 16 h, and 8 h), followed by two 30-min incubations with absolute ethanol at 60 °C. The final residues were vacuum-dried directly in the tubes without heating using a SpeedVac vacuum concentrator (Thermo Fisher, USA). The AIR preparation yields for each tube are shown in Table S1.

### 2.3. Extraction and fractionation of alginates

Sugar kelp AIR (32.287 g) was divided among sixteen 50 mL tubes, each containing approximately 2 g of sample. To each tube, 40 mL of deionized water was added, followed by vortexing and incubation at 70 °C for 24 h. This extraction was performed for three cycles. After each incubation, the mixture was centrifuged at 3000 × *g* for 30 min at room temperature to separate the residues for subsequent extraction cycles. The supernatants from all extractions were pooled, filtered through six layers of clean cheesecloth, and allowed to stand at 4 °C for 24 h. The pooled extract was then centrifuged at 3000 × *g* for 30 min at 4 °C, and the resulting supernatant was extensively dialyzed against deionized water with a molecular-weight cutoff (MWCO) of 14 kDa to facilitate the removal of low-molecular-weight components, including residual laminarin. The dialyzed solution was adjusted to a final volume of 2 L with deionized water, followed by the slow addition of 222 mL of acetic acid. The mixture was gently shaken at 30 °C for 24 h in an incubator. After centrifugation (3000 × *g*, 30 min, room temperature), the pellet was washed with a water–acetic acid mixture (9:1, v/v) and collected by centrifugation. The pellet was resuspended in deionized water, extensively dialyzed (MWCO 14 kDa) against deionized water, and freeze-dried to yield 1.990 g of material, designated SKP-G1. The supernatant was transferred to 50 mL tubes, frozen at −20 °C for 24 h, and then thawed at 4 °C for 24 h. This freeze–thaw cycle was repeated once, followed by centrifugation (3000 × *g*, 30 min). The resulting pellet was resuspended in deionized water, extensively dialyzed (MWCO 14 kDa) against deionized water, and freeze-dried to yield 0.315 g of material, designated SKP-G2. A flow diagram of the isolation process is shown in Scheme S1.

### 2.4. Chromatographic analysis

#### 2.4.1. Monosaccharide analysis by HPAEC-PAD

Approximately 2 mg of each fraction was stirred in 1 mL of 2 M trifluoroacetic acid (TFA) at 120 °C for 2 h [40]. After evaporation to dryness, the hydrolysates were co-evaporated to dryness twice with 1 mL of anhydrous methanol and subsequently redissolved in 25 mL of deionized water. An aliquot (1.5 mL) of each resulting solution was filtered through a 0.2 μm nylon syringe filter, and 10 μL was injected via an autosampler into a Dionex ICS-3000 HPAEC-PAD system (Thermo Scientific, USA) equipped with a CarboPac PA20 guard column (3 × 30 mm) and a CarboPac PA20 analytical column (3 × 150 mm). The column compartment was maintained at 15 °C, and the mobile phase, delivered at 0.3 mL/min, was composed of three aqueous solvents: 100 mM NaOH (Solvent A), 1 M sodium acetate in 100 mM NaOH (Solvent B), and water (Solvent C), using the following gradient: 20% A/0% B at 0 min; 19% A/1% B at 20 min; 15% A/5% B at 20.1 min; 80% A/20% B at 60 min; 100% A at 60.1 min (held until 70 min); and 20% A/0% B at 70.1 min (held until 80 min). PAD detection was performed using a gold-on-polytetrafluoroethylene disposable working electrode with the standard quadruple-potential waveform for carbohydrate analysis. Peak assignments were made by comparing retention times with those of monosaccharide standards. Data acquisition and analysis were performed using Chromeleon CDS software version 6.80 SR10 (Thermo Fisher, USA). Two separate experiments were conducted for each sample.

#### 2.4.2. Monosaccharide analysis by GC-MS and GC-FID

The analysis was performed following previous reports [41, 42], with modifications. Approximately 5 mg of each fraction was heated in 1 mL of 1.25 M methanolic HCl at 80 °C for 16 h under gentle magnetic stirring [37]. The reaction mixture was evaporated to dryness under N_2_, followed by two rounds of co-evaporation to dryness with 1 mL of anhydrous methanol. The resulting methyl esters of uronic acid methyl glycosides were then stirred in 1 mL of deionized water containing 10 mg/mL sodium borodeuteride (NaBD_4_) at 4 °C for 24 h to achieve carboxyl reduction and 6,6^′^-dideuteration [10, 43]. Excess reductant was quenched with glacial acetic acid, and the mixture was evaporated to dryness under N_2_. The residue was acetylated by heating in 0.5 mL of TFA and 2.5 mL of acetic anhydride (Ac_2_O) at 60 °C for 1 h [10]. The acetylated products were dissolved in 3 mL of dichloromethane (DCM) and partitioned once against 3 mL of saturated

NaHCO_3_ solution, followed by three washes with 3 mL of deionized water. The organic phase was passed through a glass wool-plugged Pasteur pipette loaded with anhydrous Na_2_SO_4_ powder and then evaporated to dryness under N_2_. Subsequent conversion of the acetylated methyl glycosides to alditol acetates involved hydrolysis with 2 M TFA at 120 °C for 2 h, reduction with NaBD_4_, and acetylation in a heated mixture of TFA and Ac_2_O (1:5, v/v) at 60 °C for 1 h [10]. The resulting derivatives were purified by partitioning and passage through an anhydrous Na_2_SO_4_ column, dissolved in ethyl acetate, and analyzed using an Agilent 7890A-5977B GC-MS system equipped with a Supelco SP-2380 column (60 m × 0.25 mm × 0.20 μm; Sigma-Aldrich). The oven temperature was programmed to start at 180 °C (1 min hold) and then increase at 2 °C/min to 250 °C (30 min hold). The same derivative preparation was also analyzed using an Agilent 7890A GC-FID system equipped with an identical column, with the oven temperature programmed to start at 55 °C (1 min hold), increase at 20 °C/min to 180 °C, and then increase at 2 °C/min to 250 °C (30 min hold). Both systems employed an inlet temperature of 250 °C and a constant helium flow rate of 0.8 mL/min; split injection (10:1) was used for GC-MS, whereas splitless injection was used for GC-FID. Derivatives were identified based on EI-MS fragmentation patterns and retention times compared with those of standards [44]. Quantitation was performed by FID, for which relative response factors of 0.745, 0.890, 0.835, and 0.535, calculated using the effective carbon number concept, were applied to the acetate derivatives of pentitol, hexitol, 6-deoxyhexitol, and 1,6-anhydrohexopyranose, respectively [45, 46]. Data were analyzed using Agilent OpenLab CDS software version 2.6. Two separate experiments were conducted for each sample.

#### 2.4.3. Glycosidic linkage analysis by GC-MS

The uronic acids in approximately 5 mg of each fraction were carboxyl-reduced to 6,6^′^-dideuterated neutral sugars via NaBD_4_ reduction of their methyl esters generated by mild methanolysis (0.5 M methanolic HCl, 80 °C, 20 min) [10, 47–49]. Excess reductant was quenched with glacial acetic acid, and borate was removed by repeated evaporation to dryness, first with a mixture of acetic acid and methanol (1:9, v/v) and then with anhydrous methanol. The samples were subsequently methylated with 0.6 mL of methyl iodide in 1 mL of DMSO in the presence of 100 mg of dry NaOH powder [10]. The methylation products were suspended in 3 mL of DCM and partitioned twice against 6 mL of a water–acetic acid mixture (9:1, v/v), followed by three washes with 6 mL of deionized water. The lower phase was evaporated to dryness, and the methylation was repeated once. The methylated samples were then converted to partially methylated alditol acetates (PMAAs) via hydrolysis with 2 M TFA at 120 °C for 2 h, reduction with NaBD_4_, and acetylation in a mixture of TFA and Ac_2_O (1:5, v/v) at 60 °C for 1 h [10]. The resulting PMAAs were dissolved in ethyl acetate and analyzed using an Agilent 7890A-5977B GC-MS system equipped with a Supelco SP-2380 column (60 m × 0.25 mm × 0.20 μm; Sigma-Aldrich). The oven temperature was programmed to start at 175 °C (1 min hold), increase at 1 °C/min to 215 °C (10 min hold), and then increase at 1.5 °C/min to 250 °C (20 min hold). EI-MS spectra of the PMAAs were interpreted by comparison with reference spectra and literature data [50]. Relative molar compositions (mol%) of the PMAAs were estimated by dividing the TIC peak area of each PMAA by its molecular mass, in accordance with the published protocol [44]. Data were analyzed using Agilent OpenLab CDS software version 2.6. Two separate experiments were conducted for each sample.

#### 2.4.4. HPSEC analysis

Each fraction (~2 mg) was dissolved in 2 mL of 0.1 M sodium phosphate buffer (pH 7.0) by magnetic stirring overnight. The resulting solution was filtered through a 0.2 μm nylon syringe filter and transferred to a 2 mL LC vial. An aliquot (100 μL) was automatically injected into an HPSEC system comprising an Agilent 1260 Infinity II instrument equipped with a G7129A vialsampler, a G7110B isocratic pump, and a G7116A multicolumn thermostat (MCT) containing, in sequence, a PL Aquagel-OH guard column (7.5 × 50 mm, 8 μm), two PL Aquagel-OH mixed-H columns (7.5 × 300 mm, 8 μm), and a Shodex OH-pak SB-806M HQ column (300 × 8 mm, 13 μm). The HPSEC system was coupled in series to an Agilent WR G7115A diode array detector (DAD), a Wyatt DAWN 18-angle static light scattering (SLS) detector, commonly referred to as a multi-angle light scattering (MALS) detector, with an integrated quasi-elastic light scattering (QELS) detector at 90° for dynamic light scattering (DLS), a Wyatt ViscoStar capillary-bridge differential viscometer (VIS), and a Wyatt Optilab differential refractive index (RI) detector. The mobile phase consisted of 0.1 M NaNO_3_ containing 0.03% (w/v) NaN_3_ and was delivered at a constant flow rate of 0.5 mL/min. The MCT module and the DAD, VIS, and RI detectors were maintained at 40 °C. Data acquisition and analysis were performed using ASTRA software version 8 (Wyatt Corporation, USA). A specific refractive index increment (dn/dc) of 0.165 mL/g was used for data analysis [51, 52]. Two separate experiments were conducted for each sample.

### 2.5. Solution-state ^1^H NMR analysis

#### 2.5.1. NMR sample preparation and spectrum collection

Each fraction (~20 mg) was dissolved in 4 mL of 0.1 M sodium phosphate buffer (pH 7.0) by magnetic stirring overnight, followed by extensive dialysis against deionized water (MWCO 14 kDa) and subsequent freeze-drying. The resulting sodium alginate was subjected to three cycles of redissolution in 1 mL of D_2_O (99.9% atom D, product no. 151882, Sigma-Aldrich) and freeze-drying. After the final freeze-drying step, the sample was redissolved in 1 mL of high-purity D_2_O (≥99.96% atom D, product no. 151890, Sigma-Aldrich), followed by the addition of 30 µL of D_2_O containing 0.75% (w/w) sodium 3-trimethylsilyl-2,2,3,3-tetradeuteriopropionate (TSP-d_4_; 99.9% atom D, product no. 293040, Sigma-Aldrich). Approximately 0.7 mL of this solution was transferred into a 5-mm Wilmad precision NMR tube (product no. Z412007, Sigma-Aldrich), and two NMR samples were prepared for each alginate fraction. After ^1^H NMR data collection, the D_2_O solution was recovered from each tube, filtered through a Pasteur pipette plugged with glass wool, and transferred into another 5-mm NMR tube for subsequent ^1^H NMR analysis. NMR spectra of the unfiltered samples were collected at 65 °C and 80 °C, whereas spectra of the glass wool-filtered samples were acquired only at 80 °C. All experiments were performed on a Bruker Avance III HD 700 MHz NMR spectrometer (Bruker Corporation, Germany) operating at 700.44 MHz and equipped with a triple-resonance TBO-Z probe with the outer coil tuned to ^1^H. 1D ^1^H NMR spectra were recorded using the standard zg30 pulse sequence with a 1 s relaxation delay, a 6.4360 s acquisition time, 1800 scans, a spectral size of 65,536, and a spectral width of 10 ppm. The raw 1D data were processed without apodization or zero-filling prior to Fourier transformation and were then phase-adjusted and baseline-corrected using Mnova software version 14.2.1 (build 27684; Mestrelab Research S.L., Santiago de Compostela, Spain). 2D ^1^H–^1^H COSY and TOCSY spectra were acquired using the cosygpmfqf and dipsi2gpphzs pulse sequences, respectively, with 80 scans per t_1_ increment, 2048 points in the t_2_ dimension, and 256 points in the t_1_ dimension. A mixing time of 80 ms was used for TOCSY. The acquisition times were 0.2925 s for COSY and 0.2922 s for TOCSY. All 2D spectra were acquired with a spectral width of 10 ppm in both dimensions. Prior to Fourier transformation, the raw 2D data were apodized with sine-square functions in Mnova software using the following optimized settings: For COSY, 90° was applied in the t_1_ dimension for SKP-G1 and 0° for SKP-G2, with 0° applied in t_2_ for both fractions. For TOCSY, 90° was applied to both t_1_ and t_2_ for both fractions. For all apodizations, the first point was set to 0.5 in t_1_, with no first-point setting applied in t_2_. This was followed by zero-filling and linear prediction in the t_1_ dimension using modified iterative soft thresholding (MIST) to extend the data to 1024 points; no zero-filling or linear prediction was applied in the t_2_ dimension. Baseline correction was performed using the “Full Auto (Bernstein Polynomials)” setting, and the spectra were symmetrized using the “COSY-like” function in Mnova software.

#### 2.5.2. Determination of frequency values of monads, diads, and G-centered triads

A modified version of the previously reported methods [26–28, 53], optimized for processing 700 MHz 1D ^1^H NMR spectra of alginates, was applied. The quantitative Global Spectral Deconvolution (qGSD) integration tool in Mnova software was used to integrate four well-resolved peaks in the anomeric region of the assigned 1D ^1^H NMR spectra of SKP-G1 and SKP-G2, including H-5 of the central G residue in the G<u>G</u>M triad (designated 5G<u>G</u>M), H-5 of the central G residue in the M<u>G</u>M triad (designated 5M<u>G</u>M), H-1 of the underlined M residue in the <u>M</u>M diad (designated 1<u>M</u>M), and H-5 of the central G residue in the G<u>G</u>G triad (designated 5G<u>G</u>G), with the corresponding integrals designated as I_1_, I_2_, I_3_, and I_4_, respectively. The frequencies of monads, diads, and G-centered triads were then calculated using the following equations:

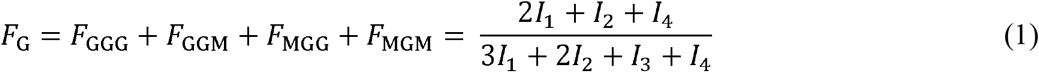

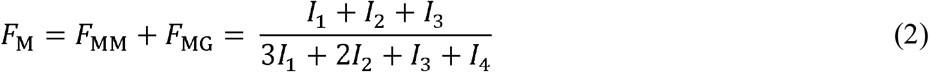

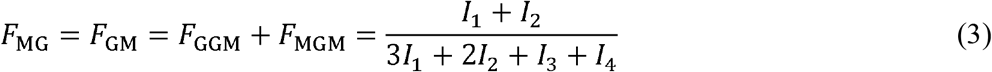

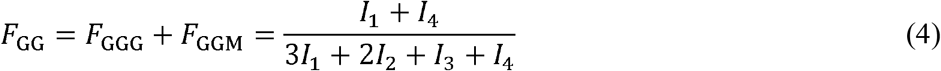

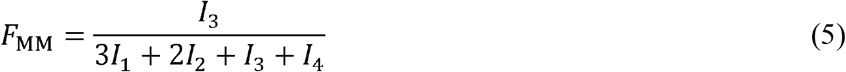

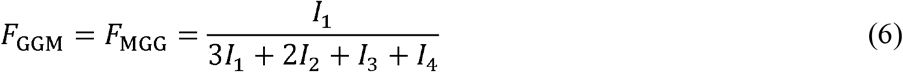

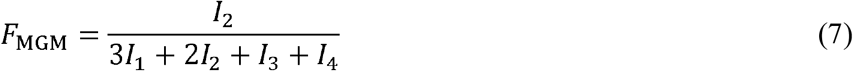

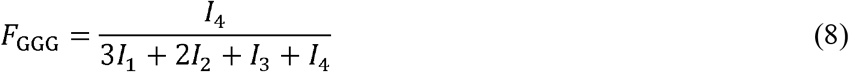

#### 2.5.3. Determination of frequency ranges of M-centered triads

Determining the exact values of the M-centered triad frequencies requires at least one peak integral corresponding to an M-centered triad, in addition to the peak integrals of all diads; however, this is not achievable using 1D ^1^H NMR because of signal overlap [24, 26], even at the high field of 700 MHz used in the present study. Nevertheless, as demonstrated in our previous study [1], once the diad frequencies are fixed, the frequencies of the corresponding triads cannot exceed those of their constituent diads and are further constrained within defined ranges by the inherent hierarchical relationships between the two sequence levels. These constraints can therefore be quantitatively expressed by the three inequality relations below, which were rigorously proved by mathematical induction [1]. These inequality relations enable calculation of the constrained frequency range of each M-centered triad based on the values of *F*_MM_ and *F*_MG_ obtained in Section 2.5.2.

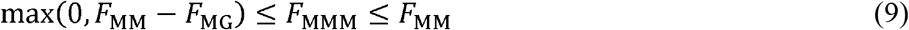

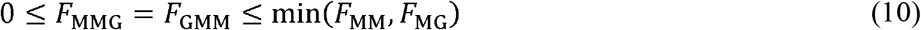

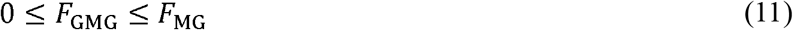

#### 2.5.4. Determination of frequency ranges for homo-G-blocks and homo-M-blocks

Based on the known frequency values of G-centered triads (GGG and GGM), the frequency range of homo-G-blocks, denoted as (G)_*m*+3_, where *m* + 3 represents the degree of polymerization (DP) of the homoblock and *m* represents the difference in DP between the homoblock and the triad, can be calculated using the following inequality relation established in our previous study [1]:

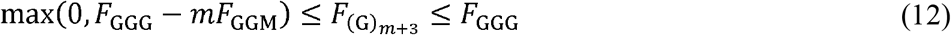

Based on the known frequency values of the MM and MG diads, the frequency range of homo-M-blocks, denoted as (M)_*m*+2_, where *m*+2 represents the DP of the homoblock and *m* represents the difference in DP between the homoblock and the diad, can be calculated as follows:

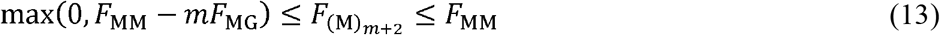

Using inequality relation (12), the frequency ranges of homo-G-blocks in the G-rich SKP-G1 were calculated based on *F*_GGG_ and *F*_GGM_ determined in Section 2.5.2. Similarly, using inequality relation (13), the frequency ranges of homo-M-blocks in the M-rich SKP-G2 were calculated based on *F*_MM_ and *F*_MG_ determined in Section 2.5.2. For demonstration and graphical visualization, the frequency ranges of both types of homoblocks were calculated up to a DP of 18.

#### 2.5.5. Evaluation of chain-growth statistical models using ^1^H NMR-derived frequency ranges of homoblocks

The probability of elongating a homoblock (G)_*n*_ to (G)_*n*+1_ by adding an additional G residue corresponds to *F*_G_ in a Bernoulli chain, *F*_GG_/*F*_G_ in a first-order Markov chain, and *F*_GGG_/*F*_GG_ in a second-order Markov chain [1, 27, 28]. Accordingly, the probability of forming a homoblock (G)_*n*_, denoted as 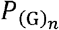, is calculated using the following equations for the Bernoulli process, first-order Markov chain, and second-order Markov chain, respectively [1]:

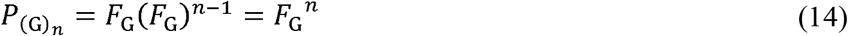

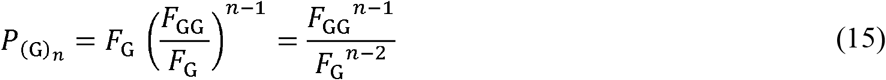

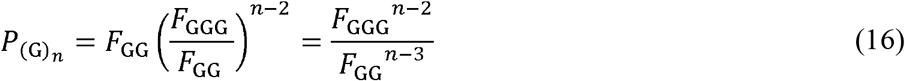

Similarly, for a homoblock (M)_*n*_, the probability 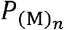 is given by the equations below for the Bernoulli process, first-order Markov chain, and second-order Markov chain, respectively:

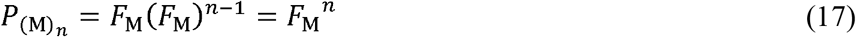

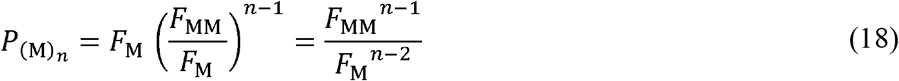

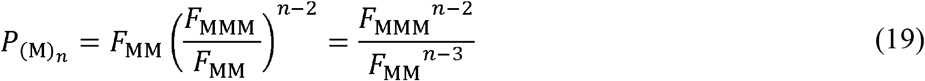

In equation (19), *F*_MM_ is required to calculate 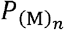 under the second-order Markov chain model, whereas *F*_MMM_, as determined in Section 2.5.3, is represented by a range rather than a single value, with the range constrained by its constituent diad frequencies, *F*_MM_ and *F*_MG_, according to equation (9), and therefore cannot be directly applied for this purpose. Accordingly, the lower limit of this range, designated *F*_MMM,min_,was used to calculate the minimum possible 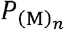, designated 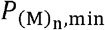, using the following equation:

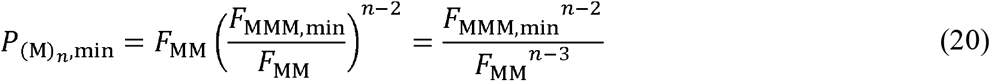

Using these equations, the frequency values of homoblocks (G)_*n*_ in SKP-G1 and (M)_*n*_ in SKP-G2 were predicted for the three chain-growth statistical models. The predicted frequencies were evaluated against the experimentally determined frequency ranges of homoblocks (Section 2.5.4). A model was deemed invalid if any predicted homoblock frequency, at any DP, fell outside the corresponding experimental range [1]. Conversely, if the predicted frequencies at all selected DPs fell within their respective experimental ranges, the model could not be ruled out based on the available data.

### 2.6. ATR-FTIR analysis

#### 2.6.1. Acquisition of ATR-FTIR spectra and their first- and second-derivative analyses

ATR-FTIR analysis of SKP-G1 and SKP-G2 was performed using a JASCO FT/IR-4X spectrometer (JASCO Corporation, Japan) equipped with a diamond ATR prism. Spectra were recorded in absorbance mode over a wavenumber range of 400–4000 cm^−1^ at a resolution of 4 cm^−1^. Data for each sample were collected from 128 co-added scans using Spectra Manager software version 2.5 (JASCO Corporation, Japan). First- and second-derivative analyses of the raw spectra were performed using OriginPro software version 2023 (OriginLab Corporation, USA), with Savitzky-Golay smoothing applied using 5–19 smoothing points and an optimized polynomial order of 2. Three separate experiments were conducted for each sample.

#### 2.6.2. Logarithmic cosine transformation (LCT) of derivative ATR-FTIR spectra

The logarithmic cosine transformation (LCT) algorithm, previously developed in our recent report for post-processing 2D HSQC NMR data [2], was adapted here for processing first- and second-derivative ATR-FTIR spectra, as detailed below.

For each derivative value, *D*, at a given wavenumber in a first- or second-derivative ATR-FTIR spectrum, the corresponding LCT-transformed value, *Z*, was calculated as:

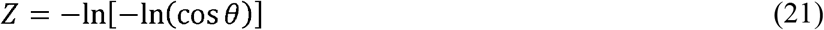

where *θ* is an auxiliary angular variable within the range of −π/2 to π/2 and is defined as:

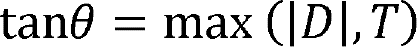

or, equivalently,

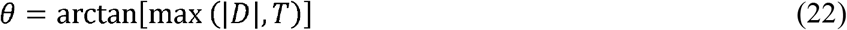

Here, |*D*| denotes the absolute value of the derivative value *D*, and *T* is a selected threshold represented by a very small positive value close to, but greater than, zero. The threshold *T* is introduced to prevent tan*θ* from becoming zero and cos*θ* from becoming 1 when *D* = 0, which would cause *Z* in equation (21) to become undefined (+∞) [2]. In addition, *T* establishes a finite maximum value of *Z* when |*D*| ≤ *T*.

Substitution of equation (22) into equation (21) gives:

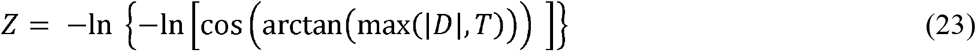

Rewriting equation (23) in an equivalent non-trigonometric form gives:

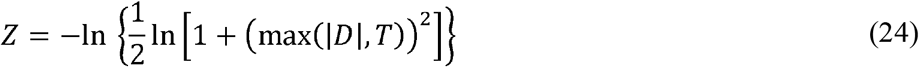

*T* was set to 1.1 × 10^−8^, following our previous report [2]. This threshold prevents very small derivative values from being rounded to zero due to Microsoft Excel’s numerical precision limitations, thereby avoiding undefined logarithmic calculations and establishing a maximum *Z* value of 36.7368005696771 [2]. Accordingly, equation (24) was implemented as:

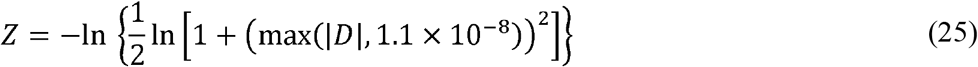

LCT-transformed spectra were generated from the first- and second-derivative spectra of SKP-G1 and SKP-G2 by plotting the *Z* values, calculated from the derivative values using equation (25), in Excel 2024 (Microsoft, USA). A proof-of-concept PCA of the original derivative spectra and the corresponding LCT-transformed derivative spectra of the two fractions was performed using R software with a script modified from our previous report [10] and provided in the supplementary data.

## 3. Results and discussion

### 3.1. Isolation, purity, and molecular weight distribution

#### 3.1.1. Extraction and fractionation

Alginate has been detected in hot-water extracts of various brown seaweeds [54–56]. Hot-water extraction is a green alternative to conventional alkaline extraction. A previous analysis of 16 brown seaweeds revealed that, although hot-water extraction produced a lower overall yield, it generated extracts with higher M levels than alkaline extraction [54]. Hot-water extraction has been employed to isolate alginates and fucoidans from the AIR of sugar kelp, with alginate subsequently precipitated by the addition of CaCl_2_ [16]. In this study, AIR samples were prepared from ball-milled sugar kelp powders with good yields (Table S1). The AIR was extensively extracted with hot water, followed by a two-step gelation process to sequentially isolate two alginic acid fractions from the water extract acidified with acetic acid: first, gelation was induced by mixing the extract with acetic acid (9:1, v/v) to yield SKP-G1; subsequently, two cycles of freezing at −20 °C and thawing at 4 °C produced SKP-G2, as detailed in Section 2.3 and Scheme S1. It was apparent that acidification alone triggered gelation in some alginates, whereas others required additional freeze–thaw cycles to form a gel. This phenomenon could be explained as follows: during freezing, ice formation concentrates both alginate and acetic acid in the remaining solution [57]. The increased alginate concentration promotes closer packing of alginate chains, while the elevated acetic acid concentration lowers the pH, protonating carboxylate groups (−COO^-^) to form carboxylic acids (−COOH) [57]. This conversion reduces electrostatic repulsion and enhances hydrogen bonding between closely packed chains, thereby favoring gel formation [57]. It is well established that acid-induced formation of stable gels is favored in alginates rich in homo-G-blocks compared with those rich in homo-M-blocks [58, 59]. Differences in monosaccharide composition and homoblock types between SKP-G1 and SKP-G2 were examined using solution-state 700 MHz ^1^H NMR experiments (Section 3.2). As part of the sample quality assessment prior to NMR analysis, the chromatographic studies described below were conducted to confirm that the purity and molecular weights of SKP-G1 and SKP-G2 met the requirements for high-resolution ^1^H NMR analysis.

#### 3.1.2. Purity assessment based on monosaccharide and glycosidic linkage analysis

Acid-catalyzed depolymerization, via acid methanolysis or aqueous acid hydrolysis, is widely employed to release uronic acids from alginate for monosaccharide composition analysis [34–36]. However, the strong glycosidic bonds in alginate necessitate harsh conditions that, while promoting depolymerization, also degrade the liberated uronic acids [13, 34–36]. Hydrolysis efficiency depends on both monosaccharide composition and glycosidic linkage; for example, G residues are more readily released than M residues [13, 34–36], and G–M linkages cleave more easily than M–G and M–M linkages [60], leading to uneven release of G and M residues during hydrolysis. Moreover, once liberated, G and M degrade at different rates, further complicating quantitation of the constituent monosaccharides [13, 61]. Despite extensive optimization, both acid methanolysis and aqueous acid hydrolysis remain unreliable for absolute quantitation due to non-synchronous release, degradation, and low and uneven recovery, although they are acceptable for estimating M/G ratios in similarly treated samples for comparative purposes [13, 34–36]. In this study, HPAEC-PAD analysis of SKP-G1 and SKP-G2 following 2 M TFA hydrolysis showed major peaks corresponding to the alginate-derived uronic acids, together with a minor fucose peak (Fig. S1). Although hydrolysis with 2 M TFA (120 °C, 2 h) enabled quantitative fucose release for evaluating minor fucoidan contamination [40, 62], it may result in low recovery of M and G residues from alginate [63]. A previous report showed that, even under optimized conditions, 2 M TFA hydrolysis achieved only ~29% total recovery of uronic acids, with recoveries of 24% for G and 5% for M residues [36], suggesting that the relative fucose peak areas (2.7% for SKP-G1 and 14.3% for SKP-G2) were substantially overestimated (Table S2), with the larger relative fucose peak area in SKP-G2 likely due in part to the lower recovery of M residues during hydrolysis. Consequently, further quantitation based on these HPAEC-PAD data was not pursued.

Carboxyl reduction has been employed to convert the acid-resistant polyuronate chains of alginate into the corresponding neutral sugar chains, thereby facilitating subsequent acid hydrolysis [38, 64]. The resulting gulomannan or mannogulocan depolymerizes more readily during acid hydrolysis, with liberated mannose remaining relatively stable under the hydrolysis conditions, while some of the liberated gulose is converted to 1,6-anhydro-β-L-gulopyranose (hereafter referred to as 1,6-anhydrogulose), yielding a mixture of these three products [38, 65–67]. This conversion is accounted for in the quantitative analysis: the constituent M and G residues in the original alginate are determined from the amount of mannose and the combined amounts of gulose and 1,6-anhydrogulose, respectively, as measured by GC or HPLC, allowing calculation of the M/G ratio [38, 65, 66]. Alternatively, carboxyl-reduced alginic acid can be subjected to methylation-GC-MS to determine glycosidic linkages [64]. In this approach, 1,6-anhydrogulose is not generated during hydrolysis of the permethylated gulomannan or mannogulocan because methylation at *O*-6 prevents formation of the 1,6-anhydro bridge required for its production. In the present study, the uronic acid residues in SKP-G1 and SKP-G2 were depolymerized using 1.25 M methanolic HCl at 80 °C for 16 h, followed by carboxyl reduction of the resulting uronic acid methyl esters with NaBD_4_ to yield the corresponding 6,6^′^-dideuterated neutral sugars. These products were subsequently converted through a multistep derivatization procedure into mixtures of alditol acetates and 1,6-anhydrogulose acetates, as detailed in Section 2.4.2. Peaks in the GC-MS chromatograms (Fig. S2) were identified based on retention times and EI-MS spectra (Figs. S3–S6), and the corresponding compounds were quantified using GC-FID responses (Fig. S7; Table S3). The results showed that SKP-G1 contained 94.2% combined M and G residues and 1.24% fucose, whereas SKP-G2 contained 83.3% combined M and G residues and 7.48% fucose, with only minor amounts of other monosaccharides detected in both samples (Table S3). Bojorges et al. (2023) reported that the use of strong acid methanolysis for depolymerization in monosaccharide analysis of high-M-block, low-molecular-weight alginates could lead to substantial underestimation of M residues compared with values obtained by ^1^H NMR [34]. This underestimation of M in SKP-G2 was further supported by linkage analysis (Figs. S8–S10), which revealed a higher proportion of 4-linked mannopyranuronic acid residues (4-Man*p*A) than of 4-linked gulopyranuronic acid residues (4-Gul*p*A) in SKP-G2 (Table S4). Although the relative proportion of M residues determined by linkage analysis remained lower than that determined by ^1^H NMR (Section 3.2.1), it was closer to the value obtained by ^1^H NMR than to that obtained from the GC-FID-based monosaccharide analysis described above. It is worth noting that the linkage analysis in this study focused on the predominant alginate components rather than the minor fucoidan contaminants in the fractions. Accordingly, the aqueous phase obtained by DCM–water partitioning of the methylation reaction mixture, which contained the methylated fucoidan components, was not collected for linkage analysis, and fucoidan-derived backbone fucose linkages were therefore not detected (Table S4). Nevertheless, minor amounts of terminal fucopyranose (*t*-Fucp) were detected in both fractions, possibly arising from fucose that was released from minor fucoidans and desulfated during the mild methanolysis step used to esterify the uronic acid residues of alginate prior to NaBD_4_ treatment for carboxyl reduction.

These results supported previous reports that both methanolysis- and TFA hydrolysis-based monosaccharide analyses could lead to an underestimation of uronic acid residues, especially M residues in high-M alginate samples, although these methods remain valuable as rapid approaches for comparative analysis of similarly treated samples and for purity estimation prior to ^1^H NMR analysis [34, 35]. It was apparent that these depolymerization-based monosaccharide analyses overestimated the relative proportion of fucose in the fractions, particularly in the M-rich SKP-G2. However, no characteristic fucose signals were observed in the 1D and 2D ^1^H NMR spectra (Sections 3.2.1 and 3.2.2), suggesting that residual fucoidan did not substantially interfere with the NMR analysis.

#### 3.1.3. HPSEC analysis

Both SKP-G1 and SKP-G2 readily dissolved in 0.1 M sodium phosphate buffer (pH 7.0), allowing for HPSEC analysis (Figs. S11–S13, Table S5). Each alginate fraction exhibited a single, broad peak in the RI chromatogram, and the DAD chromatogram at 280 nm showed a broad, low-intensity UV signal synchronous with the corresponding RI peak (Figs. S11 and S12), suggesting that the UV absorbance was associated with the polydisperse alginate fractions rather than a separately eluting protein component. The absence of prominent amide bands in the ATR-FTIR spectra (Section 3.3.1) further suggested a lack of substantial protein contamination.

The molecular weight distributions were determined by HPSEC using a MALS detector to measure static light scattering and an RI detector to measure concentration. The resulting weight-average (M_w_) and number-average (M_n_) molecular weights were 40.8 and 22.3 kDa, respectively, for SKP-G1, and 38.5 and 23.8 kDa, respectively, for SKP-G2 (Table S5). Based on these molecular weight values, both fractions were classified as low-molecular-weight alginates. According to the literature [68], alginates produced by fractionation or selective processing typically exhibit a polydispersity index (M_w_/M_n_) below 2. The resulting polydispersity indices were 1.8 for SKP-G1 and 1.6 for SKP-G2 (Table S5).

Using a differential viscometer, the number-average, weight-average, and z-average intrinsic viscosities were determined to be 101.2, 179.9, and 368.2 mL/g, respectively, for SKP-G1, and 68.8, 106.8, and 211.2 mL/g, respectively, for SKP-G2 (Table S5). These relatively low intrinsic viscosities indicated that the alginate chains occupied relatively small hydrodynamic volumes in solution, consistent with their low molecular weights. The power-law exponent α of the Mark-Houwink-Sakurada (MHS) equation was 0.975 for SKP-G1 and 0.937 for SKP-G2 (Table S5), within the typical range of 0.91–1.13 for alginates in a solution with an ionic strength of 0.1 M [68]. In general, α values in the ranges of 0–0.5, 0.5–0.8, and 0.8–1.8 correspond to spherical, random-coil, and rod-like conformations in solution, respectively [68]. Thus, the α values of both fractions were consistent with relatively extended, rod-like chain conformations in solution. Alginate chain stiffness increases in the order of MG alternating blocks < homo-M-blocks < homo-G-blocks [69]. The higher α value for SKP-G1 was consistent with the ^1^H NMR data, which indicated that this fraction was G-block-rich, whereas SKP-G2 contained more M blocks, as detailed in Section 3.2.1.

Small hydrodynamic radius (R_h_) values of 16.4 nm for SKP-G1 and 12.9 nm for SKP-G2 were determined by differential viscometry (Table S5). In addition, using a right-angle DLS detector, R_h_ was independently measured (Table S5), complementing the R_h_ values obtained from differential viscometry. The radius of gyration (R_g_), or root-mean-square radius, was determined from the angular dependence of scattered light by MALS. As expected for low-molecular-weight alginates such as SKP-G1 and SKP-G2, the R_g_ and R_h_ values measured by light scattering exhibited low precision (Table S5) because of the weak scattering intensity associated with their low molecular weights [70]. Although the accuracy of these size measurements could be improved by increasing the sample concentration to enhance light scattering intensity, such experiments were not pursued because molecular size determination was not a primary objective of the present study.

### 3.2. ^1^H NMR analysis

SKP-G1 and SKP-G2 readily dissolved in 100□mM sodium phosphate buffer (pH 7.0) and, after dialysis against deionized water followed by repeated redissolution in D_2_O and freeze-drying to achieve extensive deuterium exchange of labile protons, were redissolved in D_2_O for high-field ^1^H NMR analysis at 700 MHz. The resulting high-quality 1D and 2D ^1^H NMR spectra, collected at 65 °C and 80 °C, exhibited minimal variation, with only very minor peak displacements aside from the expected pronounced water peak shift. No significant differences were observed between spectra acquired at 80 °C from the D_2_O solution with and without glass wool filtration. Compared with previous studies using lower-field NMR (*e.g*., 300–500 MHz) [26, 28–32], the spectra obtained in this work demonstrated higher resolution and better-separated signals, attributable to the 700 MHz field strength and the lower molecular weights of these fractions. Although partial depolymerization via mild acid hydrolysis (*e.g*., 30 min at 100 °C, pH 3) is conventionally employed to reduce viscosity for high-molecular-weight alginate [53], SKP-G1 and SKP-G2 yielded excellent ^1^H NMR spectra without such mild acid pretreatment, thereby preserving their native structural integrity. Moreover, triethylenetetramine-N,N,N^′^,N^′′^,N^′′′^,N^′′′^-hexaacetic acid (TTHA) or ethylenediaminetetraacetic acid (EDTA) can be added to chelate calcium and prevent its binding to alginate G-blocks [53, 71], but this step is not universally necessary [21, 25]. In the present study, we omitted this step because both fractions dissolved completely in sodium phosphate buffer without forming a visible calcium phosphate precipitate, suggesting that residual Ca^2+^ ions did not substantially impair sample dissolution. After dialysis, redissolution in D_2_O produced clear NMR solutions with uniformly high spectral quality, even for the high-G-block fraction, regardless of filtration.

#### 3.2.1. Analysis of monad, diad, and triad frequencies via 1D ^1^H NMR

Signals in the anomeric region of the 1D ^1^H NMR spectra of both fractions (Fig. 1) were clearly resolved and assigned according to the literature [26], allowing unambiguous identification of G-centered triads (GGG, GGM, MGG, and MGM) and the MM and MG diads, thereby enabling determination of the frequencies of monads (*F*_M_ and *F*_G_), diads (*F*_MM_, *F*_MG_, *F*_GM_, and *F*_GG_), and G-centered triads (*F*_GGG_, *F*_MGM_, *F*_GGM_, and *F*_MGG_) (Tables 1 and S6). Although M-centered triads cannot be individually resolved by ^1^H NMR because of overlapping signals, their frequencies (*F*_MMM_, *F*_GMG_, *F*_MMG_, and *F*_GMM_) are confined within ranges defined by the diad frequencies *F*_MM_ and *F*_MG_, as delineated by the inequality relations proven through rigorous mathematical induction in our previous work [1]. As shown in Table 1, the results revealed similar frequencies for monads, diads, and G-centered triads, as well as comparable frequency ranges for M-centered triads (expressed as intervals), calculated from spectra collected at 65 °C without filtration (designated NF65), at 80 °C without filtration (designated NF80), and at 80 °C with glass wool filtration (designated GW80). The results indicated that SKP-G1 was rich in G blocks, whereas SKP-G2 was predominantly composed of M blocks. For example, under the GW80 condition, *F*_GGG_ was 0.52 for SKP-G1, whereas *F*_MMM_ ranged from 0.52 to 0.62 for SKP-G2 (Table 1).

**Table 1.** Results of ^1^H NMR-based monad, diad, and triad frequency analysis of SKP-G1 and SKP-G2.

| Values | SKP-G1 |  |  | SKP-G2 |  |  |
| --- | --- | --- | --- | --- | --- | --- |
|  | NF65 | NF80 | GW80 | NF65 | NF80 | GW80 |
| $F_G$ | 0.75 | 0.75 | 0.73 | 0.31 | 0.29 | 0.29 |
| $F_M$ | 0.25 | 0.25 | 0.27 | 0.69 | 0.71 | 0.71 |
| $F_{GG}$ | 0.60 | 0.60 | 0.57 | 0.21 | 0.19 | 0.19 |
| $F_{GM}$ | 0.15 | 0.15 | 0.16 | 0.10 | 0.09 | 0.10 |
| $F_{MG}$ | 0.15 | 0.15 | 0.16 | 0.10 | 0.09 | 0.10 |
| $F_{MM}$ | 0.11 | 0.10 | 0.11 | 0.60 | 0.62 | 0.62 |
| $F_{GGG}$ | 0.55 | 0.56 | 0.52 | 0.21 | 0.19 | 0.19 |
| $F_{GGM}$ | 0.06 | 0.05 | 0.05 | 0.00 | 0.00 | 0.00 |
| $F_{MGG}$ | 0.06 | 0.05 | 0.05 | 0.00 | 0.00 | 0.00 |
| $F_{MGM}$ | 0.09 | 0.10 | 0.11 | 0.10 | 0.09 | 0.10 |
| $F_{MMM}$ | [0.00, 0.11] | [0.00, 0.10] | [0.00, 0.11] | [0.50, 0.60] | [0.52, 0.62] | [0.52, 0.62] |
| $F_{MMG}$ | [0.00, 0.11] | [0.00, 0.10] | [0.00, 0.11] | [0.00, 0.10] | [0.00, 0.09] | [0.00, 0.10] |
| $F_{GMM}$ | [0.00, 0.11] | [0.00, 0.10] | [0.00, 0.11] | [0.00, 0.10] | [0.00, 0.09] | [0.00, 0.10] |
| $F_{GMG}$ | [0.00, 0.15] | [0.00, 0.15] | [0.00, 0.16] | [0.00, 0.10] | [0.00, 0.09] | [0.00, 0.10] |
| M/G | 0.34 | 0.33 | 0.37 | 2.28 | 2.50 | 2.50 |
| $\eta$ | 0.77 | 0.80 | 0.82 | 0.45 | 0.46 | 0.47 |

**Note:** Values are means from two separate experiments conducted for each sample. [a, b] denotes a closed interval on the real number line, including all values $x$ such that $a \leq x \leq b$ . The M-to-G ratio (M/G) is calculated as $F_M/F_G$ . Parameter $\eta$ is calculated as $F_{GM}/(F_G \times F_M)$ [53]. “NF65”, “NF80”, and “GW80” denote spectra collected at 65 °C without filtration, at 80 °C without filtration, and at 80 °C with glass wool filtration, respectively.

**Fig. 1.**
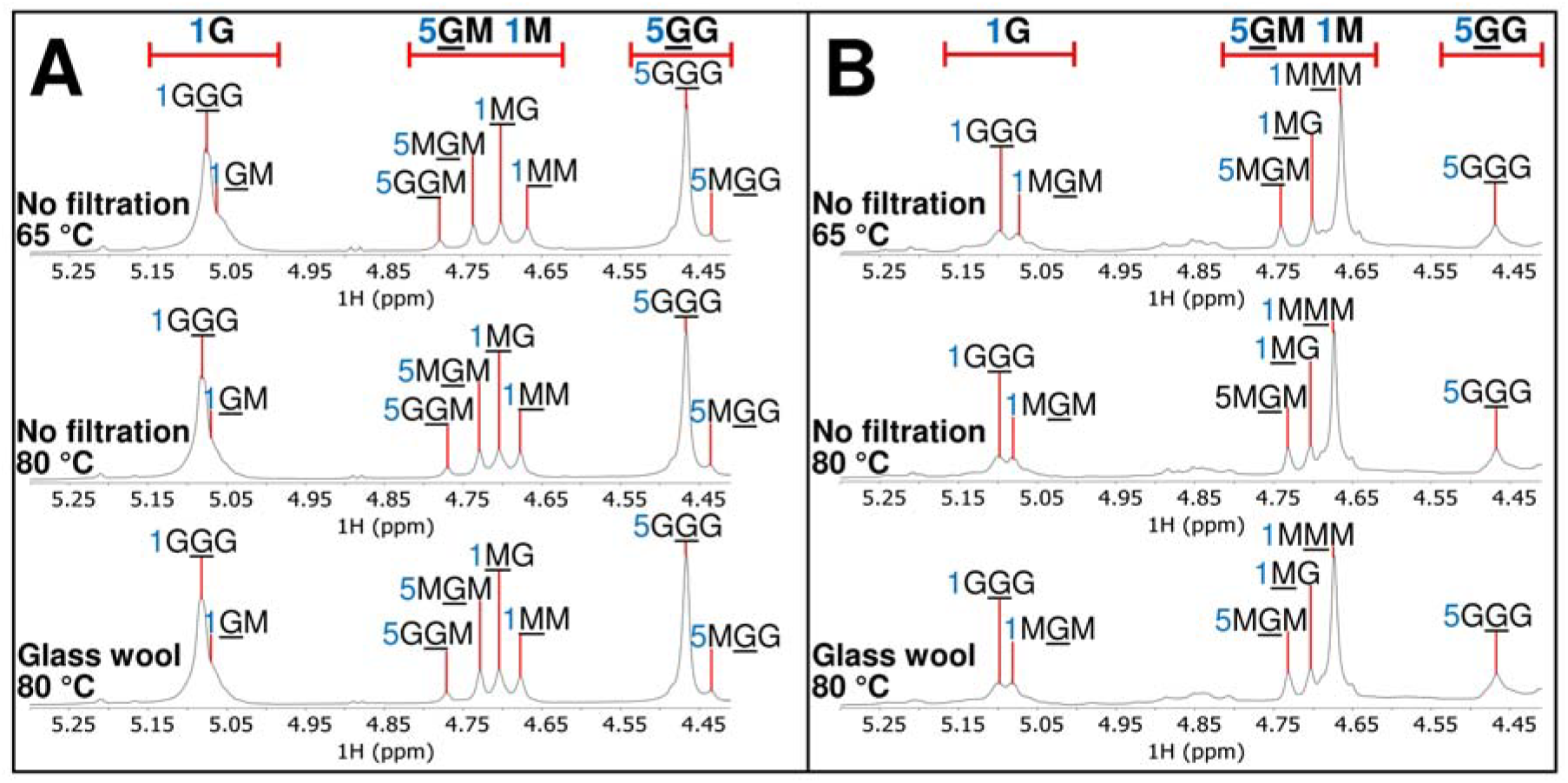
Assignment of peaks in the anomeric regions of the 1D ^1^H NMR spectra of (A) SKP-G1 and (B) SKP-G2 in D_2_O, collected at 65 °C without filtration (upper), at 80 °C without filtration (middle), and at 80 °C with glass wool filtration (lower). For example, the annotation “1G<u>G</u>G” denotes H-1 of the underlined central G residue in the G<u>G</u>G triad, whereas “1<u>M</u>G” indicates H-1 of the underlined M residue in the <u>M</u>G diad.

This strategy for calculating frequency ranges enabled us to assess the relative contributions of the H-1 signals from the underlined M residues in the M<u>M</u>M and G<u>M</u>M triads (designated 1M<u>M</u>M and 1G<u>M</u>M, respectively), both of which are unresolved in conventional 1D ^1^H NMR, to the H-1 signal of their merged <u>M</u>M diad (designated 1<u>M</u>M). In SKP-G2, *F*_MMM_ ranged from 0.52 to 0.62, whereas *F*_GMM_ ranged from 0.00 to 0.10 (Table 1). Notably, the minimum *F*_MMM_ was more than five times the maximum *F*_GMM_, indicating that 1M<u>M</u>M contributed at least five-sixths of the peak area of 1<u>M</u>M, with 1G<u>M</u>M accounting for at most one-sixth. Based on this substantial difference, it was reasonable to assign the sharp, dominant 1MM peak predominantly to 1MMM in SKP-G2.

#### 3.2.2. Interpretation of 2D ^1^H–^1^H NMR spectra

The combination of high magnetic field strength, elevated sample concentrations enabled by the low molecular weights of the fractions, and optimized NMR acquisition parameters resulted in high-quality 2D ^1^H–^1^H COSY and TOCSY spectra acquired at 80 °C (Figs. 2–5) and 65 °C (Figs. S14–S17). Following the assignment of H-1 signals for G and M and H-5 signals for G in various diads and triads within the 4.4–5.2 ppm region of the 1D ^1^H NMR spectra (Fig. 1), the chemical shifts of all protons in each sugar ring were determined from COSY cross-peaks arising from scalar couplings between neighboring protons, with the assignments further supported by TOCSY spectra, which reveal cross-peaks between protons within the same ring. In SKP-G1, the underlined G residue in the G<u>G</u>G triad exhibited the strongest signals, and cross-peaks from the underlined residues in the <u>G</u>M and <u>M</u>M diads were also detected and assigned (Figs. 2, 3, S14, and S15). In SKP-G2, the underlined M residue in the M<u>M</u>M triad exhibited the strongest signals; notably, no G<u>G</u>M triad signals were detected by 1D ^1^H NMR (Fig. 1B, Table 1), indicating that all <u>G</u>M diad signals arose from M<u>G</u>M rather than G<u>G</u>M and thereby supporting the assignment of all <u>G</u>M cross-peaks to M<u>G</u>M (Figs. 4, 5, S16, and S17). The complete sets of chemical shifts obtained from the 2D ^1^H–^1^H NMR spectra for the underlined residues in the G<u>G</u>G, M<u>G</u>M, and M<u>M</u>M triads and the <u>M</u>G diad, acquired at 65 °C and 80 °C, are summarized in Table 2. In addition, weak signals corresponding to reducing-end G and M residues were detected, and partial chemical shift data extracted from these signals are listed in Table S7. These chemical shifts obtained from 2D ^1^H–^1^H NMR enabled peak assignments in the 1D ^1^H NMR spectra beyond the 4.4–5.2 ppm region (Figs. S18 and S19).

**Table 2.**
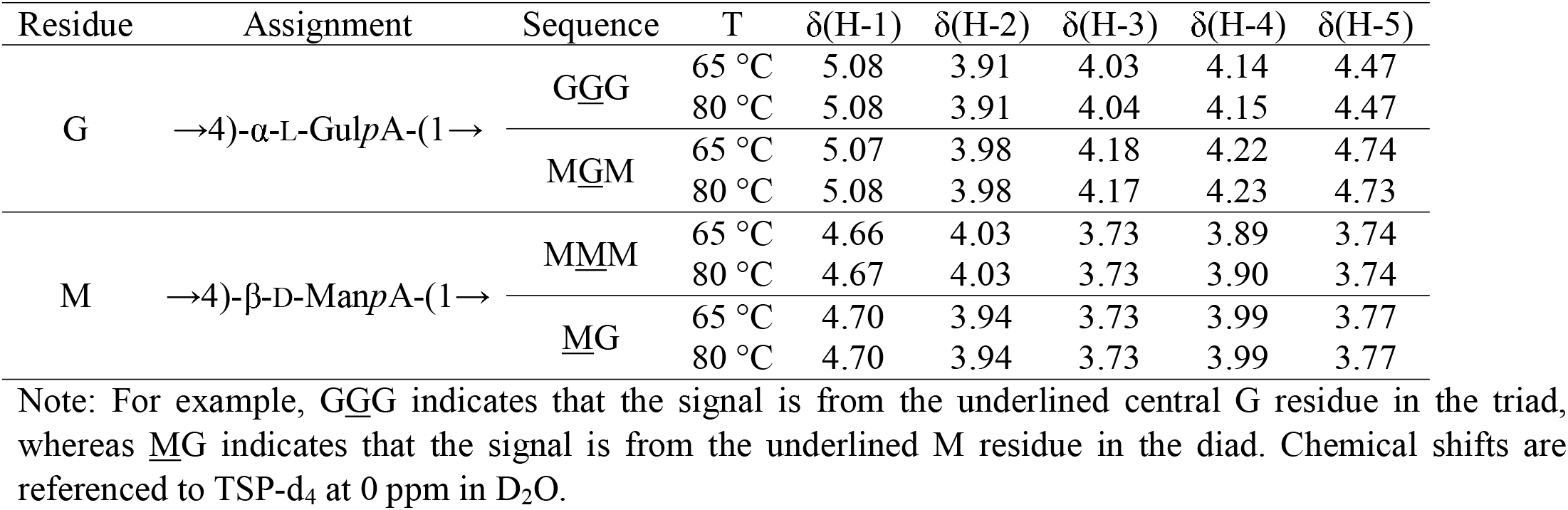
Summary of ^1^H NMR chemical shifts (δ, ppm) at 65 °C and 80 °C for M and G residues in selected alginate diad and triad sequences.

**Fig. 2.**
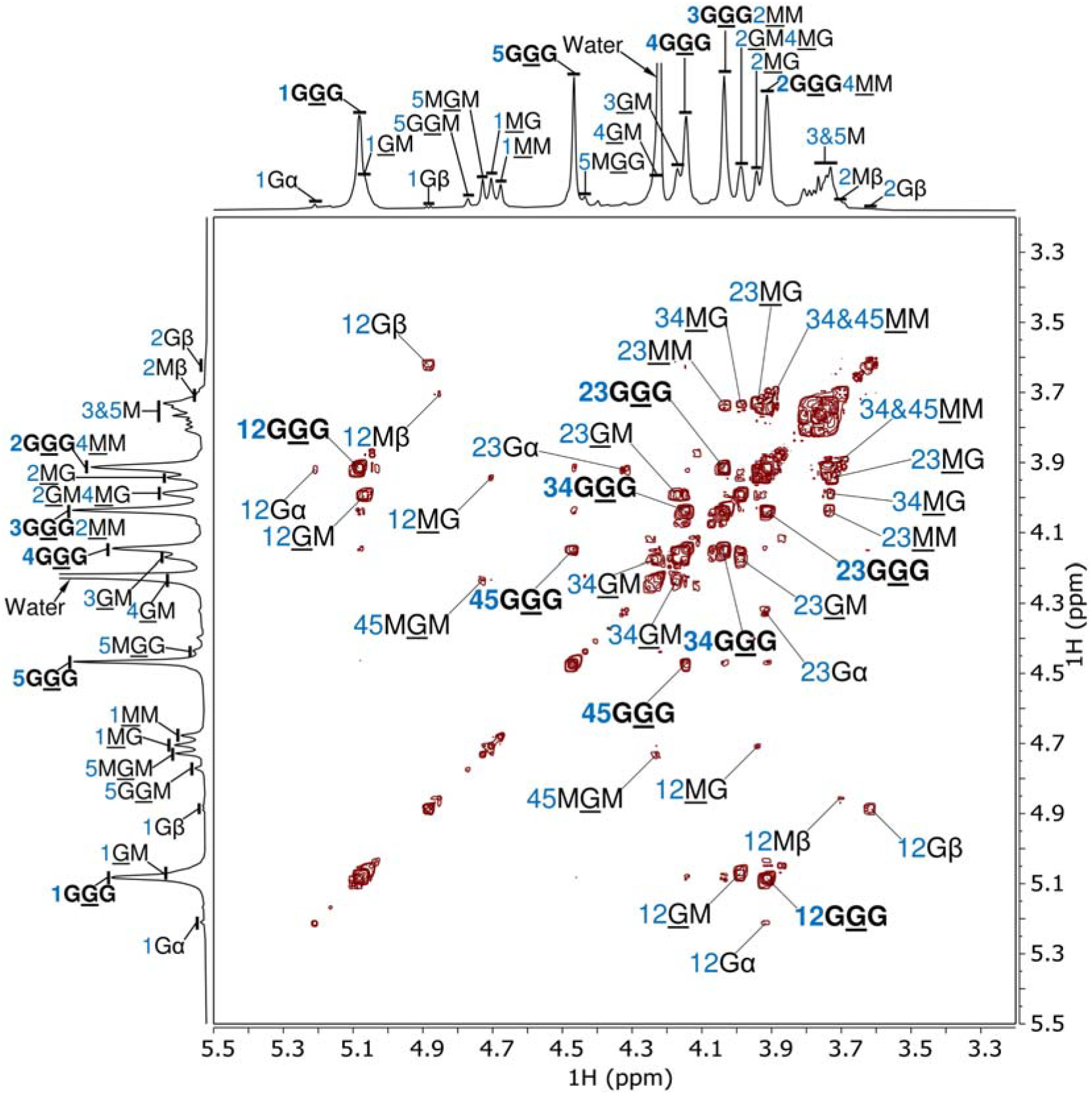
Assignment of the 2D ^1^H–^1^H COSY NMR spectrum (700 MHz, 80 °C) of SKP-G1 in D_2_O. Residues G and M denote →4)-α-L-Gul*p*A-(1→ and →4)-β-D-Man*p*A-(1→, respectively, whereas the minor residues Gα, Gβ, and Mβ represent the reducing end residues →4)-α-L-Gul*p*A, →4)-β-L-Gul*p*A, and →4)-β-D-Man*p*A, respectively. For example, the annotations “12G<u>G</u>G” and “12<u>G</u>M” indicate the cross-peaks between H-1 and H-2 of the underlined G residue in the G<u>G</u>G triad and <u>G</u>M diad, respectively, whereas “12Gα” indicates the cross-peak between H-1 and H-2 of residue Gα. In the stacked 1D ^1^H NMR spectra, the annotation “1G<u>G</u>G” denotes H-1 of the underlined central G residue in the G<u>G</u>G triad, whereas “3G<u>G</u>G2<u>M</u>M” indicates overlapping signals of “3G<u>G</u>G” and “2<u>M</u>M”. Annotations for the dominant G<u>G</u>G triad are highlighted in bold.

**Fig. 3.**
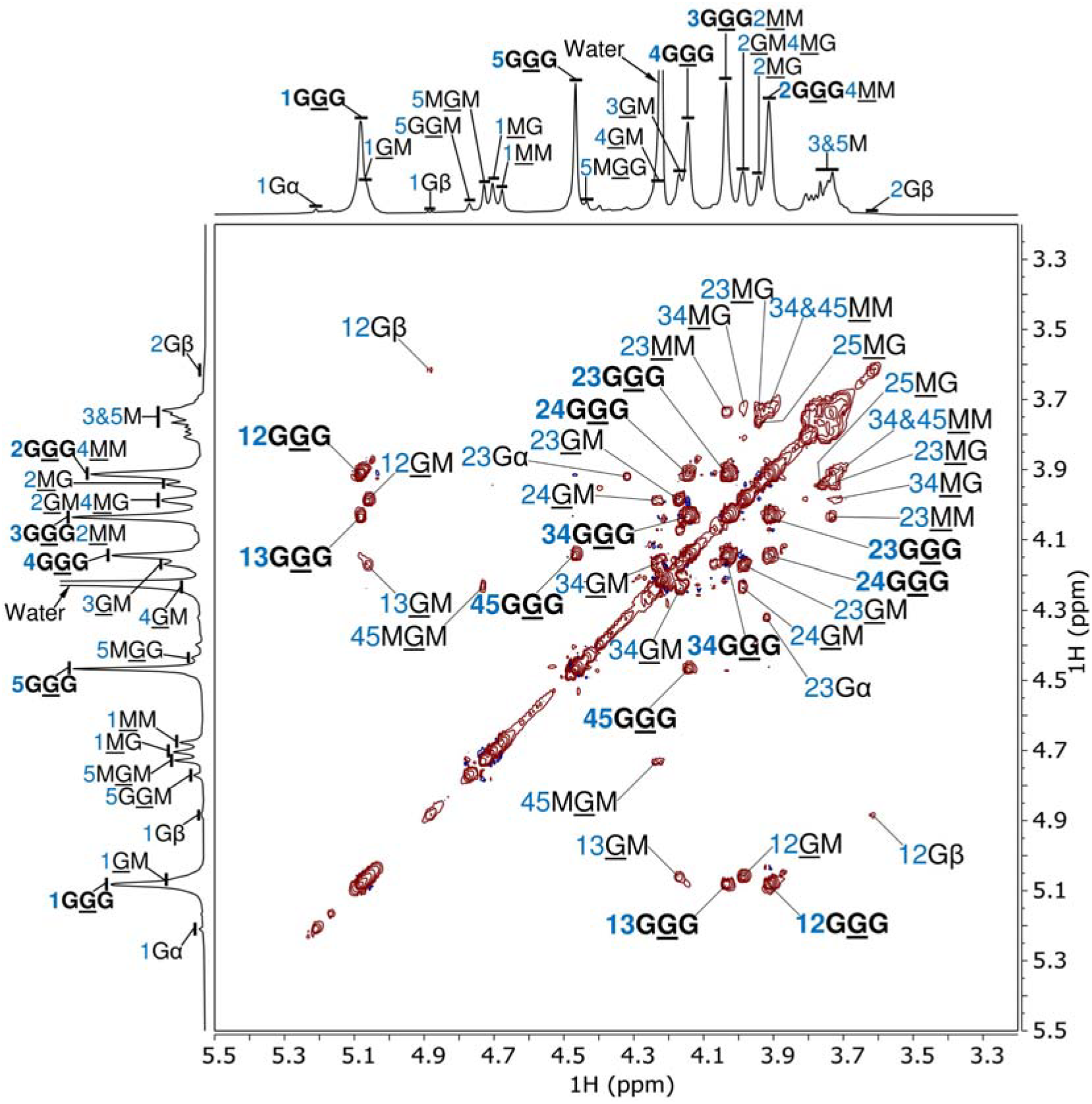
Assignment of the 2D ^1^H–^1^H TOCSY NMR spectrum (700 MHz, 80 °C) of SKP-G1 in D_2_O. Residues G and M denote →4)-α-L-Gul*p*A-(1→ and →4)-β-D-Man*p*A-(1→, respectively, whereas the minor residues Gα and Gβ represent the reducing end residues →4)-α-L-Gul*p*A and →4)-β-L-Gul*p*A, respectively. For example, the annotations “12G<u>G</u>G” and “12<u>G</u>M” indicate the cross-peaks between H-1 and H-2 of the underlined G residue in the G<u>G</u>G triad and <u>G</u>M diad, respectively, whereas “12Gβ” indicates the cross-peak between H-1 and H-2 of residue Gβ. In the stacked 1D ^1^H NMR spectra, the annotation “1G<u>G</u>G” denotes H-1 of the underlined central G residue in the G<u>G</u>G triad, whereas “3G<u>G</u>G2<u>M</u>M” indicates overlapping signals of “3G<u>G</u>G” and “2<u>M</u>M”. Annotations for the dominant G<u>G</u>G triad are highlighted in bold.

**Fig. 4.**
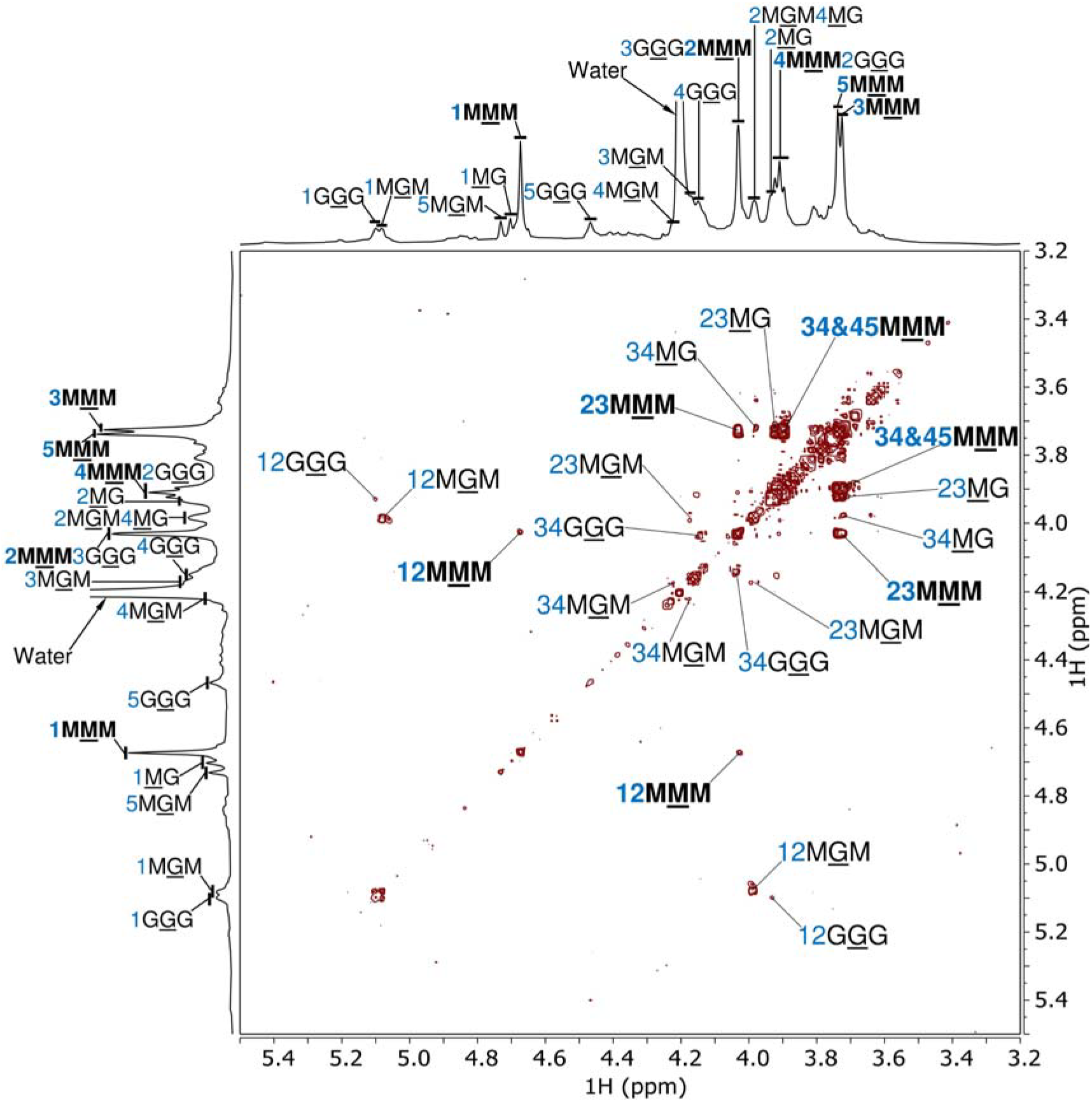
Assignment of the 2D ^1^H–^1^H COSY NMR spectrum (700 MHz, 80 °C) of SKP-G2 in D_2_O. Residues G and M denote →4)-α-L-Gul*p*A-(1→ and →4)-β-D-Man*p*A-(1→, respectively. For example, the annotations “12M<u>M</u>M” and “12G<u>G</u>G” indicate the cross-peaks between H-1 and H-2 of the underlined central M and G residues in the M<u>M</u>M and G<u>G</u>G triads, respectively. In the stacked 1D ^1^H NMR spectra, the annotation “1M<u>M</u>M” denotes H-1 of the underlined central M residue in the M<u>M</u>M triad, whereas “4M<u>M</u>M2G<u>G</u>G” indicates overlapping signals of “4M<u>M</u>M” and “2G<u>G</u>G”. Annotations for the dominant M<u>M</u>M triad are highlighted in bold.

**Fig. 5.**
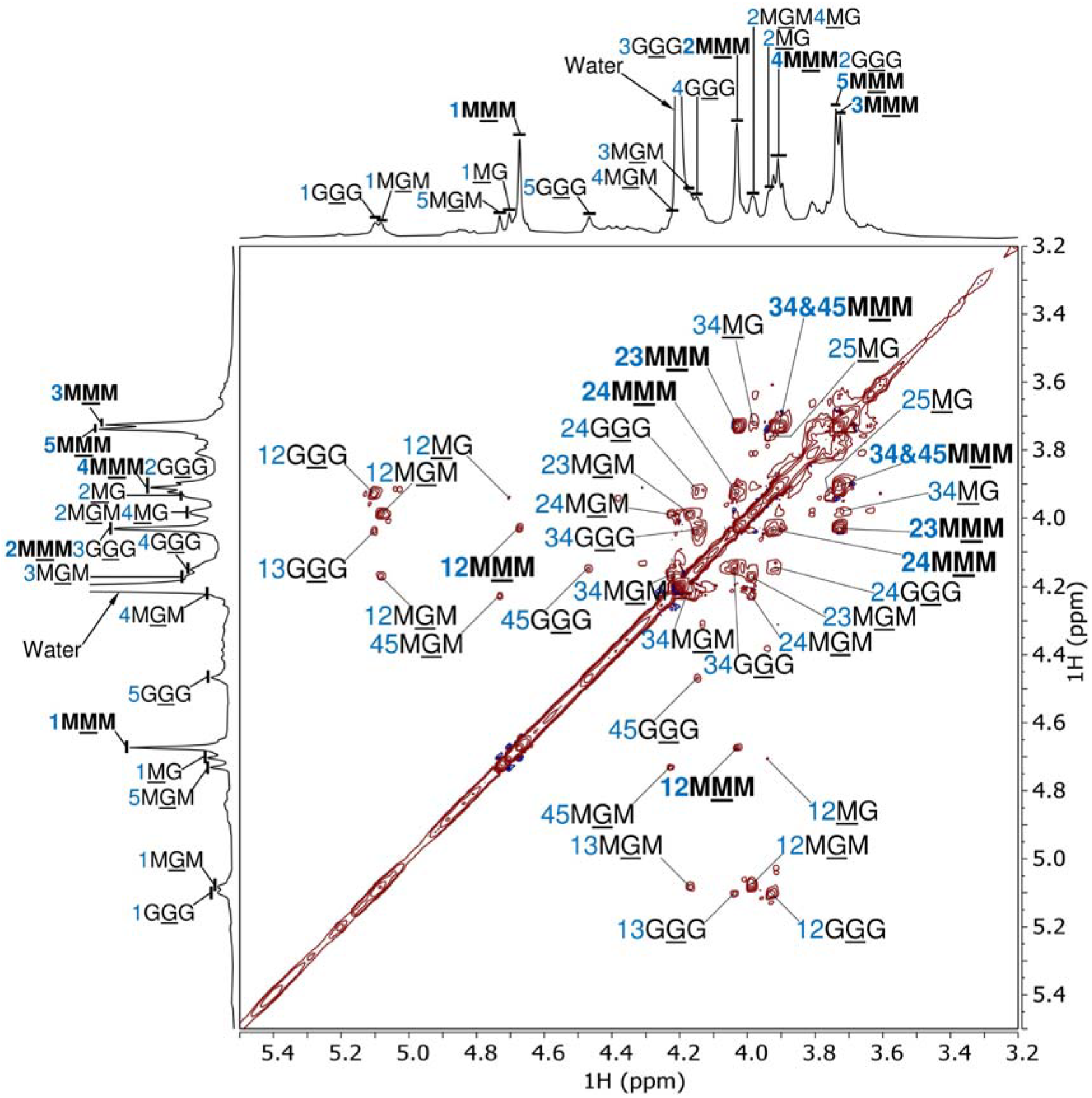
Assignment of the 2D ^1^H–^1^H TOCSY NMR spectrum (700 MHz, 80 °C) of SKP-G2 in D_2_O. Residues G and M denote →4)-α-L-Gul*p*A-(1→ and →4)-β-D-Man*p*A-(1→, respectively. For example, the annotations “12M<u>M</u>M” and “12G<u>G</u>G” indicate the cross-peaks between H-1 and H-2 of the underlined central M and G residues in the M<u>M</u>M and G<u>G</u>G triads, respectively. In the stacked 1D ^1^H NMR spectrum, the annotation “1M<u>M</u>M” denotes H-1 of the underlined central M residue in the M<u>M</u>M triad, whereas “4M<u>M</u>M2G<u>G</u>G” indicates overlapping signals of “4M<u>M</u>M” and “2G<u>G</u>G”. Annotations for the dominant M<u>M</u>M triad are highlighted in bold.

#### 3.2.3. Evaluation of chain-growth statistical models using ^1^H NMR-derived homoblock frequency data

Chain-growth statistical models, including the Bernoulli, first-order Markov, and second-order Markov chain models, provide a robust framework for predicting the sequence distribution and block architecture of linear polysaccharides such as alginate [1, 27, 28]. In these models, the probability of monomer addition is conditioned on preceding units: the first-order Markov model depends on the immediately preceding monomer; the second-order Markov model depends on the preceding diad; and the Bernoulli model assumes independence from prior units [1, 27, 28]. Traditionally, evaluation of these models has relied on NMR-derived diad and triad frequency data [27, 28]. However, a recent study introduced rigorous inequality relations that enable direct calculation of frequency ranges for homoblock polyads, extending model evaluation beyond triads [1]

For the G-block-rich SKP-G1, the frequency ranges of homo-G-blocks (G)_*n*_ were calculated from the triad frequencies *F*_GGG_ and *F*_GGM_ using inequality relation (12), whereas for the M-block-rich SKP-G2, the frequency ranges of homo-M-blocks (M)_*n*_ were calculated from the diad frequencies *F*_MM_ and *F*_MG_ using inequality relation (13), as detailed in Section 2.5. These homoblock frequency ranges, derived directly from experimental NMR data rather than from statistical approximations, provide a robust basis for evaluating chain-growth statistical models. As shown in Fig. 6, individual frequency values, such as *F*_G_, *F*_GG_, and *F*_GGG_ for SKP-G1 and *F*_M_ and *F*_MM_ for SKP-G2, are presented as discrete points, whereas homoblock frequency ranges are depicted as bars up to a DP of 18. Across all DPs, the upper limits of these bars remain constant, whereas the lower limits decrease progressively with increasing DP until they reach zero and remain constant thereafter. This trend reflects the mathematical properties of inequality relations (12) and (13), in which the maximum value is independent of DP, whereas the minimum decreases with increasing DP until it reaches zero.

**Fig. 6.**
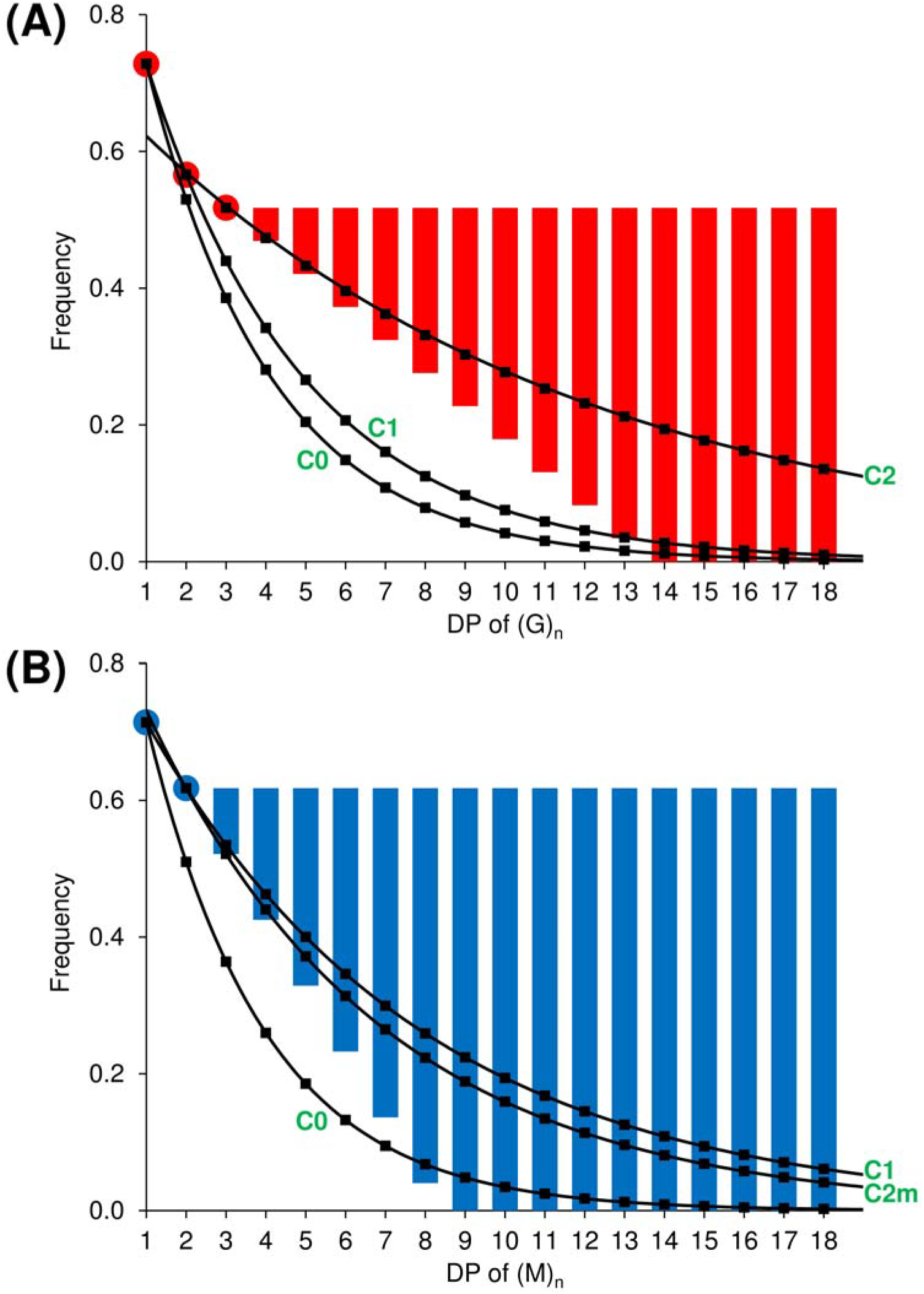
Comparison of homoblock frequency values predicted by chain-growth statistical models with NMR-derived frequency ranges for (A) SKP-G1 and (B) SKP-G2. Red bars represent the experimental frequency ranges for homo-G-blocks, (G)_*n*_, and blue bars represent those for homo-M-blocks, (M)_*n*_, with denoting DP. Black curves, denoted C0 and C1 in both panels, represent values predicted by the Bernoulli and first-order Markov chain models, respectively. The C2 curve in (A) represents the predicted values of 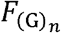, whereas the C2m curve in (B) represents the predicted minimum possible values of 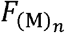, both based on the second-order Markov chain model, as detailed in Section 2.5.5.

The frequencies of (G)_*n*_ for SKP-G1 and (M)_*n*_ for SKP-G2 were predicted using the corresponding monad frequencies for the Bernoulli model according to equations (14) and (17), respectively, and the corresponding monad and diad frequencies for the first-order Markov model according to equations (15) and (18), respectively. For the second-order Markov model, the frequencies of (G)_*n*_ for SKP-G1 were predicted based on *F*_GG_ and *F*_GGG_ using equation (16), whereas for SKP-G2, where *F*_MMM_ is provided as a range rather than a single value, the minimum of that range was used to calculate the minimum possible 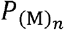 at each DP according to equation (20).

The predicted values, shown as discrete points along the model-predicted curves, were then compared with the experimental frequency ranges of homoblocks presented as bars in Fig. 6. For SKP-G1, the Bernoulli model curve (C0 in Fig. 6A) fell below the experimental points or bars for homoblock (G)_*n*_ at DPs 2–13, and the first-order Markov model curve (C1 in Fig. 6A) fell below the experimental bars for (G)_*n*_ at DPs 3–12, indicating that both models are invalid for homo-G-block growth in this G-rich fraction. In contrast, the second-order Markov model curve (C2 in Fig. 6A) remained within the experimental bars, suggesting that its validity cannot be conclusively refuted based on the available data. Similarly, for SKP-G2, the Bernoulli model curve (C0 in Fig. 6B) fell below the experimental points or bars for homoblock (M)_*n*_ across DPs 2–13, confirming its invalidity. However, both the first-order Markov model curve (C1 in Fig. 6B) and the curve representing the minimum predicted homo-M-block frequency calculated using the second-order Markov model (C2m in Fig. 6B) remained within all experimental bars shown for (M)_*n*_ in SKP-G2, indicating that the validity of these models cannot be ruled out based on the current dataset.

The ^1^H NMR results presented in Section 3.2 established the contrasting G- and M-rich block structures of SKP-G1 and SKP-G2 and provided homoblock polyad frequency ranges for evaluating chain-growth statistical models. We next investigated whether these structural differences were reflected in distinct features in the ATR-FTIR spectra and their first- and second-derivative spectra, and whether LCT-transformed derivative spectra could make these features more visually discernible and provide distinct spectral fingerprints for potential chemometric differentiation of the two fractions.

### 3.3. ATR-FTIR analysis

#### 3.3.1. Band assignment of ATR-FTIR spectra and their first- and second-derivative spectra

Derivative treatment improves spectral resolution, enabling precise identification of closely spaced bands and subtle spectral features in ATR-FTIR spectra [20–23, 31]. In the first-derivative spectrum, zero-crossing points correspond to local extrema in the original spectrum; with increasing wavenumber, a zero crossing from positive to negative indicates a local maximum (peak apex), whereas a transition from negative to positive indicates a local minimum (valley) [22, 23]. In the second-derivative spectrum, a local maximum in the original spectrum corresponds to the lowest point of a negative peak, whereas a local minimum corresponds to the highest point of a positive peak [20, 21, 31].

In the ATR-FTIR spectra (Fig. 7), SKP-G1 and SKP-G2 exhibited major broad bands at approximately 3355 and 3302 cm^−1^, respectively, corresponding to O–H stretching vibrations [72–74], along with minor bands near 2926 and 2921 cm^−1^, respectively, corresponding to C–H stretching [21, 75]. The bands at 1731 cm^−1^ (SKP-G1) and 1730 cm^−1^ (SKP-G2) were assigned to C=O stretching in carboxyl groups (–COOH) [72, 76, 77]. Signals at 1633 and 1614 cm^−1^ corresponded to O–C–O asymmetric stretching in carboxylate groups (–COO^-^) [31, 78], while those at 1403 cm^−1^ (SKP-G1) and 1410 cm^−1^ (SKP-G2) were attributed to C–OH deformation, with contributions from O–C–O symmetric stretching in carboxylate groups [78]. These findings indicated the coexistence of protonated and deprotonated carboxyl groups in both samples, consistent with previous studies [79, 80]. Slight shifts in O–C–O asymmetric and symmetric stretching between the two samples likely resulted from differences in metal ion coordination and block structures. Relevant bands were also observed in the corresponding regions of the first- and second-derivative spectra (Figs. S20–S23).

**Fig. 7.**
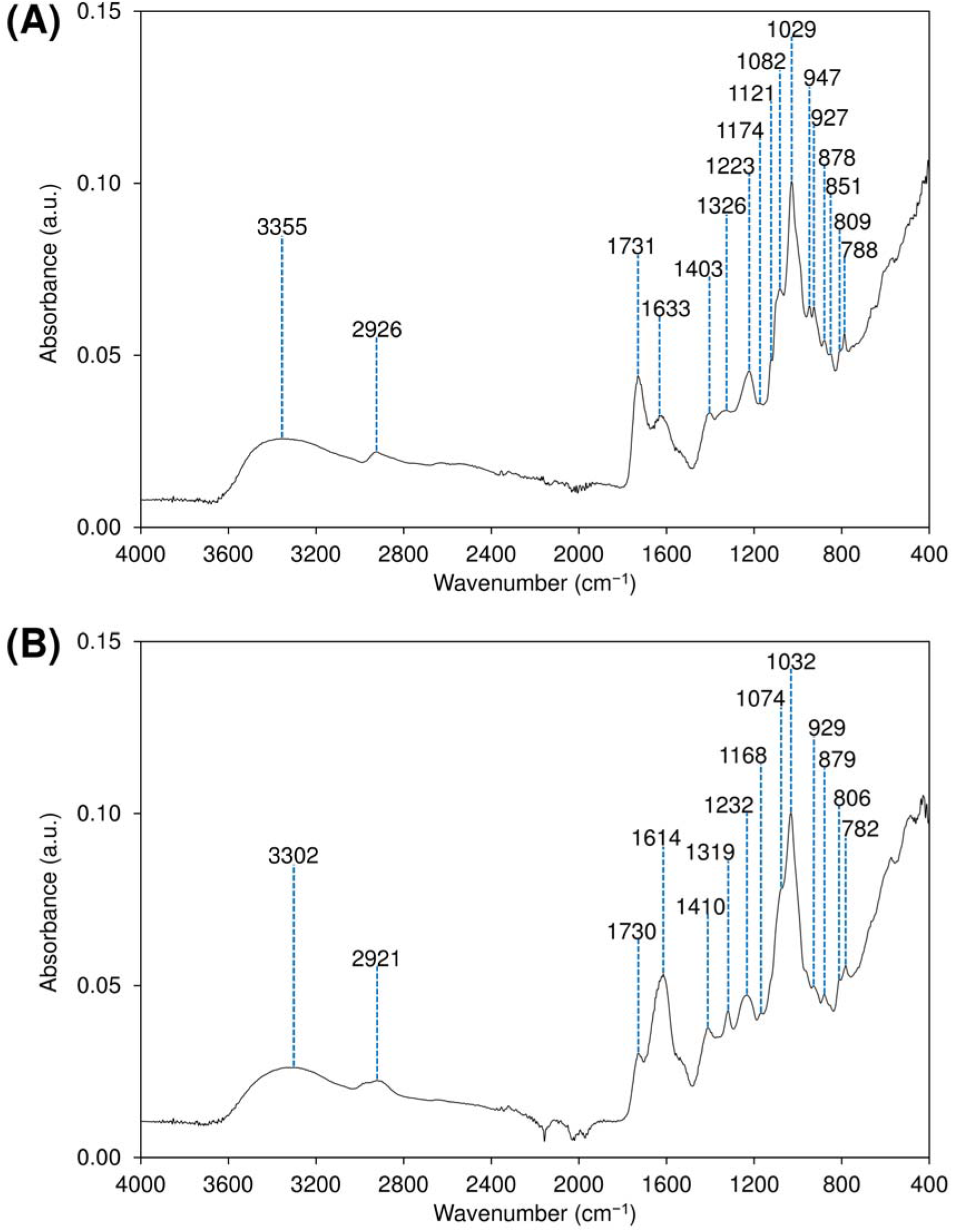
ATR-FTIR spectra of (A) SKP-G1 and (B) SKP-G2. The blue vertical lines indicate the wavenumbers of selected bands.

The bands at 1326 cm^−1^ (SKP-G1) and 1319 cm^−1^ (SKP-G2) were attributed to C–C–H and O–C–H deformation in pyranose rings [73, 81], with the latter band associated with M residues [13, 14, 25]. The band at 1223 cm^−1^ in the SKP-G1 spectrum (Fig. 7A) corresponded to C–O stretching in carboxylic acids [82]. A small band at 1232 cm^−1^ in SKP-G2 (Fig. 7B) likely originated from sulfated esters (S=O) of fucoidan, a common minor contaminant in extracted alginate [76]. Notably, although this band could also arise from amide III vibrations in proteins, the absence of amide I (C=O at 1670 cm^−1^) and amide II (N–H bending at 1540 cm^−1^) suggested that substantial protein contamination was unlikely [83]. The band at 1121 cm^−1^ in SKP-G1 (Fig. 7A) was assigned to C–OH stretching vibrations [20, 77]. Weak bands at 1174 cm^−1^ (SKP-G1) and 1168 cm^−1^ (SKP-G2) were attributed to C–O stretching, C–C stretching, and C–C–C bending in the pyranose rings of uronic acids [72, 84], as well as to C–O symmetric stretching of glycosidic linkages [31].

The strong band centered around 1030 cm^−1^ in alginate and alginic acid was primarily attributed to C–O stretching in the pyranose ring [21, 31, 72, 78, 85], although some studies associated it with pyranose-ring C–C stretching [31, 73, 85, 86] or O–H bending [20], assigned it to asymmetric C–O–C stretching in glycosidic linkages [74, 87], or left it unclassified because of its complex bonding environment [84, 88]. The major shoulder near 1080 cm^−1^ was typically linked to C–O and C–C stretching vibrations of pyranuronate rings [21, 31, 81, 82, 84, 85], although other reports attributed it to carboxylic C–O stretching [89, 90], C–O stretching in glycosidic C–O–C bonds [86], or C–O stretching in weakly hydrogen-bonded C–OH structures [91]. SKP-G1 exhibited a sharp, intense band at 1029 cm^−1^ and a major shoulder at 1082 cm^−1^, while SKP-G2 displayed a sharp band at 1032 cm^−1^ with a strong shoulder at 1074 cm^−1^ (Fig. 7); these features closely aligned with zero-crossing points in the corresponding first-derivative spectra (Fig. 8). In the second-derivative spectra (Fig. 9), the 1029 and 1082 cm^−1^ bands in SKP-G1 resolved into pairs at 1032/1025 and 1084/1070 cm^−1^, respectively, while the 1032 and 1074 cm^−1^ bands in SKP-G2 resolved into pairs at 1028/1036 and 1079/1066 cm^−1^, respectively, reflecting the underlying bonding complexity. Slight shifts between fractions (*e.g*., 1032 cm^−1^ vs. 1029 cm^−1^) may reflect differences in metal ion binding, monomer composition, and block structures [21, 74, 88, 92]. Some studies linked the 1030 cm^−1^ band to G residues and its shoulder at 1080 cm^−1^ to M residues [72, 86], while others assigned the bands in the opposite manner [25, 85]. However, spectral analyses using fractions with varying M/G ratios or block structures did not support direct residue assignment [75, 92], and monomer ratio estimation based on these bands was deemed unreliable [93]. For instance, absorbance near 1080 cm^−1^ exhibited minimal variation with changes in the M/G ratio because it reflected pyranose-ring stretching in both M and G residues [88], a conclusion further supported by the present study.

**Fig. 8.**
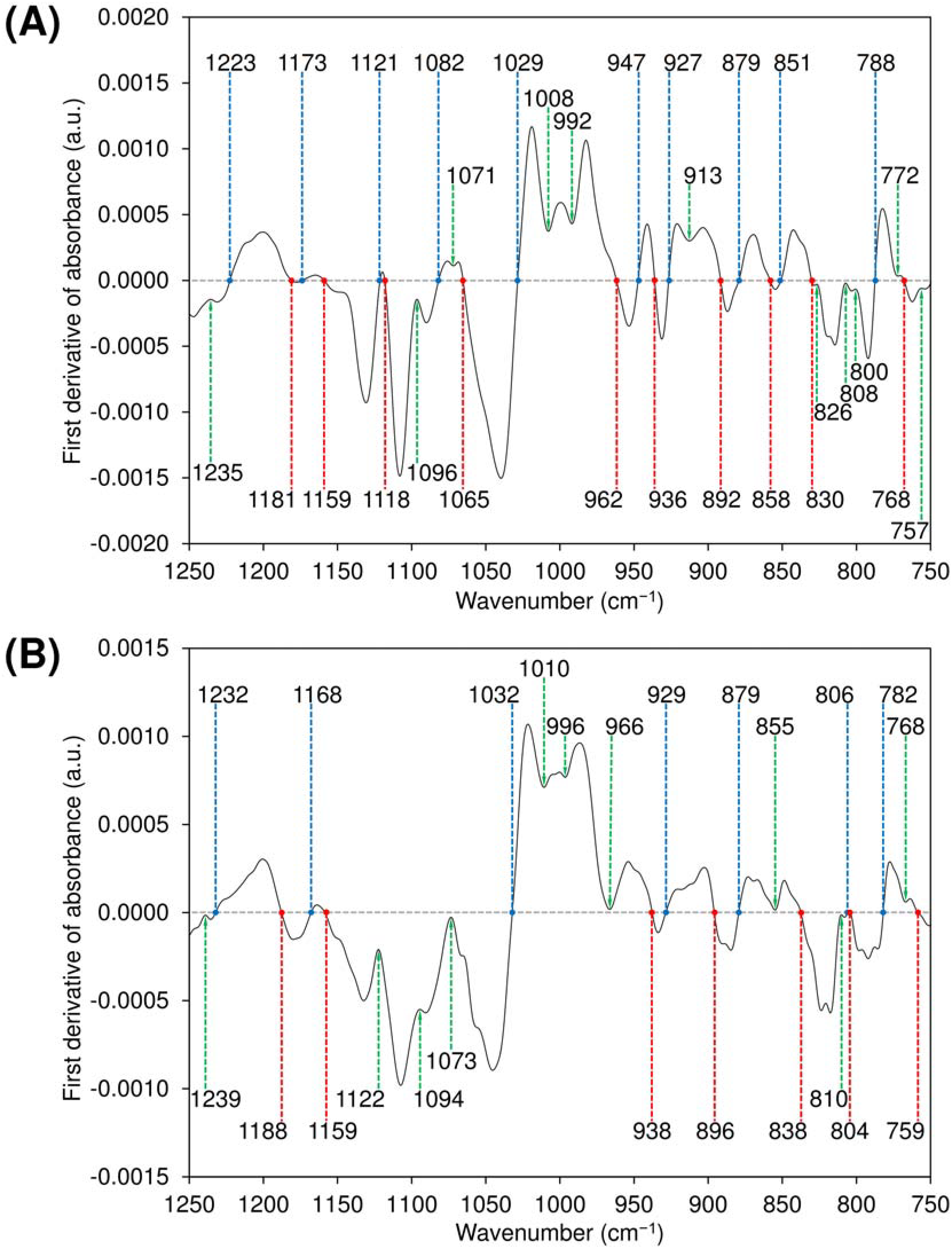
First-derivative ATR-FTIR spectra of (A) SKP-G1 and (B) SKP-G2 in the 750–1250 cm^−1^ region. The blue line indicates a zero crossing from positive to negative as the wavenumber increases, whereas the red line indicates a zero crossing in the opposite direction, from negative to positive. The green line marks the local extremum closest to zero.

**Fig. 9.**
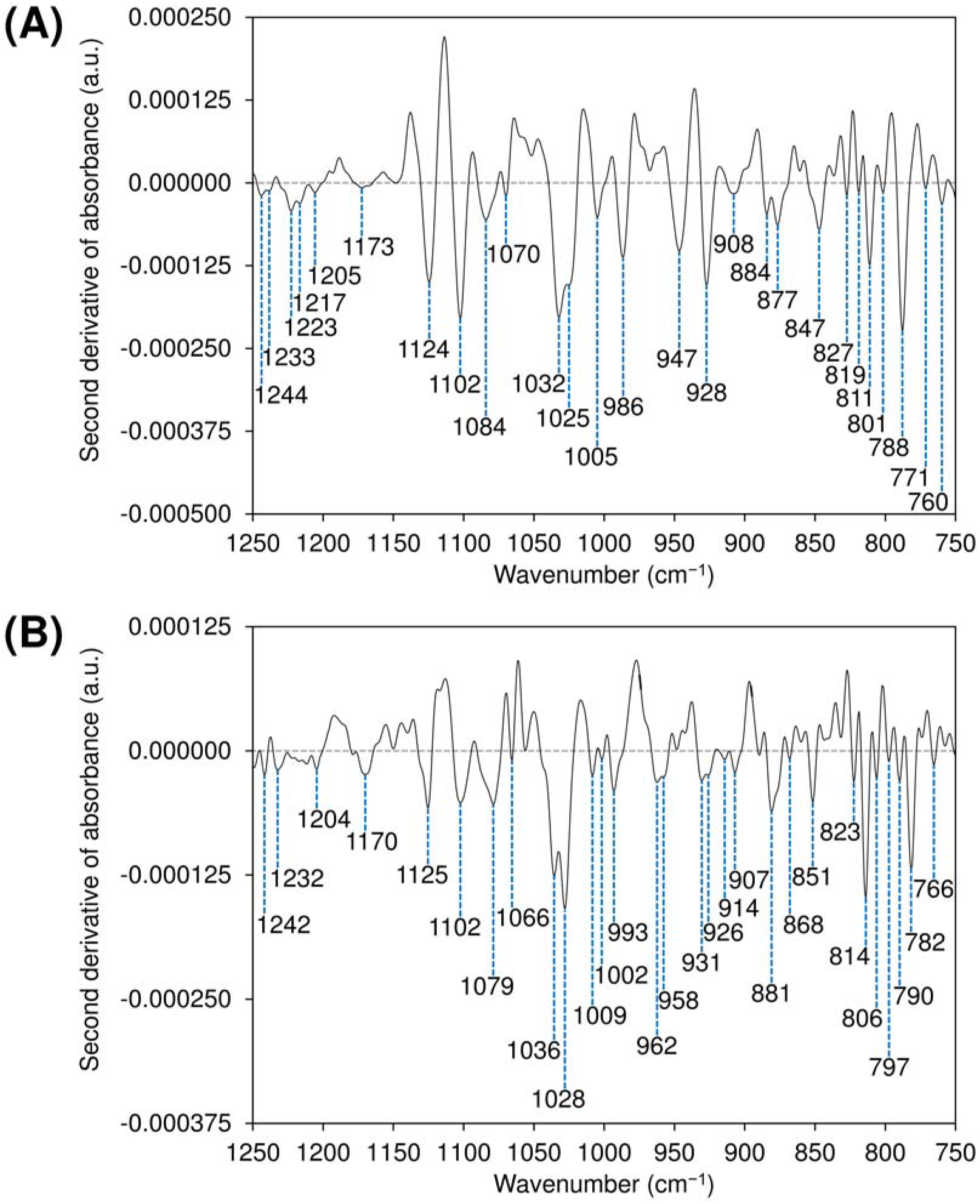
Second-derivative ATR-FTIR spectra of (A) SKP-G1 and (B) SKP-G2 in the 750–1250 cm^−1^ region. The blue vertical lines indicate the wavenumbers of selected negative second-derivative bands.

The small bands at 947 and 927 cm^−1^ in SKP-G1 and at 929 cm^−1^ in SKP-G2 (Fig. 7), along with their corresponding first- and second-derivative signals (Figs. 8 and 9), were attributed to C–O stretching vibrations of uronic acids [21, 78, 93]. The band at 878 cm^−1^ in SKP-G1 (Fig. 7A), which resolved into bands at 884 and 877 cm^−1^ in the second-derivative spectra (Fig. 9), and the band at 879 cm^−1^ in SKP-G2 (Fig. 7B), which resolved into bands at 881 and 868 cm^−1^, were assigned to C1–H deformation in M residues [72, 73, 75]. Bands at 809 cm^−1^ (SKP-G1) and 806 cm^−1^ (SKP-G2), along with their derivative signals, were characteristic of M residues [72–74], whereas bands at 788 cm^−1^ (SKP-G1) and 782 cm^−1^ (SKP-G2), along with their corresponding derivative features, were indicative of G residues [84, 92].

#### 3.3.2. LCT-transformed derivative ATR-FTIR spectra as a fingerprinting tool

The LCT algorithm was previously developed in our study of per-*O*-ethylated carrageenans as part of a novel workflow for post-processing 2D ^1^H–^13^C HSQC NMR spectra [2]. In that study, first-derivative processing followed by LCT was applied to individual 1D ^1^H spectral slices at each ^13^C spectral position, producing a 2D spectral representation with enhanced local resolution that facilitated the subsequent extraction of ^1^H and ^13^C chemical shifts. In the present study, the LCT transformation was extended to ATR-FTIR spectroscopy and applied separately to the first- and second-derivative spectra of SKP-G1 and SKP-G2 to investigate its utility for spectral feature extraction and visualization.

The mathematical form of equations (21)–(25) in Section 2.6.2 ensures that as a derivative value approaches zero, its corresponding LCT-converted value increases, enabling the LCT algorithm to extract major features from derivative spectra by assigning greater weight to points closer to zero. Zero-crossing points in the derivative spectrum correspond to sharp spikes in the LCT-transformed spectrum, while peaks near but not crossing the zero line generate weaker, broader peaks in the LCT-processed signals. In both cases, peak intensity and sharpness depend on the proximity of the derivative value to zero and the rate at which neighboring derivative values deviate from zero. Additionally, the appearance of LCT-processed derivative spectra depends on the size of the selected Savitzky-Golay smoothing window applied to the raw spectrum prior to differentiation, as it governs the extent of noise reduction and suppression of minor signals. LCT-transformed spectra for SKP-G1 and SKP-G2 were presented separately for the ranges 750–1250 cm^−1^ (Figs. 10 and 11), 1250–1750 cm^−1^ (Figs. S24 and S25), and 2800–3200 cm^−1^ (Figs. S26 and S27). The effect of increasing the smoothing window was more pronounced in the latter two regions, which exhibited higher noise levels than the 750–1250 cm^−1^ region.

**Fig. 10.**
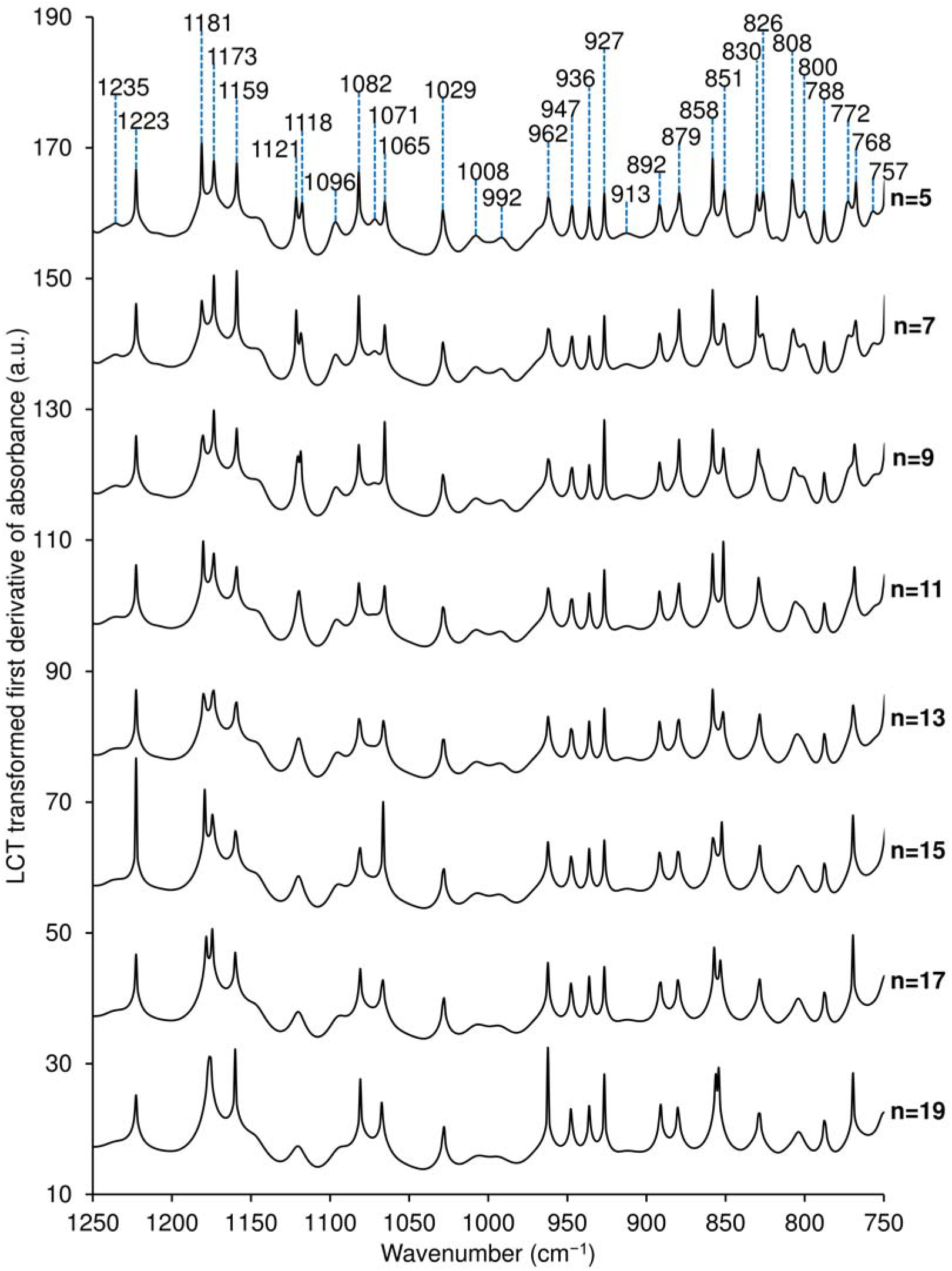
LCT-transformed first-derivative ATR-FTIR spectra of SKP-G1 in the 750–1250 cm^−1^ region, with the lowercase letter n denoting increasing Savitzky-Golay smoothing points from 5 to 19.

**Fig. 11.**
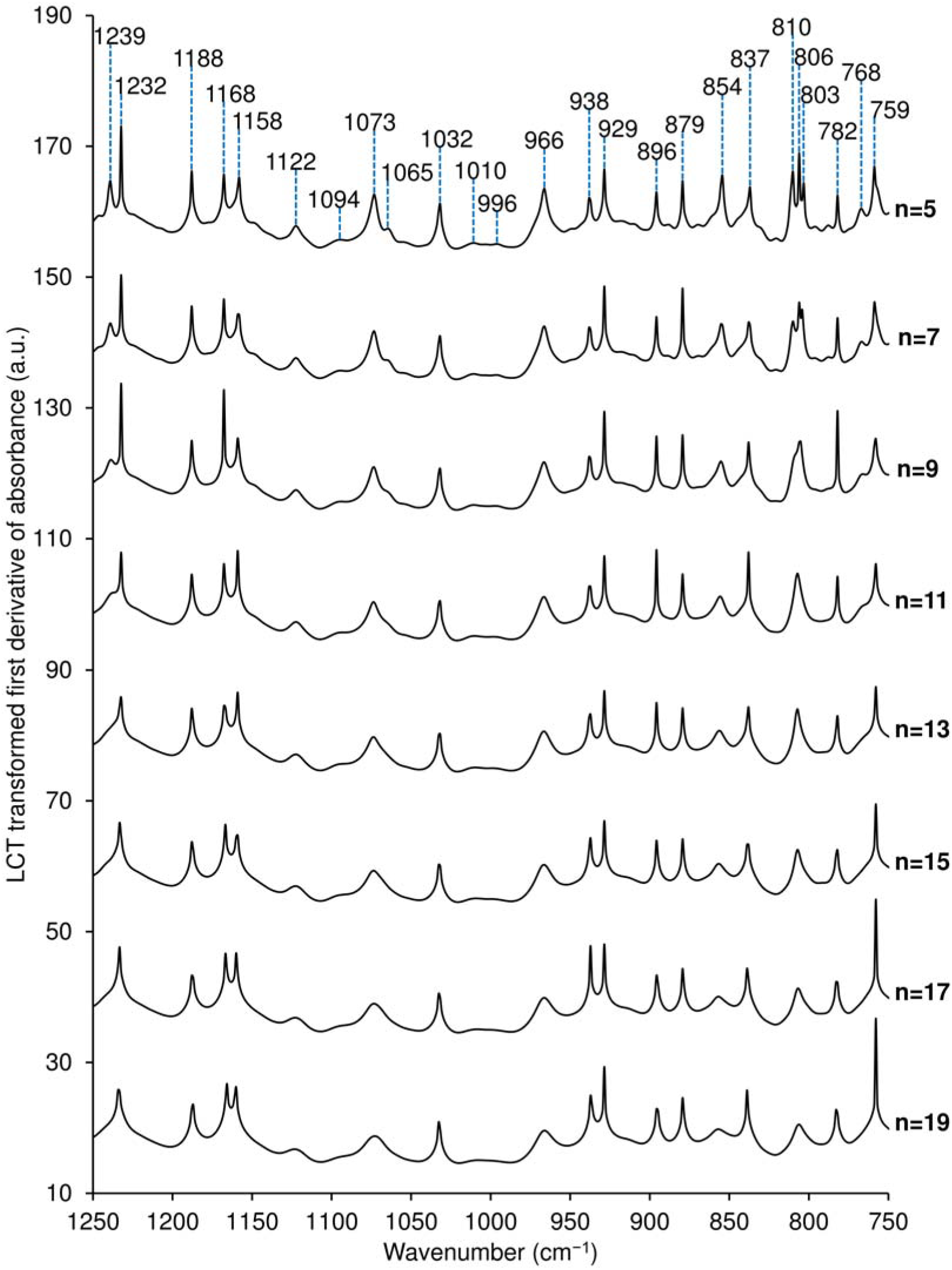
LCT-transformed first-derivative ATR-FTIR spectra of SKP-G2 in the 750–1250 cm^−1^ region, with the lowercase letter n denoting increasing Savitzky-Golay smoothing points from 5 to 19.

The LCT-transformed first-derivative ATR-FTIR spectra of SKP-G1 (Figs. 10, S24, and S26) and SKP-G2 (Figs. 11, S25, and S27) exhibited sharp spikes at zero-crossing points in their first-derivative spectra, corresponding to alternating local maxima and minima in the Savitzky-Golay-smoothed original spectra. This spiking effect made spectral features more visually discernible and facilitated identification of peaks in the original spectra and their corresponding zero-crossing wavenumbers in the first-derivative spectra. In contrast, broader, weaker peaks arose from local extrema of derivative features that approached but did not cross the zero line, reflecting unresolved shoulders and troughs in the smoothed original spectra. These features were particularly useful for identifying unresolved shoulder signals in the original spectra.

Distinct spectral features also emerged in the LCT-transformed second-derivative ATR-FTIR spectra of SKP-G1 and SKP-G2 (Figs. 12 and 13). Sharp spikes appeared at zero-crossing points in the second-derivative spectra, corresponding to alternating local maxima and minima in the first-derivative spectra. In addition, these zero-crossing points coincided with inflection points in the smoothed original spectra, marking transitions in concavity between concave-up and concave-down regions. Broader, weaker peaks resulted from local extrema of derivative features that approached but did not cross the zero line in the second-derivative spectra, indicating unresolved shoulders and troughs in the first-derivative spectra.

**Fig. 12.**
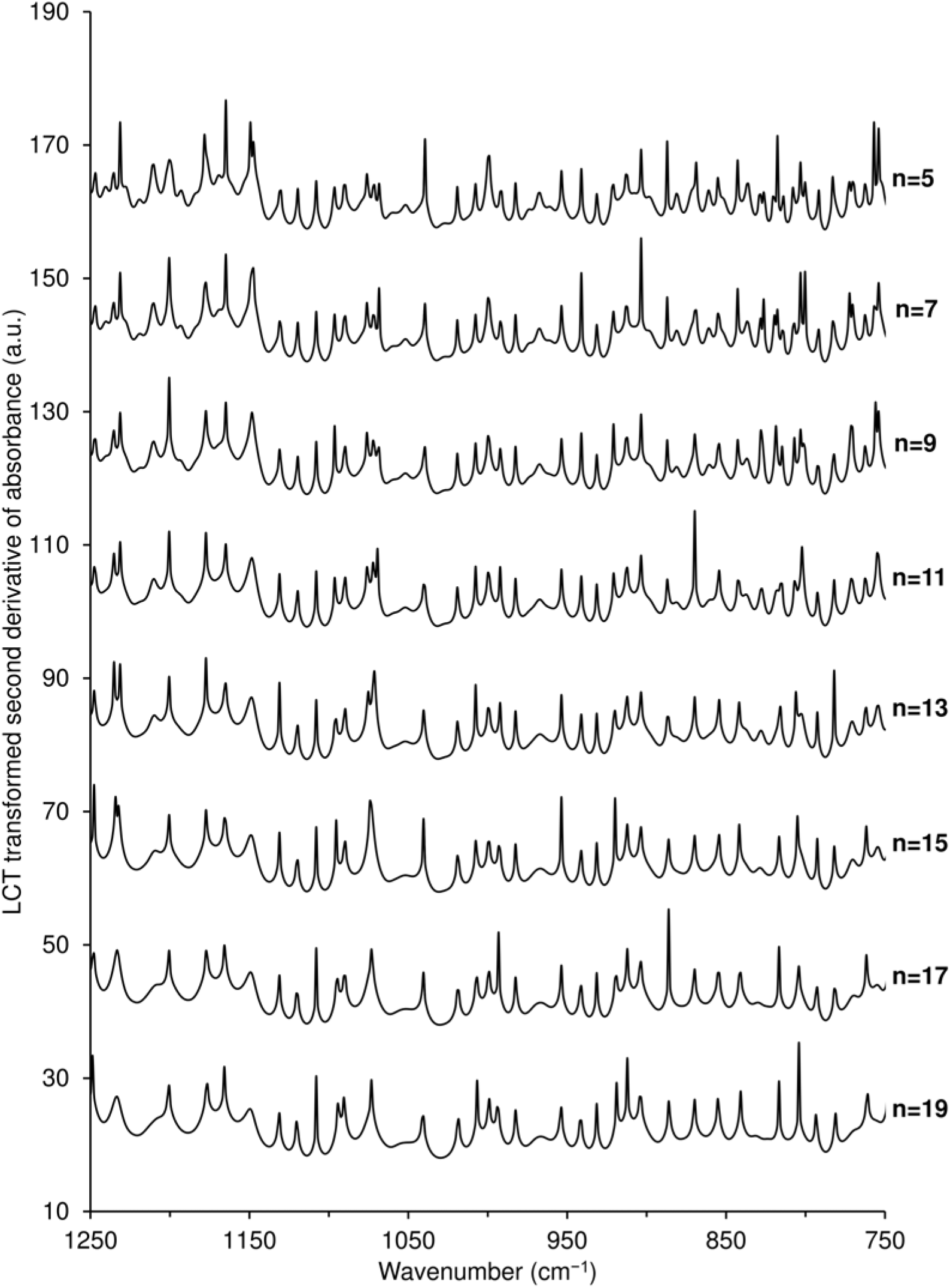
LCT-transformed second-derivative ATR-FTIR spectra of SKP-G1 in the 750–1250 cm^−1^ region, with the lowercase letter n denoting increasing Savitzky-Golay smoothing points from 5 to 19.

**Fig. 13.**
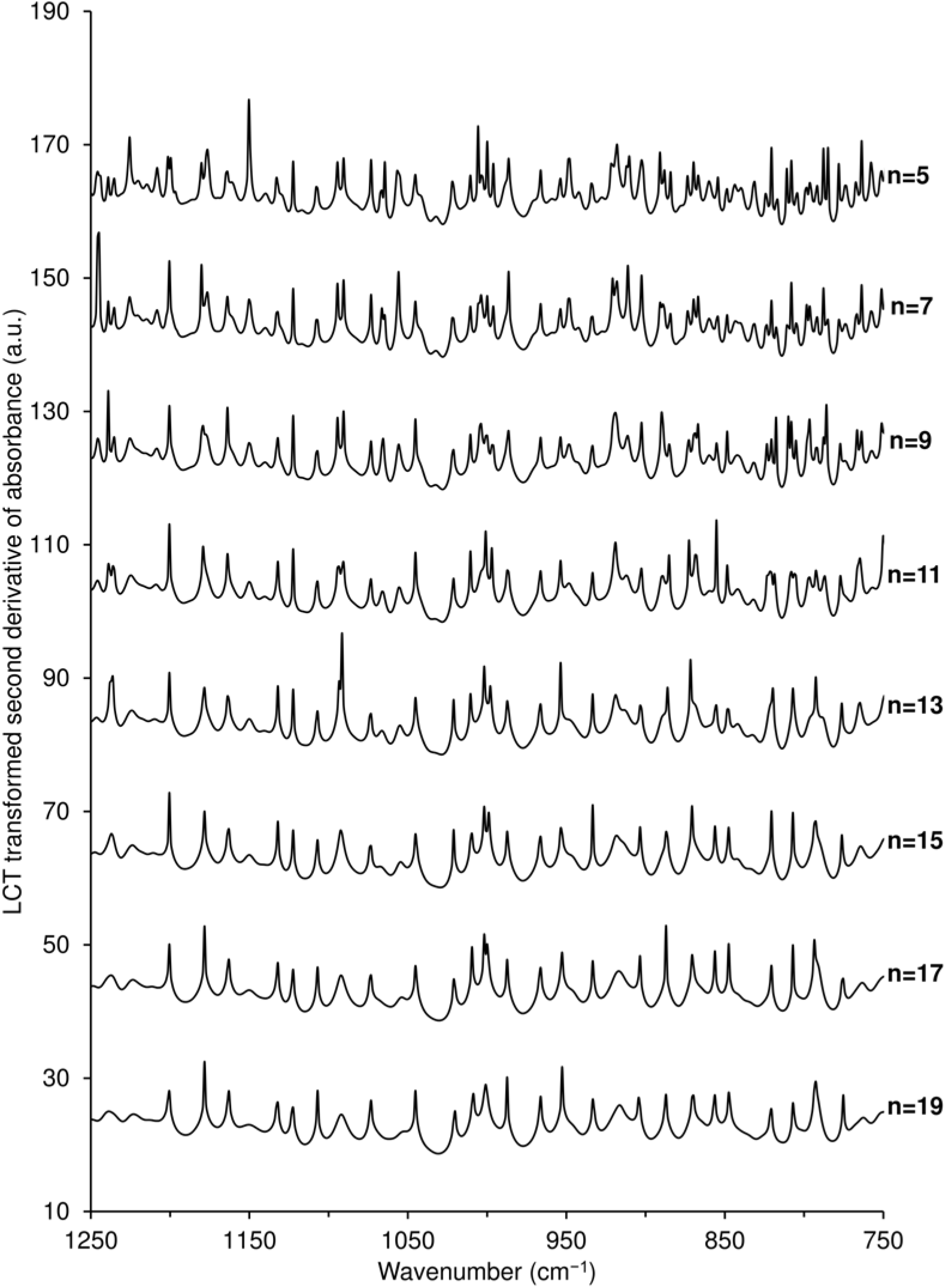
LCT-transformed second-derivative ATR-FTIR spectra of SKP-G2 in the 750–1250 cm^−1^ region, with the lowercase letter n denoting increasing Savitzky-Golay smoothing points from 5 to 19.

Both LCT-transformed first- and second-derivative ATR-FTIR spectra preserved key features of the original spectra while enhancing the visibility of previously indistinct details, with the latter highlighting additional spectral features owing to the additional differentiation step. LCT-processed derivative spectra emphasized features often overlooked in conventional analysis, including valleys and shoulder troughs, thereby enabling effective spectral feature extraction and visualization. The distinct spectral patterns of SKP-G1 and SKP-G2 suggest that LCT could provide a fingerprinting approach for characterizing alginic acid or alginate samples with varying M/G ratios and block structures.

It is important to note that, in the fundamental equation (21), the LCT-converted value *Z* is a monotonically increasing function of cos*θ*, where cos*θ* ranges from 0 to 1. As the absolute value of the derivative, |*D*|, approaches 0, cos*θ* approaches 1, and *Z* tends toward +∞, becoming undefined. To ensure practical applicability, a threshold *T* was introduced to cap the value of *Z*, as detailed in Section 2.6.2. Conversely, as |*D*| increases toward +∞, cos*θ* approaches 0, and *Z* tends toward −∞. A zero crossing of *Z* occurs when cos*θ* = e^−1^, corresponding to 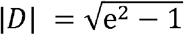 (approximately 2.53). This indicates that *Z* is positive when |*D*| is smaller than 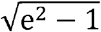 and negative when |*D*| is greater than 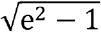. In this study, the maximum observed |*D*| for the first-derivative analysis was on the order of 10^−2^, whereas those for the second-derivative analysis were even smaller. Thus, all LCT-converted values obtained in the current study remained strictly positive. For broader applications beyond ATR-FTIR-based fingerprinting, particularly when processing derivative values exceeding 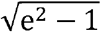, an additional constraint may be applied to ensure 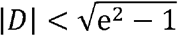 if maintaining positive *Z* values is required.

#### 3.3.3. PCA of derivative ATR-FTIR data with and without LCT

PCA was applied to assess the variation introduced by different Savitzky-Golay smoothing window sizes in the first- and second-derivative spectra of each fraction, with and without LCT conversion (Fig. S28), and to explore its utility for differentiating SKP-G1 and SKP-G2 based on these spectra as a proof of concept. PCA was performed over the 750–1250 cm^−1^ wavenumber range because of its higher peak density, greater spectral information content, and superior signal-to-noise ratio compared with other regions of the spectra.

For overall smoothing-induced variability, PCA of the first-derivative data (Fig. S28A) revealed broad dispersion of SKP-G1 along the dominant PC1 axis, whereas SKP-G2 exhibited a narrower distribution along PC1. Following LCT conversion (Fig. S28B), SKP-G1 showed substantially tighter clustering along PC1, accompanied by a slight expansion along PC2, whereas SKP-G2 remained narrowly distributed along PC1 with little change along PC2. Compared with the first-derivative data (Fig. S28A), the second-derivative data (Fig. S28C) exhibited a different dispersion pattern, with SKP-G1 showing reduced dispersion along PC2, whereas SKP-G2 showed increased dispersion along PC2 while remaining narrowly distributed along PC1. Following LCT conversion of the second-derivative data (Fig. S28D), SKP-G1 exhibited tighter clustering along PC1 with little change along PC2, whereas SKP-G2 showed a slight expansion along both axes. Overall, these visual patterns suggest that LCT may reduce sensitivity to changes in smoothing window size, particularly for SKP-G1 along PC1, which accounted for the largest proportion of variance in each PCA analysis. However, the effects on SKP-G2 and along PC2 were less consistent, and the two fractions remained visually distinct in all four PCA score plots.

For within-fraction variation under individual smoothing settings, the PCA score plots revealed differences in the dispersion of replicate measurements across smoothing window sizes (Fig. S28). However, a consistent reduction in replicate dispersion with increasing window size from 5 to 19 points was not evident for both fractions or derivative orders. Following LCT conversion, tighter clustering along PC1 was apparent under some smoothing conditions, particularly for SKP-G1, although this pattern was not consistent across all conditions (Figs. S28B and S28D). To facilitate visualization of the individual smoothing conditions with reduced overlap, a simplified plot showing three selected smoothing windows (5, 11, and 19 points) is provided in Fig. 14. The two fractions remained visually distinct along PC1 in all four PCA score plots, while their within-fraction dispersion varied among the different spectral treatments.

**Fig. 14.**
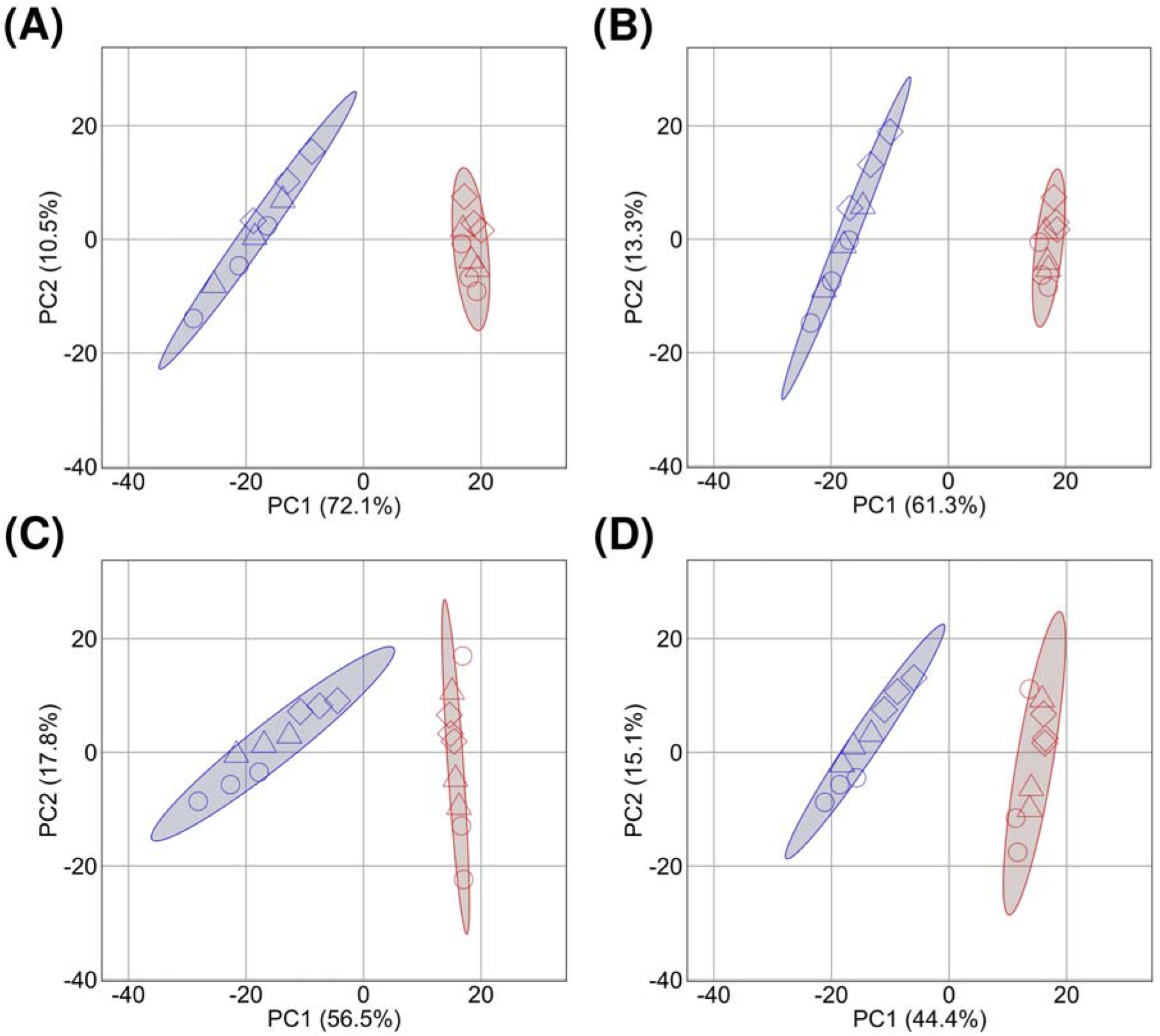
PCA score plots for SKP-G1 (blue) and SKP-G2 (red) based on (A) first-derivative ATR-FTIR spectra, (B) LCT-transformed first-derivative ATR-FTIR spectra, (C) second-derivative ATR-FTIR spectra, and (D) LCT-transformed second-derivative ATR-FTIR spectra. Symbols in the shapes of a circle, triangle, and diamond represent Savitzky-Golay smoothing windows of 5, 11, and 19 points, respectively. The 95% data ellipses are calculated from the pooled PCA scores across all three selected smoothing windows for each fraction.

Despite the limited dataset of three independent measurements per fraction, SKP-G1 and SKP-G2 remained visually distinct, primarily along PC1, in the PCA score plots generated using 5-point and 19-point smoothing windows (Figs. S29 and S30, respectively). For SKP-G1, LCT conversion resulted in visibly tighter clustering along PC1 in both first- and second-derivative analyses, although this pattern was less consistent for SKP-G2. Given the limited number of independent measurements, these findings should be considered exploratory and require further validation using larger datasets. Nevertheless, this proof-of-concept study illustrates the potential of the approach for distinguishing alginate fractions with different structural characteristics.

## 4. Conclusions

In conclusion, two-step gelation of a hot-water extract of sugar kelp AIR, involving acetic acid-induced gelation followed by freeze–thaw-induced gelation, yielded G-block-rich and M-block-rich alginic acid fractions, providing contrasting materials for the characterization of distinct alginate block structures. High-field (700 MHz) 2D ^1^H–^1^H COSY and TOCSY NMR facilitated comprehensive assignment of homoblock triad signals. Previously established inequality relations [1] were applied to diad and triad frequencies derived from 1D ^1^H NMR spectra to calculate constrained frequency ranges for homoblock polyads beyond the triad level. These experimentally derived frequency ranges provided a basis for evaluating predictions from Bernoulli and Markov chain-growth statistical models for homo-G and homo-M-block elongation over a selected range of lengths up to DP 18. Applying the recently developed LCT algorithm [2] to first- and second-derivative ATR-FTIR spectra made subtle spectral features more visually discernible and generated distinct fingerprints for the two fractions. A proof-of-concept application of LCT-transformed derivative ATR-FTIR spectra to chemometric differentiation yielded distinct PCA score distributions for the two fractions. Together, these improved ^1^H NMR and ATR-FTIR approaches broaden the analytical toolkit for alginate block structure characterization. From an industrial perspective, these approaches could support improved structural characterization and quality assessment of seaweed-derived alginates, facilitating their selection for applications where alginate block composition and sequence influence gelation behavior and material properties.

## CRediT authorship contribution statement

Conceptualization, X.X. and D.W.A.; methodology, X.X., T.M., and D.W.A.; software, X.X. and B.B.; validation, X.X., J.P.T., T.M., S.C.S., and D.W.A.; formal analysis, X.X. and B.B.; investigation, X.X., B.B., K.E.L., and V.W.; resources, D.W.A.; data curation, X.X., B.B., and K.E.L.; writing - original draft preparation, X.X.; writing - review and editing, X.X., T.M., and D.W.A.; visualization, X.X. and B.B.; supervision, D.W.A.; project administration, D.W.A.; funding acquisition, D.W.A. All authors have read and approved the final version of the manuscript.

## Data availability

Data will be made available on request.

## Acknowledgments

We thank Cascadia Seaweed Corporation for providing the dry sugar kelp and the HPSEC system.

## Appendix A. Supplementary data

The accompanying “Supplementary Information” includes Scheme S1, Figures S1–S30, Tables S1–S7, and R scripts for PCA, providing additional information on the isolation, chromatographic analysis, ^1^H NMR characterization, and ATR-FTIR analysis of the SKP-G1 and SKP-G2 fractions.

