## Supplementary Information for "Improved ^1^H NMR and ATR-FTIR approaches for structural characterization of high-G-block and high-M-block alginic acids isolated from sugar kelp by a two-step gelation method"

**Crown copyright statement**

© His Majesty the King in Right of Canada, as represented by the Minister of Agriculture and Agri-Food, 2026.

Crown copyright applies to contributions made by employees of Agriculture and Agri-Food Canada (AAFC) in the course of their official duties. The AAFC-affiliated authors of this manuscript are Xiaohui Xing (X.X.), Barinder Bajwa (B.B.), Kristin Elizabeth Low (K.E.L.), Jeffrey P. Tingley (J.P.T.), and D. Wade Abbott (D.W.A.).

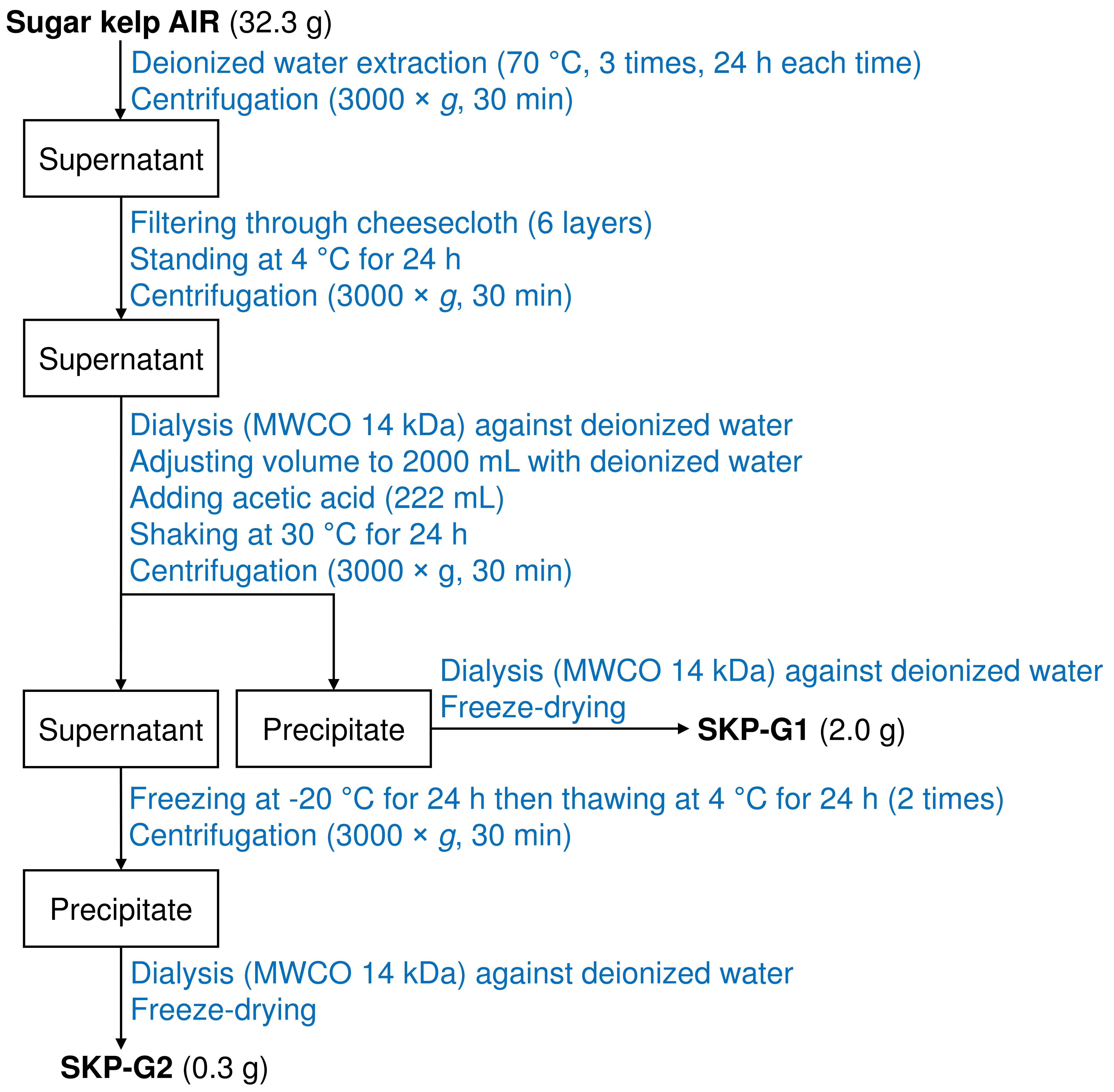

**Scheme S1.** Flow diagram showing the steps for isolating SKP-G1 and SKP-G2 from sugar kelp AIR.

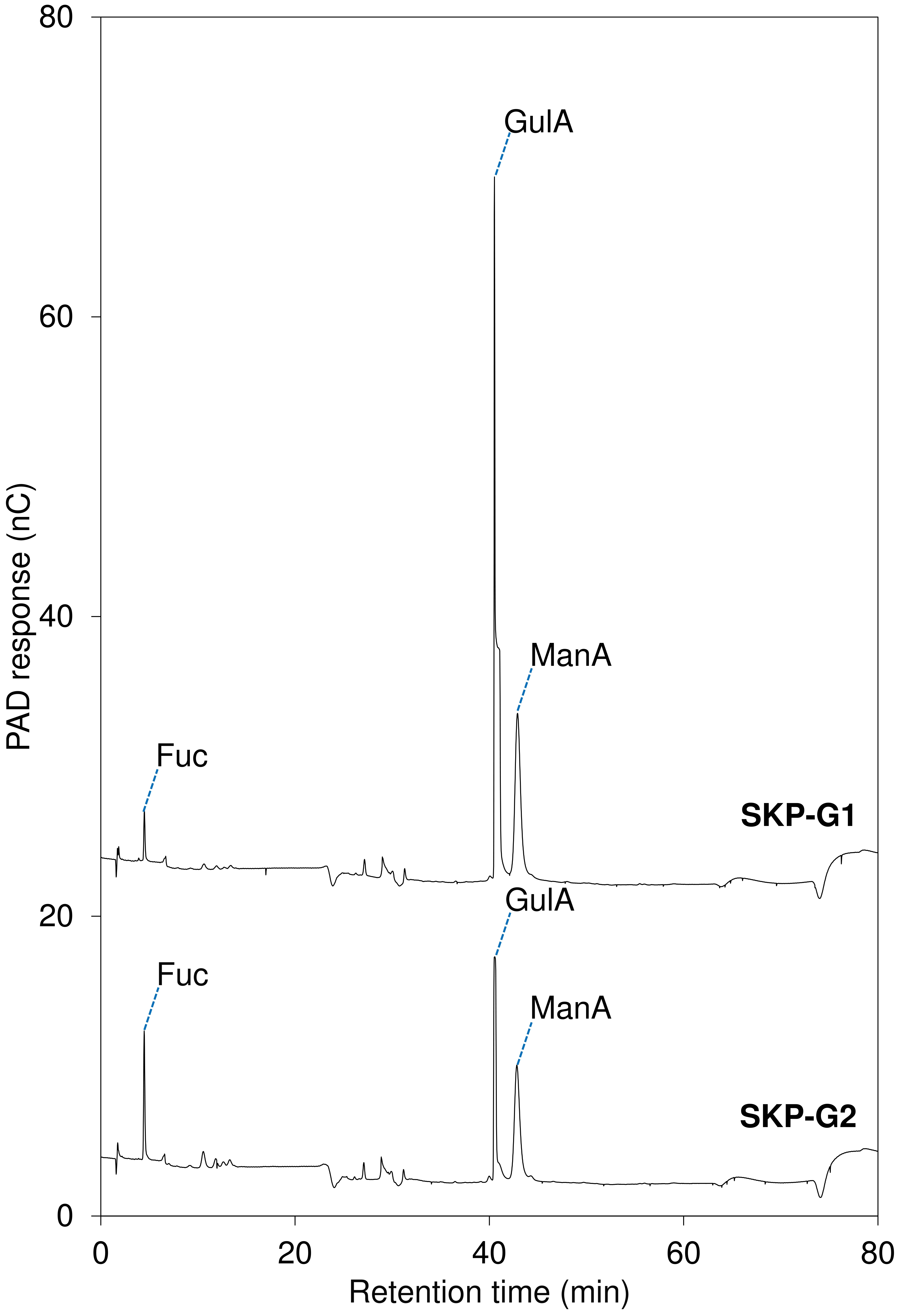

**Fig. S1.** HPAEC-PAD chromatograms of monosaccharides released from SKP-G1 (upper) and SKP-G2 (lower) by hydrolysis with 2 M TFA at 120 °C for 2 h.

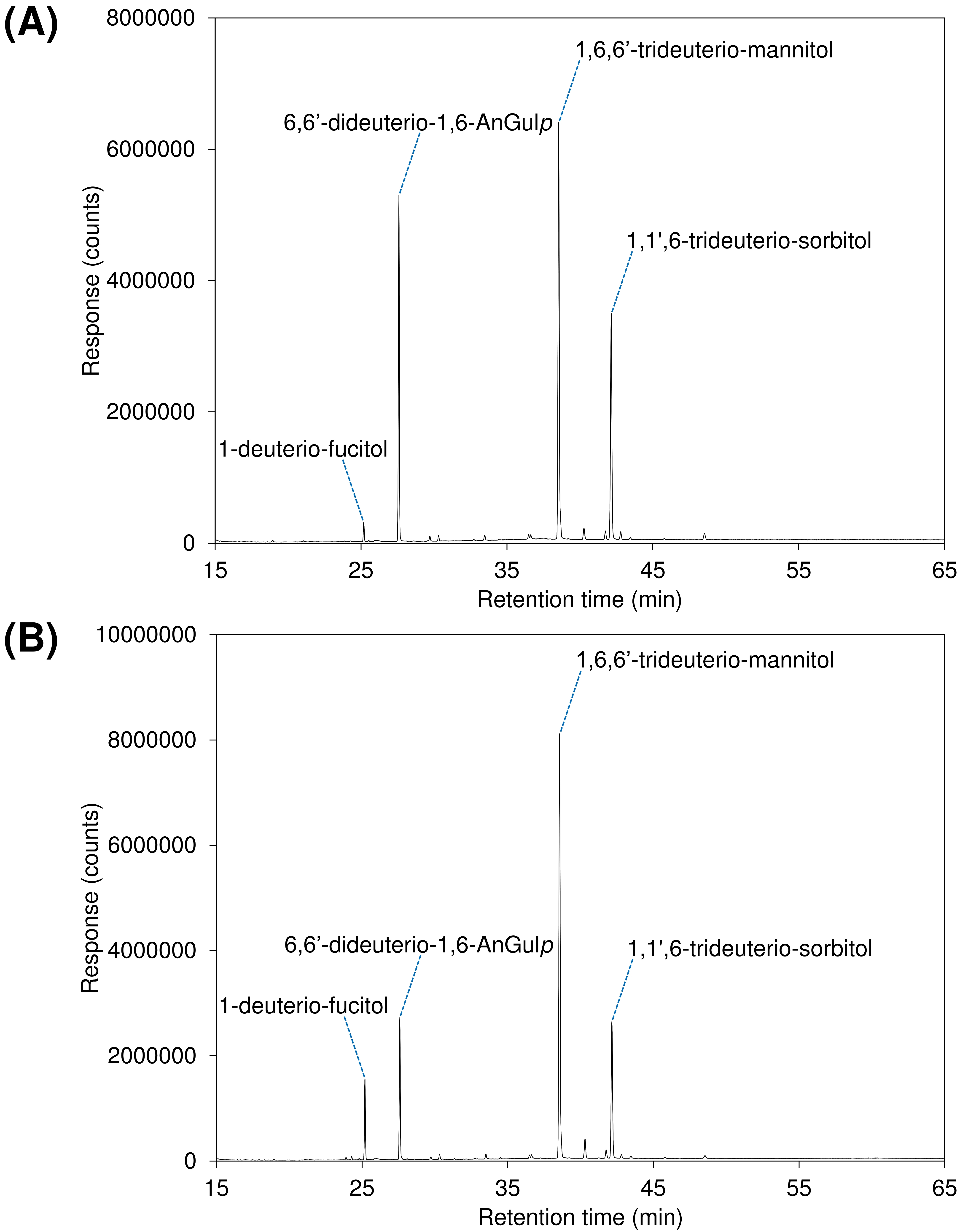

**Fig. S2.** Total ion current (TIC) chromatograms of acetate derivatives of monosaccharide alditols and 1,6-anhydrogulose from (A) SKP-G1 and (B) SKP-G2.

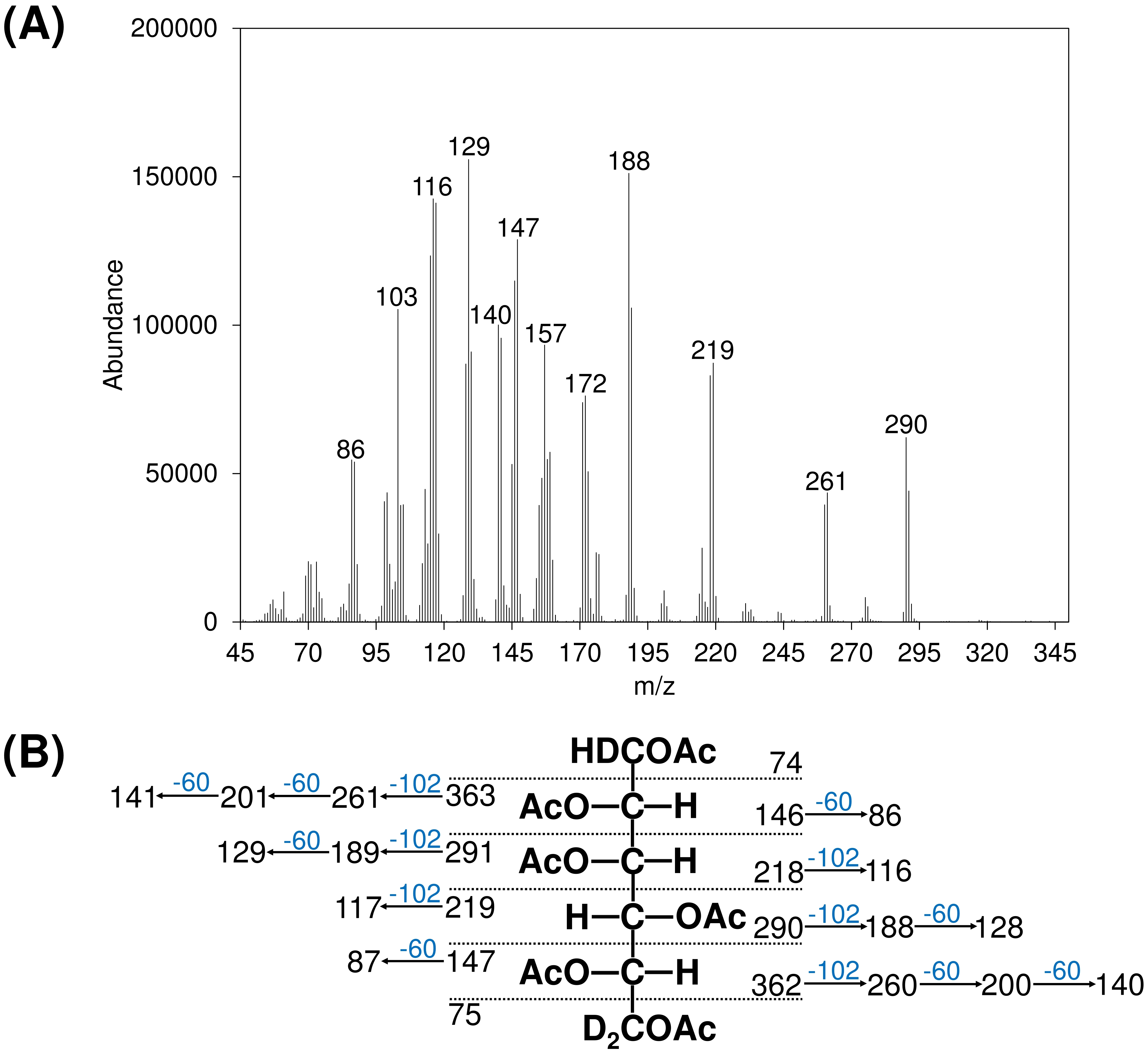

**Fig. S3.** (A) EI-MS spectrum and (B) assigned fragmentation pattern of the 1,1′,6-trideutero-sorbitol acetate derivative derived from GulA.

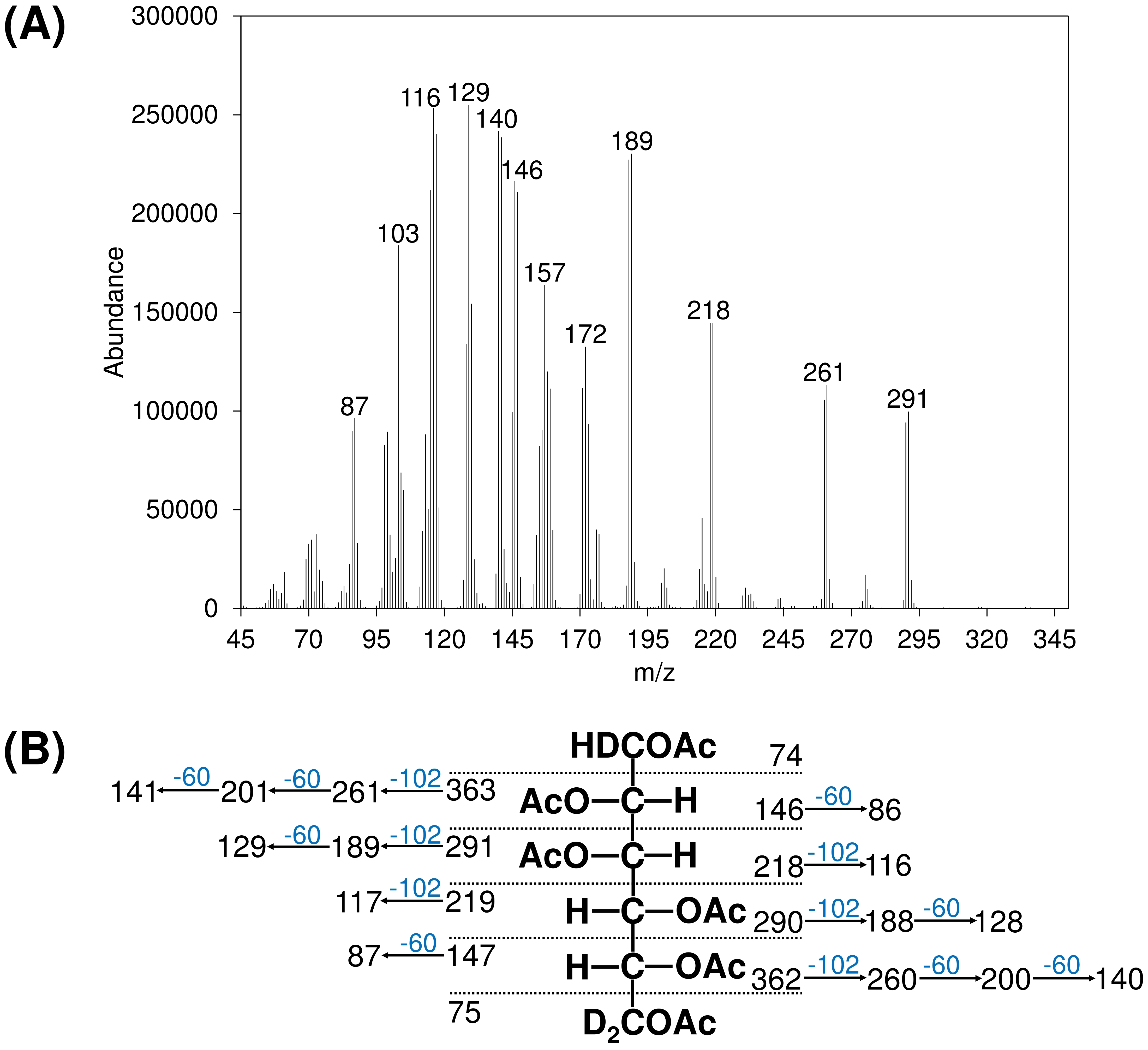

**Fig. S4.** (A) EI-MS spectrum and (B) assigned fragmentation pattern of the 1,6,6′-trideutero-mannitol acetate derivative derived from ManA.

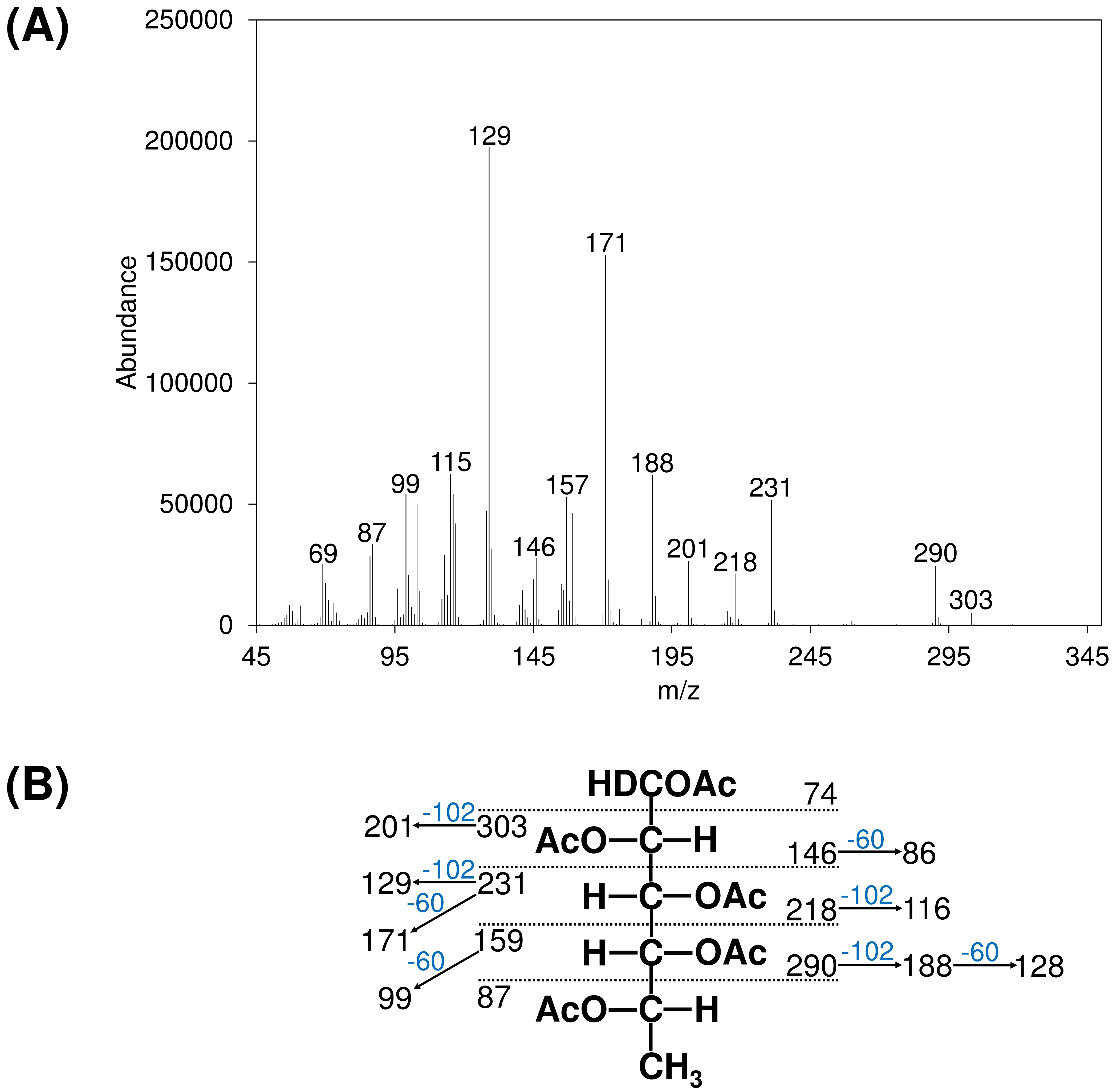

**Fig. S5.** (A) EI-MS spectrum and (B) assigned fragmentation pattern of the 1-deutero-fucitol acetate derivative derived from fucose.

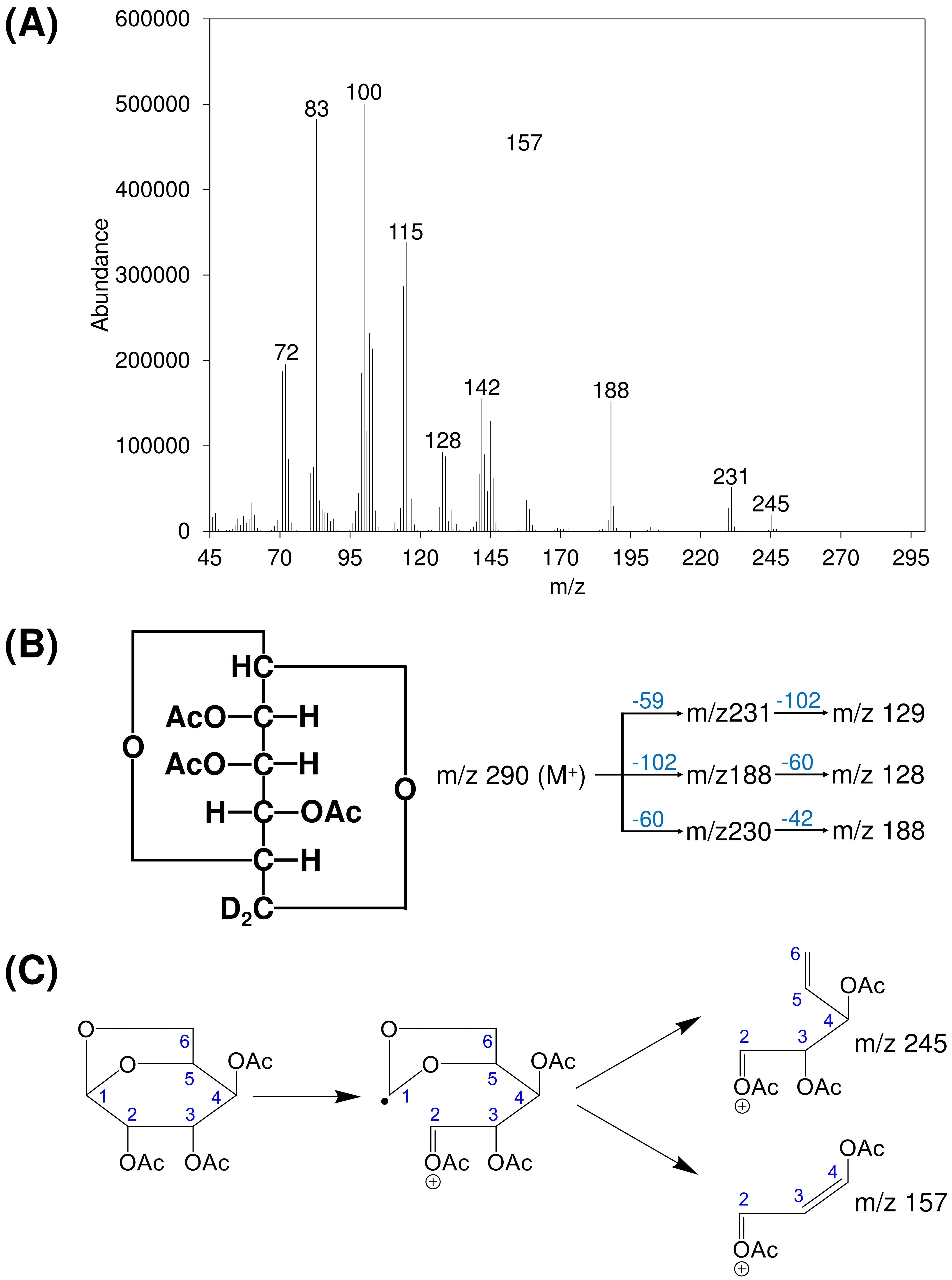

**Fig. S6.** (A) EI-MS spectrum and (B, C) assigned fragmentation patterns of the acetate derivative of 1,6-anhydro-Gul*p*, derived from GulA.

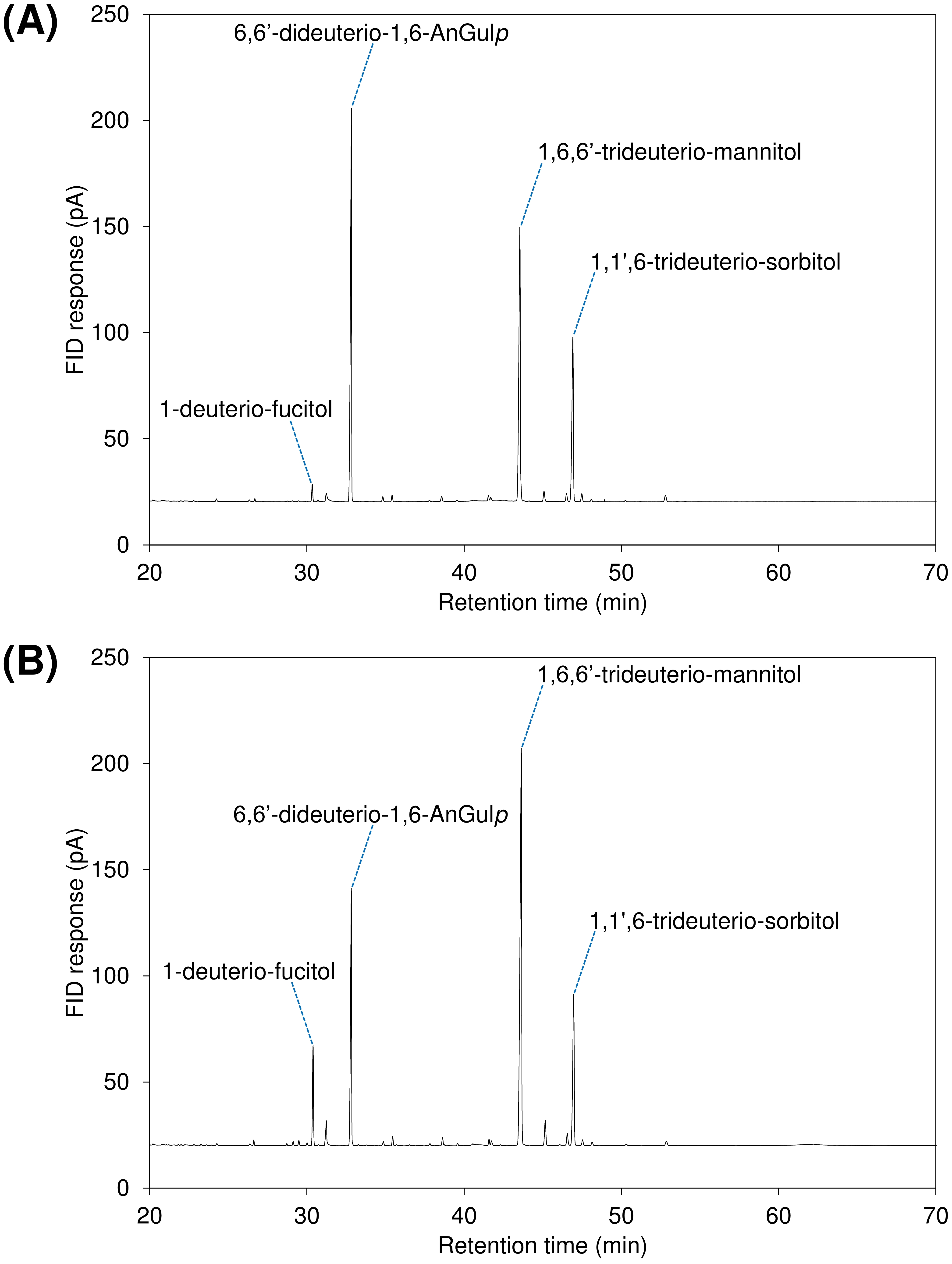

**Fig. S7**. Flame ionization detection (FID) chromatograms of acetate derivatives of monosaccharide alditols and 1,6-anhydrogulose from (A) SKP-G1 and (B) SKP-G2.

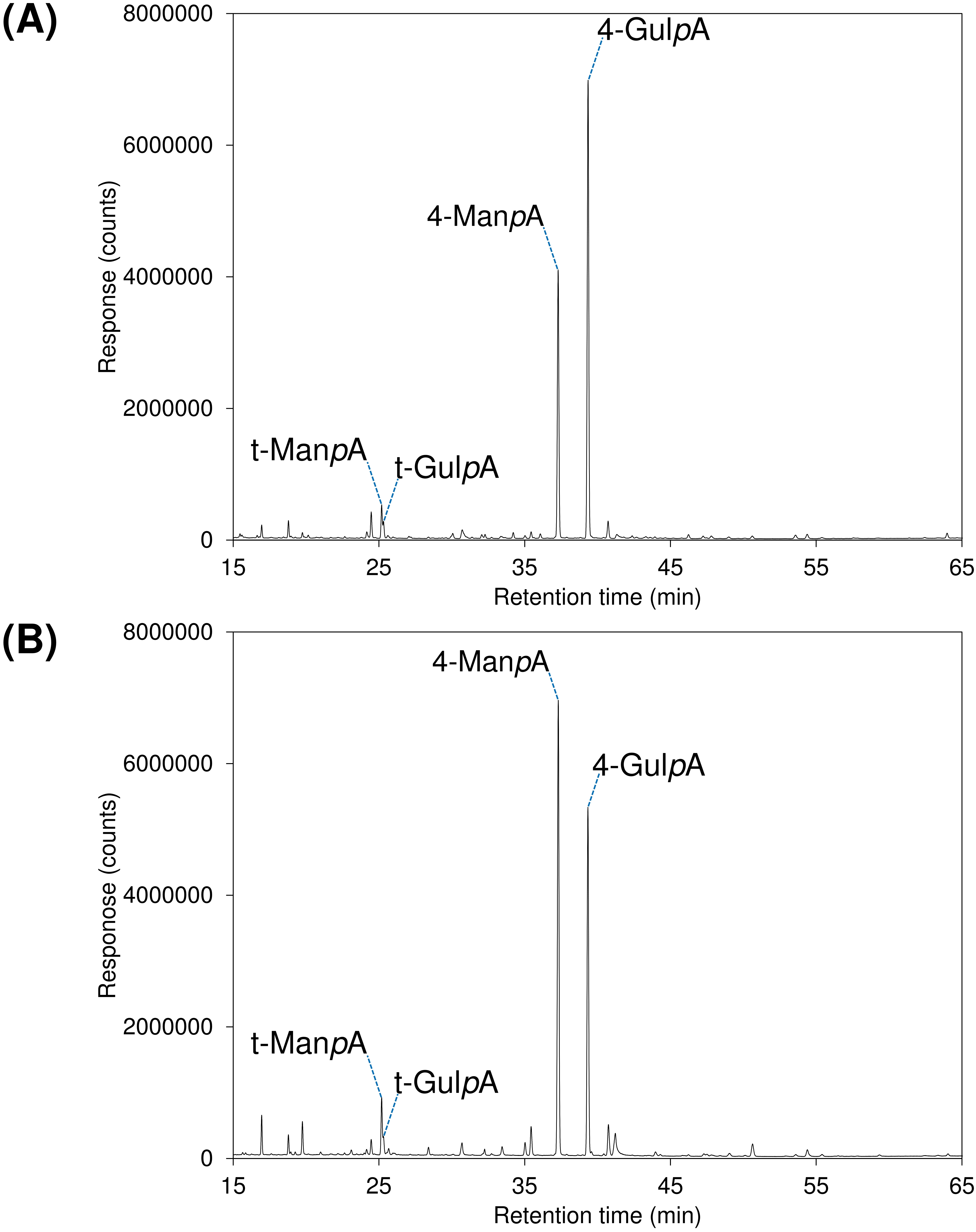

**Fig. S8**. Total ion current (TIC) chromatograms of partially methylated alditol acetates (PMAAs) prepared from (A) SKP-G1 and (B) SKP-G2.

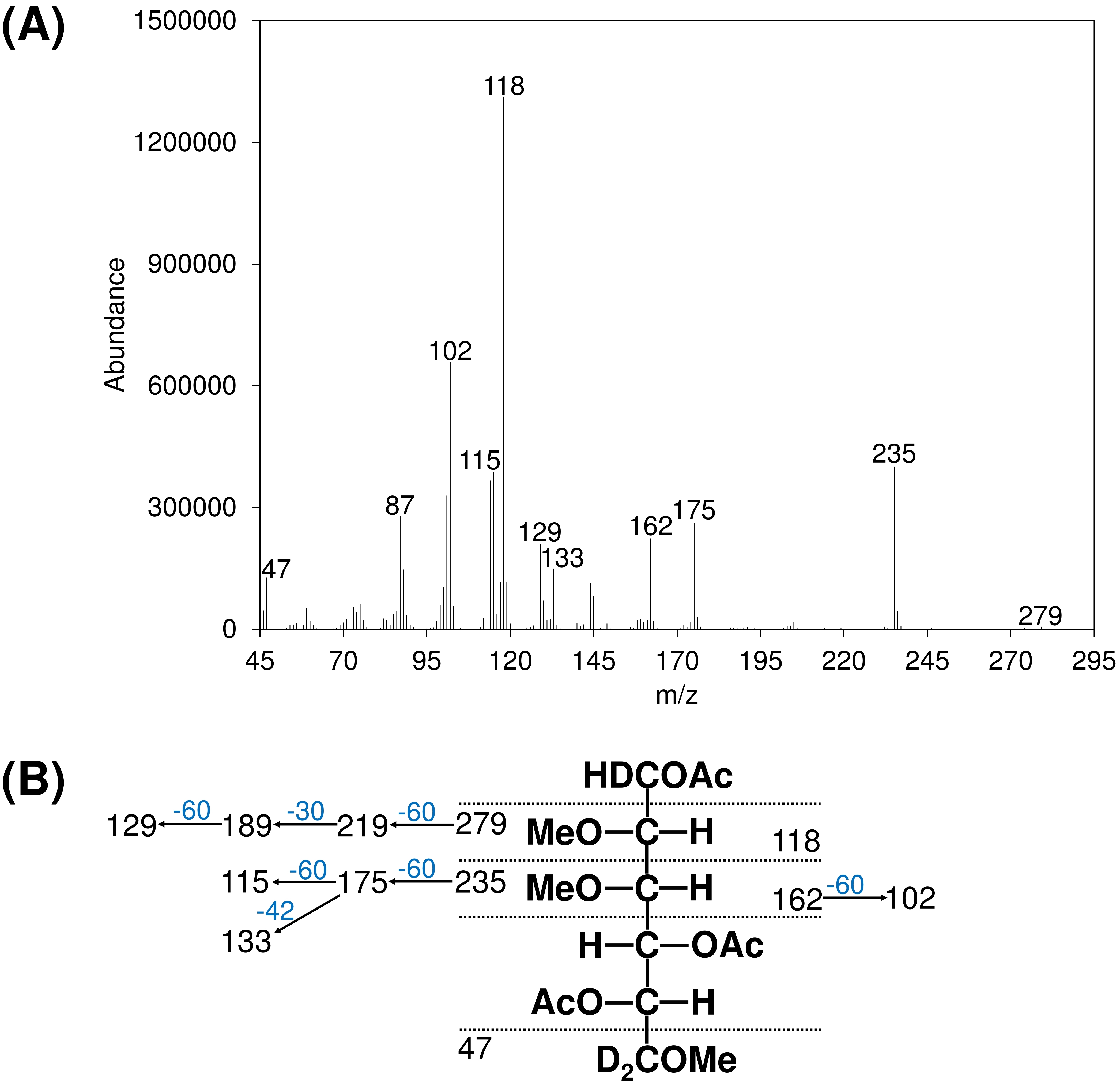

**Fig. S9**. (A) EI-MS spectrum and (B) assigned fragmentation pattern of the PMAA derived from 4-Gul*p*A.

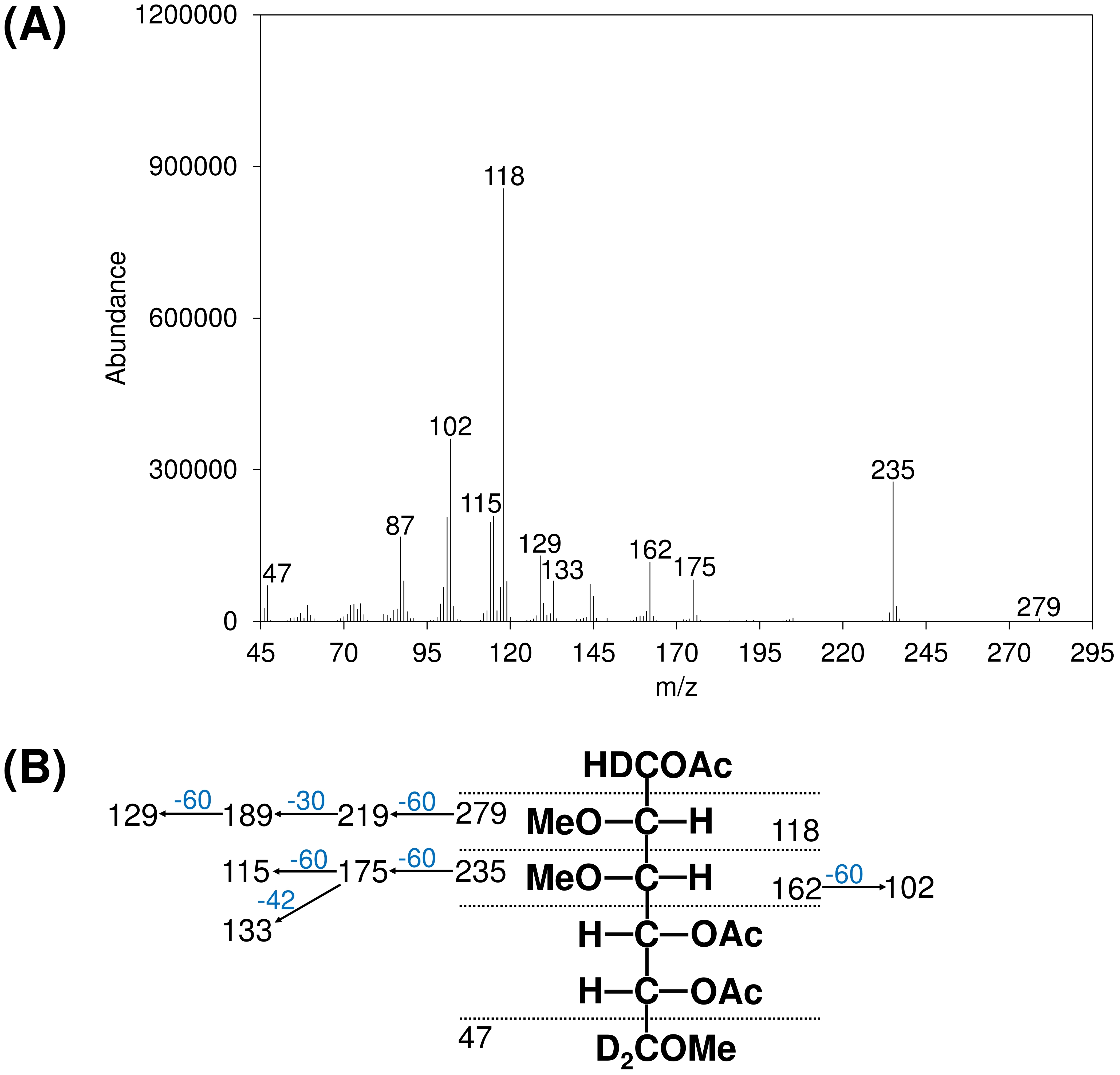

**Fig. S10.** (A) EI-MS spectrum and (B) assigned fragmentation pattern of the PMAA derived from 4-Man*p*A.

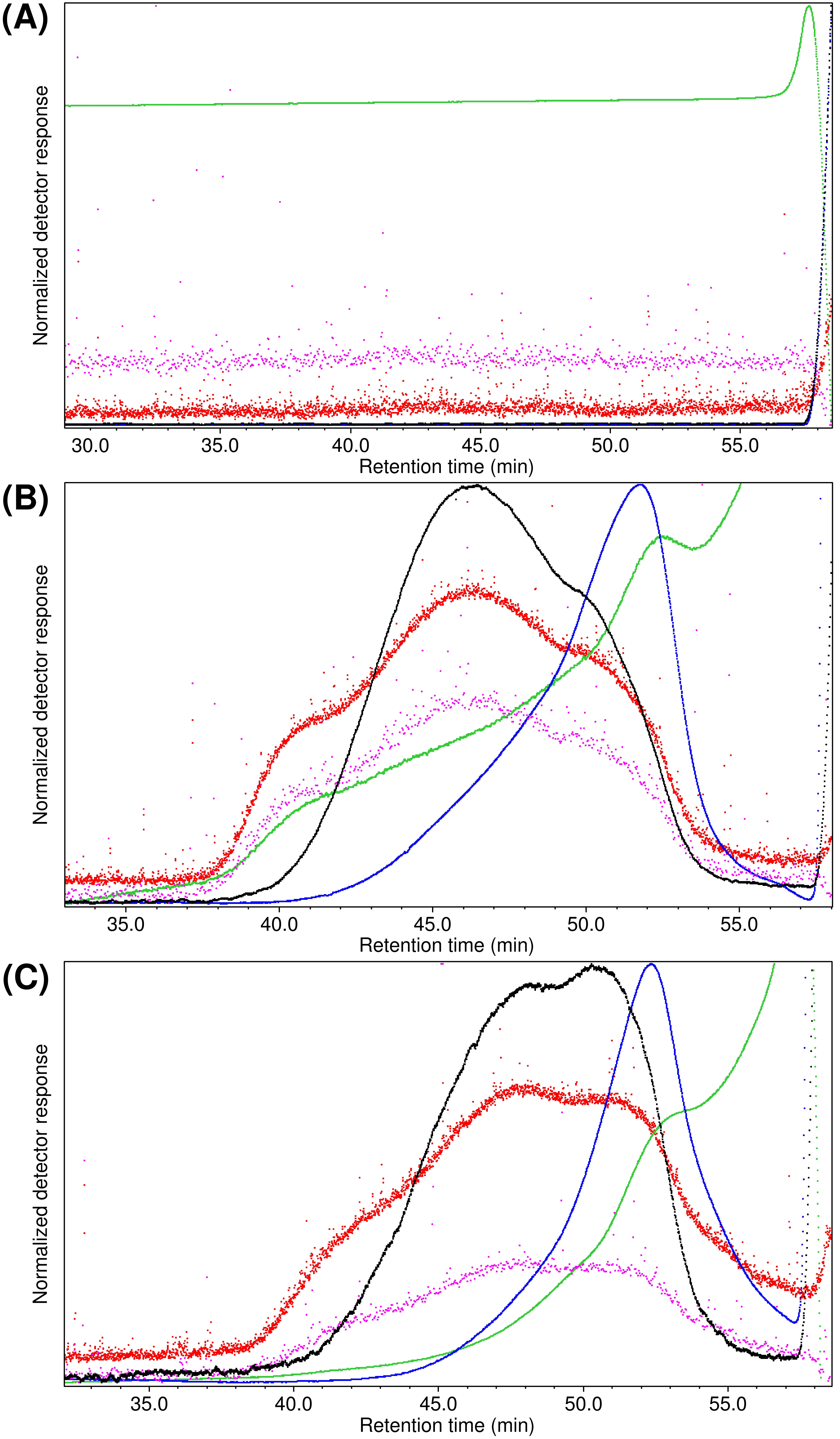

**Fig. S11.** Main elution region of the HPSEC chromatograms monitored by RI (blue), UV at 280 nm (green), viscometer DP signal (black), static light scattering at 90° (red), and dynamic light scattering at 90° (pink) following injection of (A) 0.1 M phosphate buffer (pH 7), (B) SKP-G1 dissolved in the buffer, and (C) SKP-G2 dissolved in the buffer. Each detector signal was normalized to its maximum value within the selected chromatogram window.

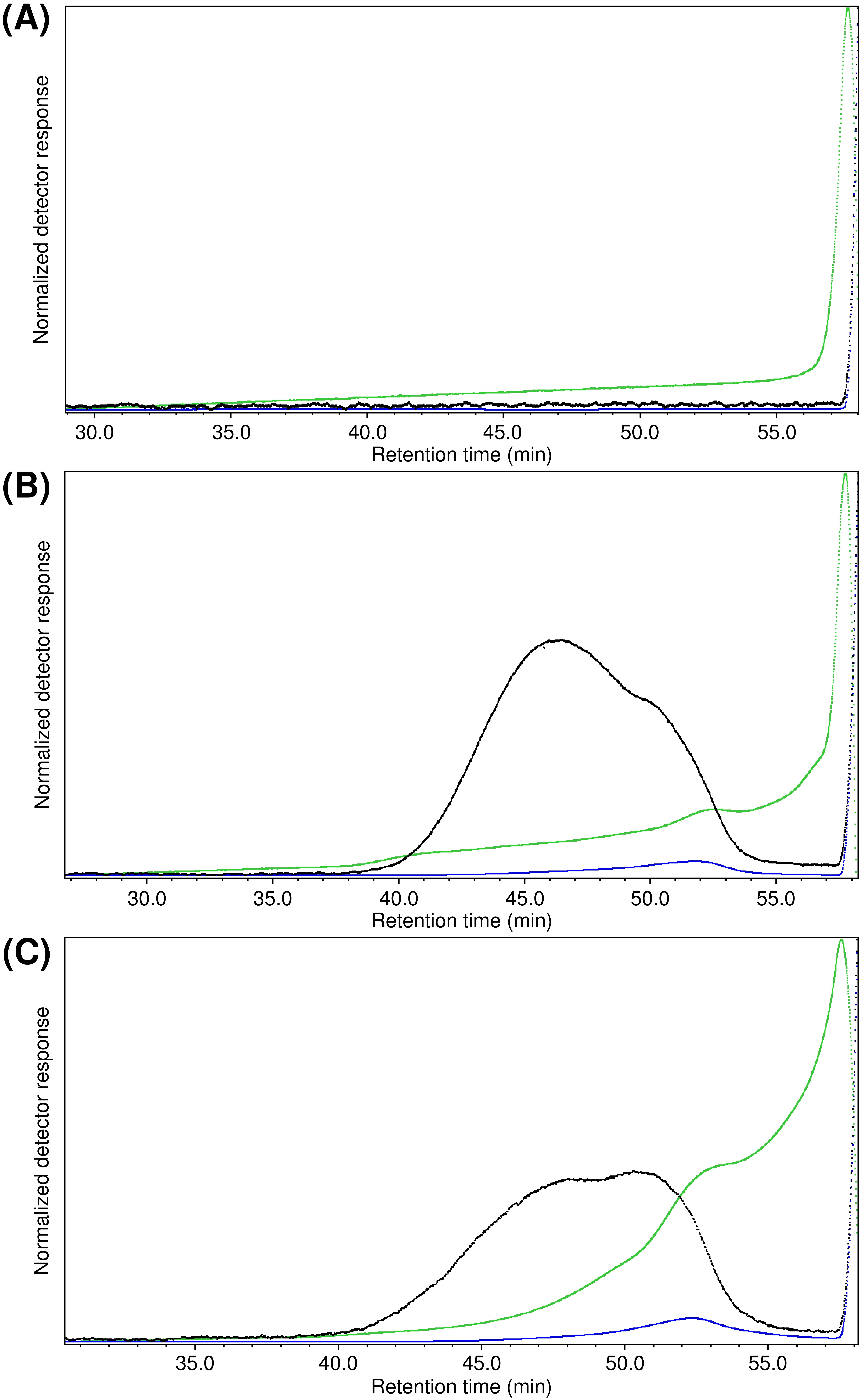

**Fig. S12.** Main elution region of the HPSEC chromatograms monitored by RI (blue), UV at 280 nm (green), and the viscometer DP signal (black) following injection of (A) 0.1 M phosphate buffer (pH 7), (B) SKP-G1 dissolved in the buffer, and (C) SKP-G2 dissolved in the buffer. Each detector signal was normalized to its maximum value within the selected chromatogram window.

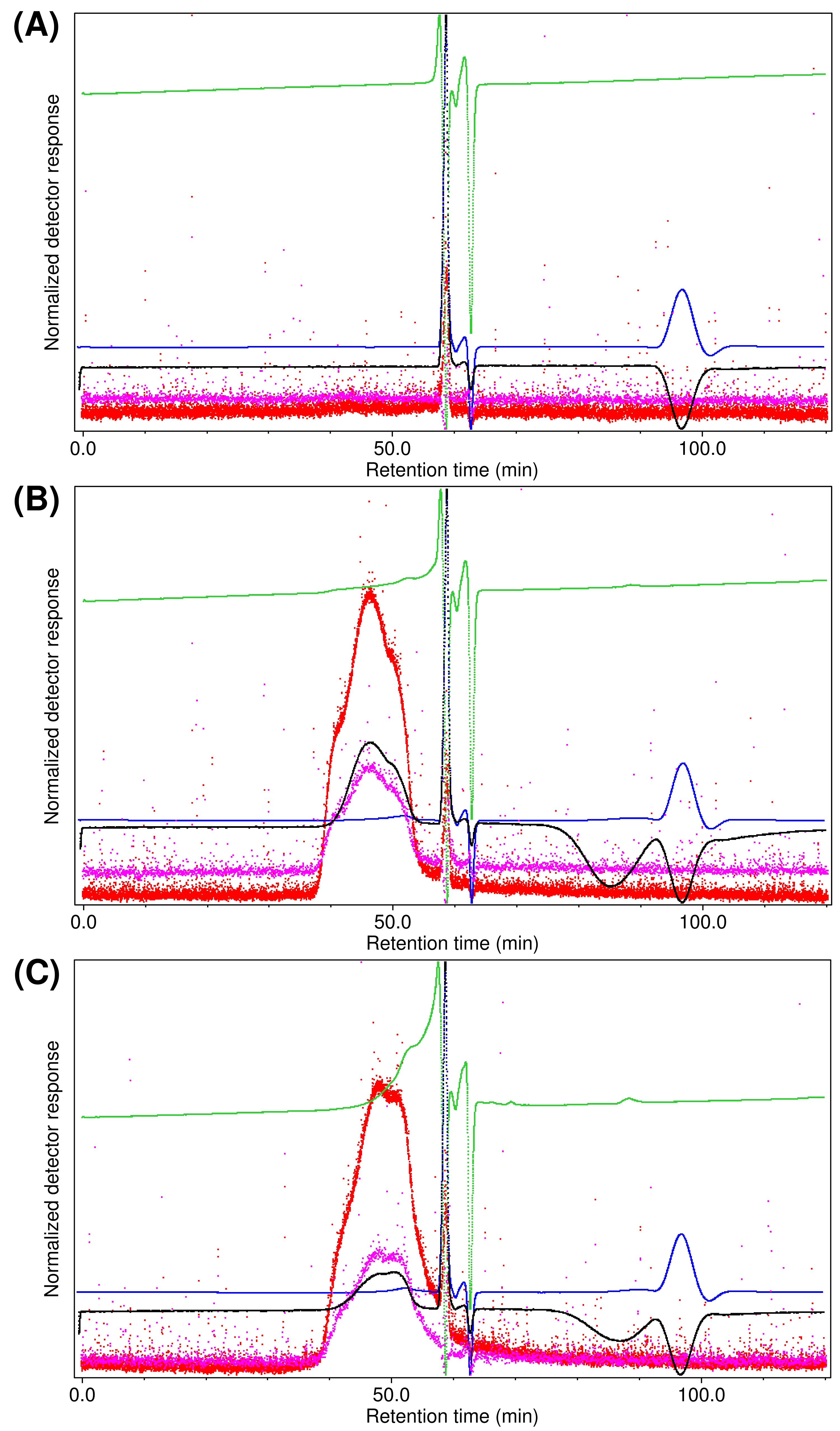

**Fig. S13.** Full HPSEC chromatograms (0–120 min) showing the main elution region and the post-void-volume region, corresponding to emptying of the viscometer delay column, following injections of (A) 0.1 M phosphate buffer (pH 7), (B) SKP-G1 dissolved in the buffer, and (C) SKP-G2 dissolved in the buffer. Chromatograms were monitored by RI (blue), UV at 280 nm (green), viscometer DP signal (black), static light scattering at 90° (red), and dynamic light scattering at 90° (pink). Each detector signal was normalized to its maximum value within the selected chromatogram window. In the post-void-volume region (64–120 min), residual sample in the delay column is flushed through the opposite side of the differential viscometer bridge from that used during the main elution, producing negative DP signals and positive RI signals that appear as delay (wash-out) peaks.
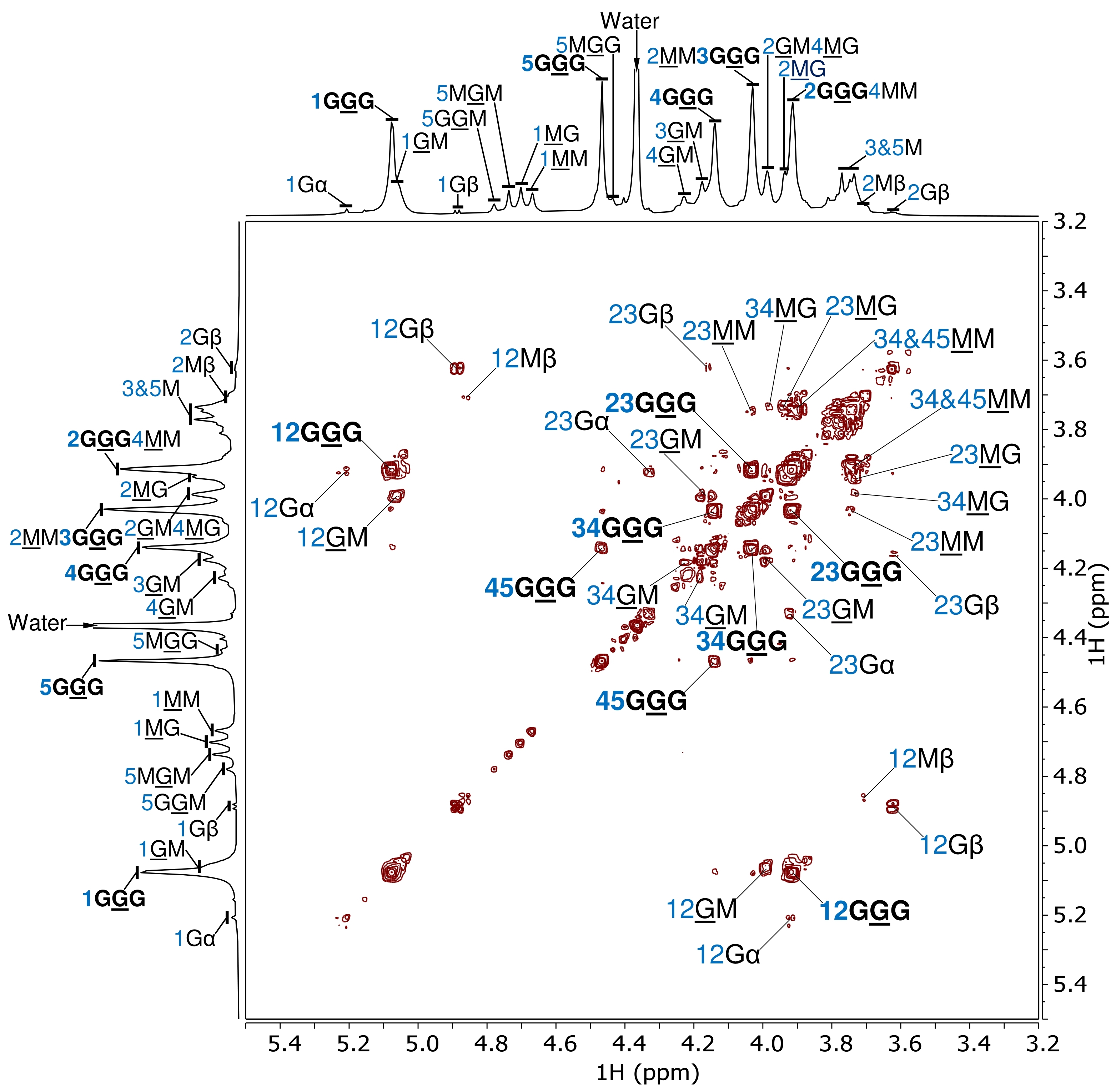

**Fig. S14.** Assignment of the 2D ^1^H–^1^H COSY NMR spectrum (700 MHz, 65 °C) of SKP-G1 in D_2_O. Residues G and M denote →4)-α-l-Gul*p*A-(1→ and →4)-β-d-Man*p*A-(1→, respectively, whereas the minor residues Gα, Gβ, and Mβ represent the reducing end residues →4)-α-l-Gul*p*A, →4)-β-l-Gul*p*A, and →4)-β-d-Man*p*A, respectively. For example, the annotations “12GGG” and “12GM” indicate the cross-peaks between H-1 and H-2 of the underlined G residue in the GGG triad and GM diad, respectively, while “12Gα” indicates the cross-peak between H-1 and H-2 of residue Gα. In the stacked 1D ^1^H NMR spectrum, the annotation “1GGG” denotes H-1 of the underlined central G residue in the GGG triad, while “3GGG2MM” indicates overlapping signals of “3GGG” and “2MM”. Annotations for the dominant GGG triad are highlighted in bold.

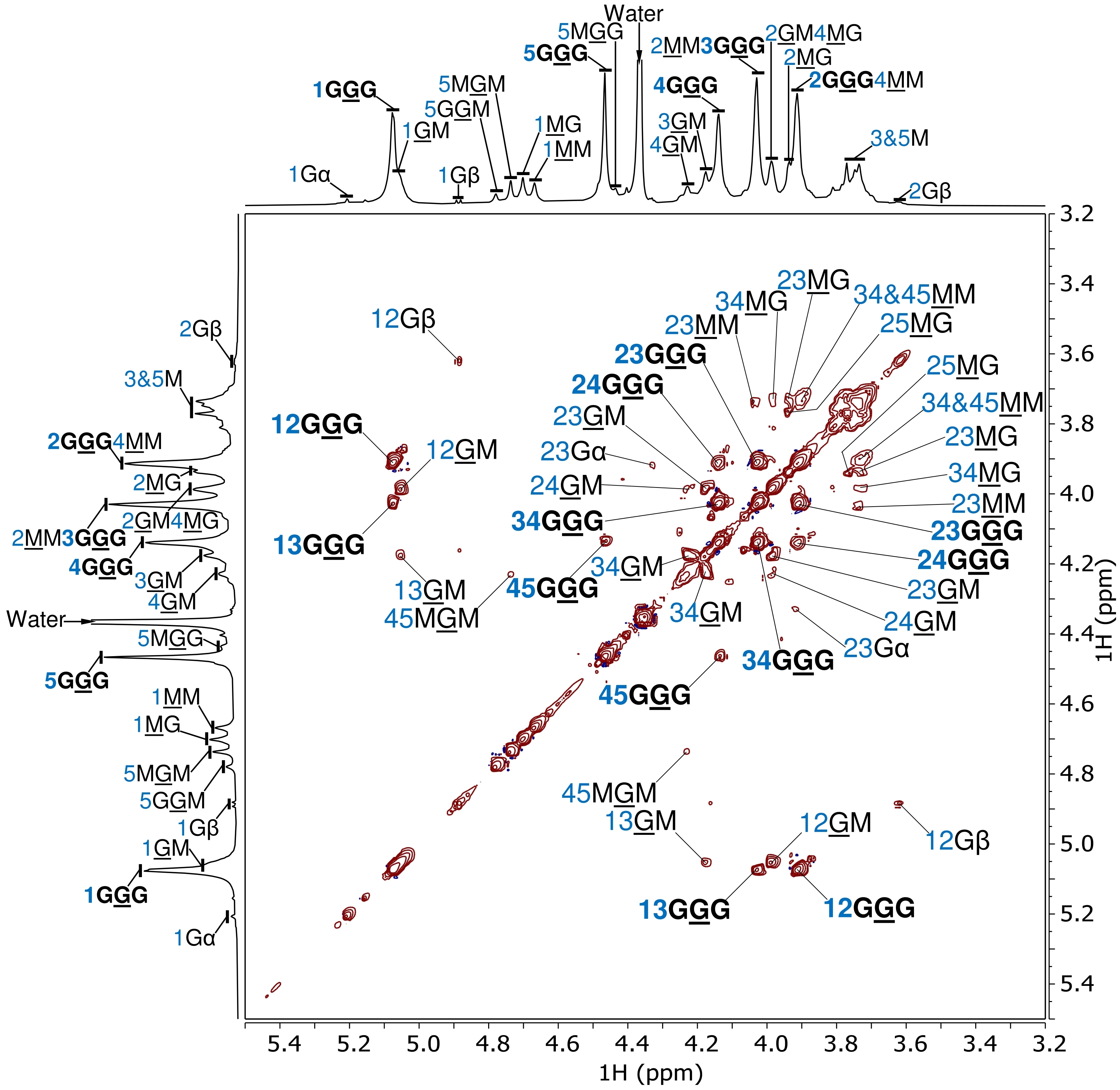

**Fig. S15.** Assignment of the 2D ^1^H–^1^H TOCSY NMR spectrum (700 MHz, 65 °C) of SKP-G1 in D_2_O. Residues G and M denote →4)-α-l-Gul*p*A-(1→ and →4)-β-d-Man*p*A-(1→, respectively, whereas the minor residues Gα and Gβ represent the reducing end residues →4)-α-l-Gul*p*A and →4)-β-l-Gul*p*A, respectively. For example, the annotations “12GGG” and “12GM” indicate the cross-peaks between H-1 and H-2 of the underlined G residue in the GGG triad and GM diad, respectively, while “12Gβ” indicates the cross-peak between H-1 and H-2 of residue Gβ. In the stacked 1D ^1^H NMR spectrum, the annotation “1GGG” denotes H-1 of the underlined central G residue in the GGG triad, while “3GGG2MM” indicates overlapping signals of “3GGG” and “2MM”. Annotations for the dominant GGG triad are highlighted in bold.

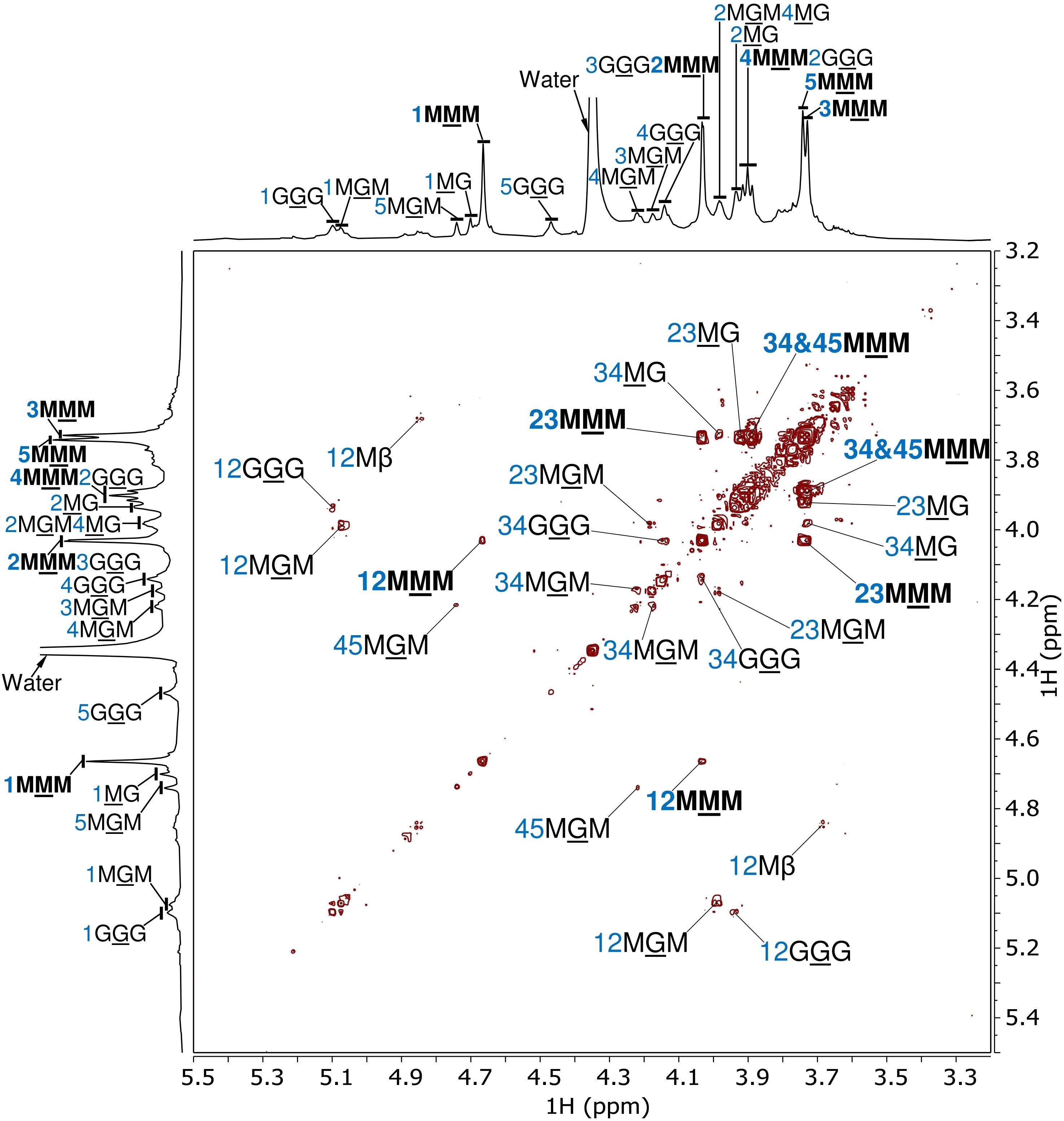

**Fig. S16.** Assignment of the 2D ^1^H–^1^H COSY NMR spectrum (700 MHz, 65 °C) of SKP-G2 in D_2_O. Residues G and M denote →4)-α-l-Gul*p*A-(1→ and →4)-β-d-Man*p*A-(1→, respectively. For example, the annotations “12MMM” and “12GGG” indicate the cross-peaks between H-1 and H-2 of the underlined central M and G residues in the MMM and GGG triads, respectively. In the stacked 1D ^1^H NMR spectrum, the annotation “1MMM” denotes H-1 of the underlined central M residue in the MMM triad, whereas “4MMM2GGG” indicates overlapping signals of “4MMM” and “2GGG”. Annotations for the dominant MMM triad are highlighted in bold.

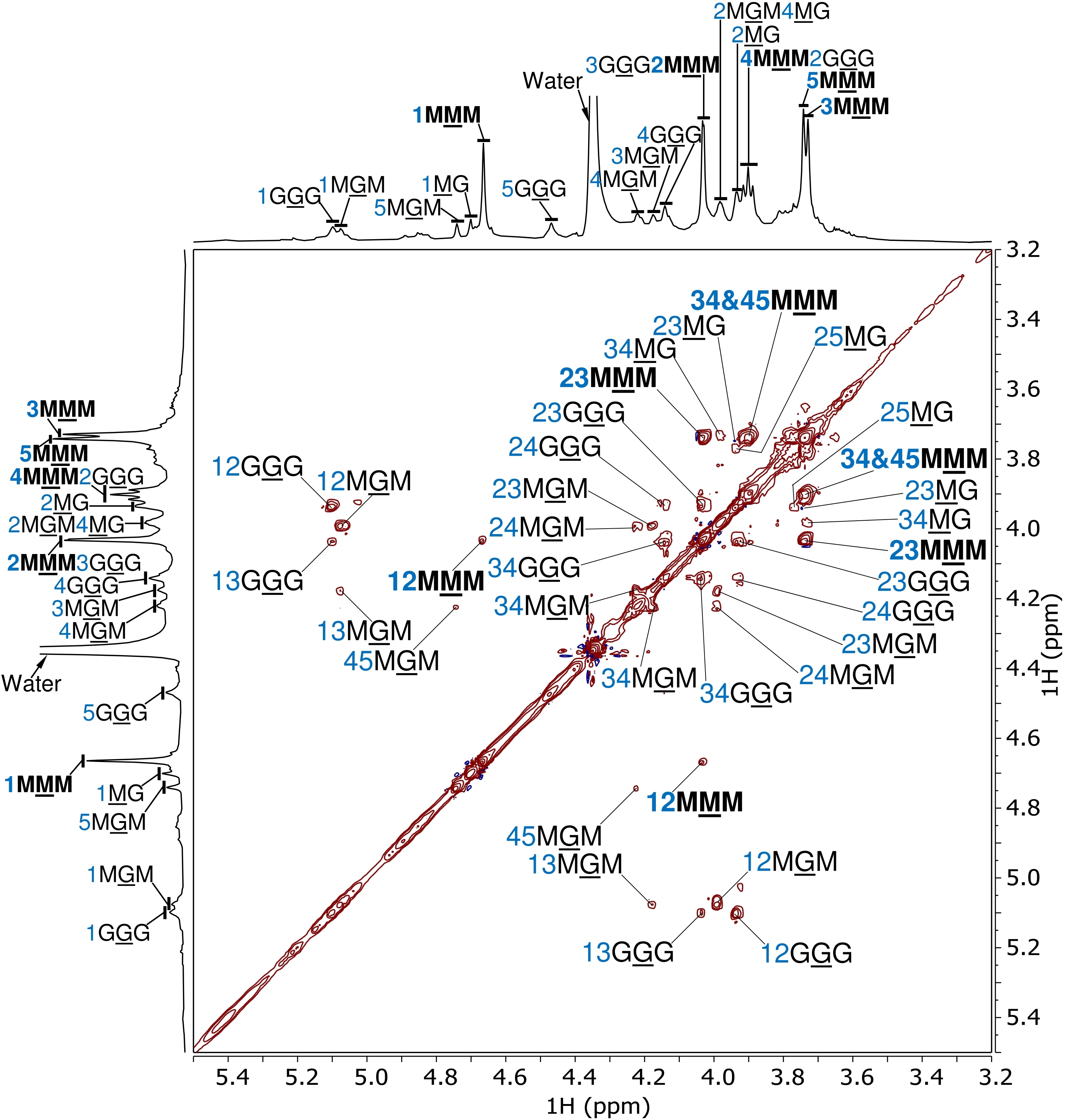

**Fig. S17.** Assignment of the 2D ^1^H–^1^H TOCSY NMR spectrum (700 MHz, 65 °C) of SKP-G2 in D_2_O. Residues G and M denote →4)-α-l-Gul*p*A-(1→ and →4)-β-d-Man*p*A-(1→, respectively. For example, the annotations “12MMM” and “12GGG” indicate the cross-peaks between H-1 and H-2 of the underlined central M and G residues in the MMM and GGG triads, respectively. In the stacked 1D ^1^H NMR spectrum, the annotation “1MMM” denotes H-1 of the underlined central M residue in the MMM triad, whereas “4MMM2GGG” indicates overlapping signals of “4MMM” and “2GGG”. Annotations for the dominant MMM triad are highlighted in bold.

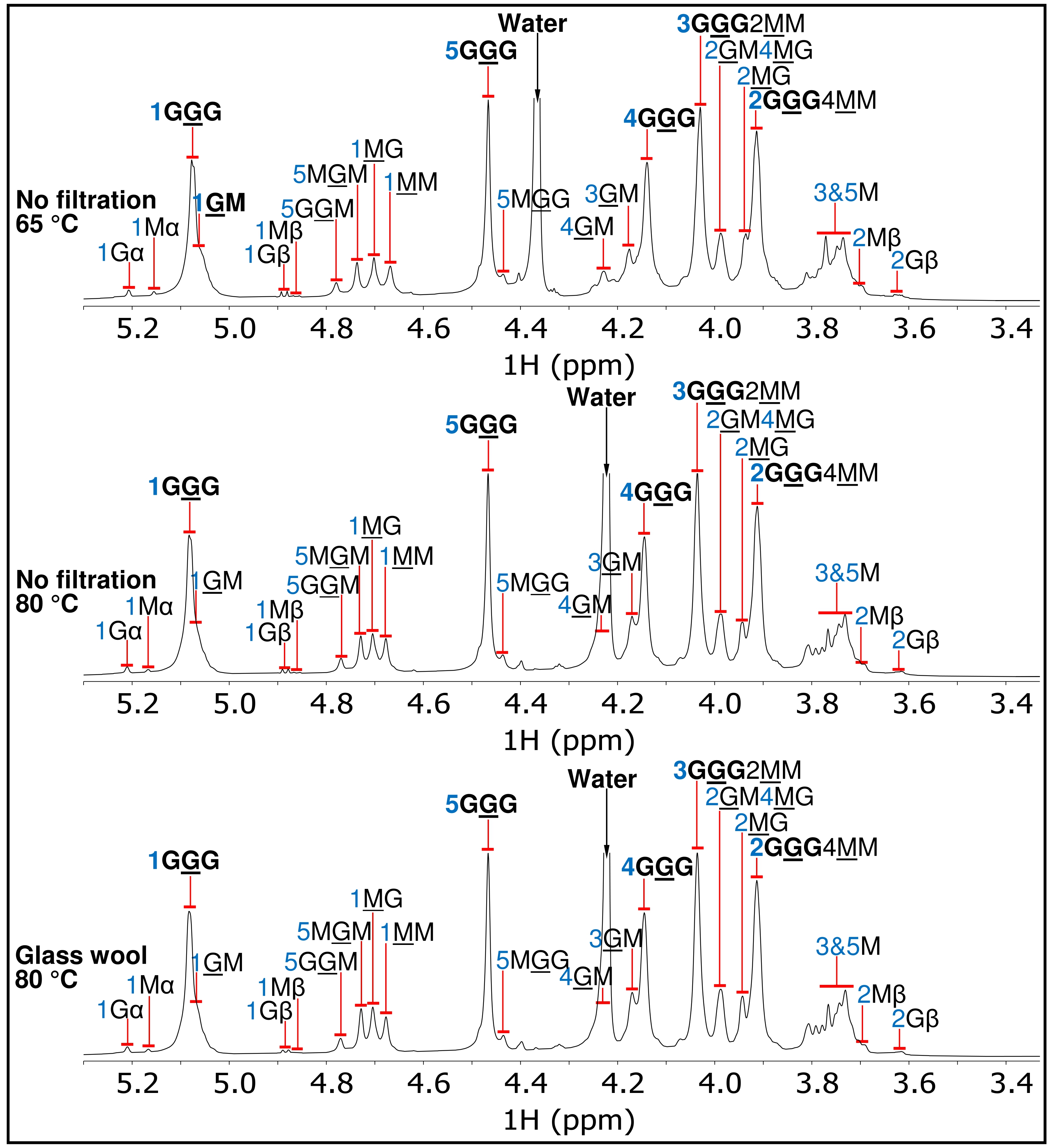

**Fig. S18.** Assignment of the 1D ^1^H NMR spectra of SKP-G1 in D_2_O collected at 65 °C without filtration (upper), at 80 °C without filtration (middle), and at 80 °C with glass wool filtration (lower). For example, the annotation “1GGG” denotes H-1 of the underlined central G residue in the GGG triad, whereas “3GGG2MM” indicates overlapping signals of “3GGG” and “2MM”. Annotations for the dominant GGG triad are highlighted in bold.

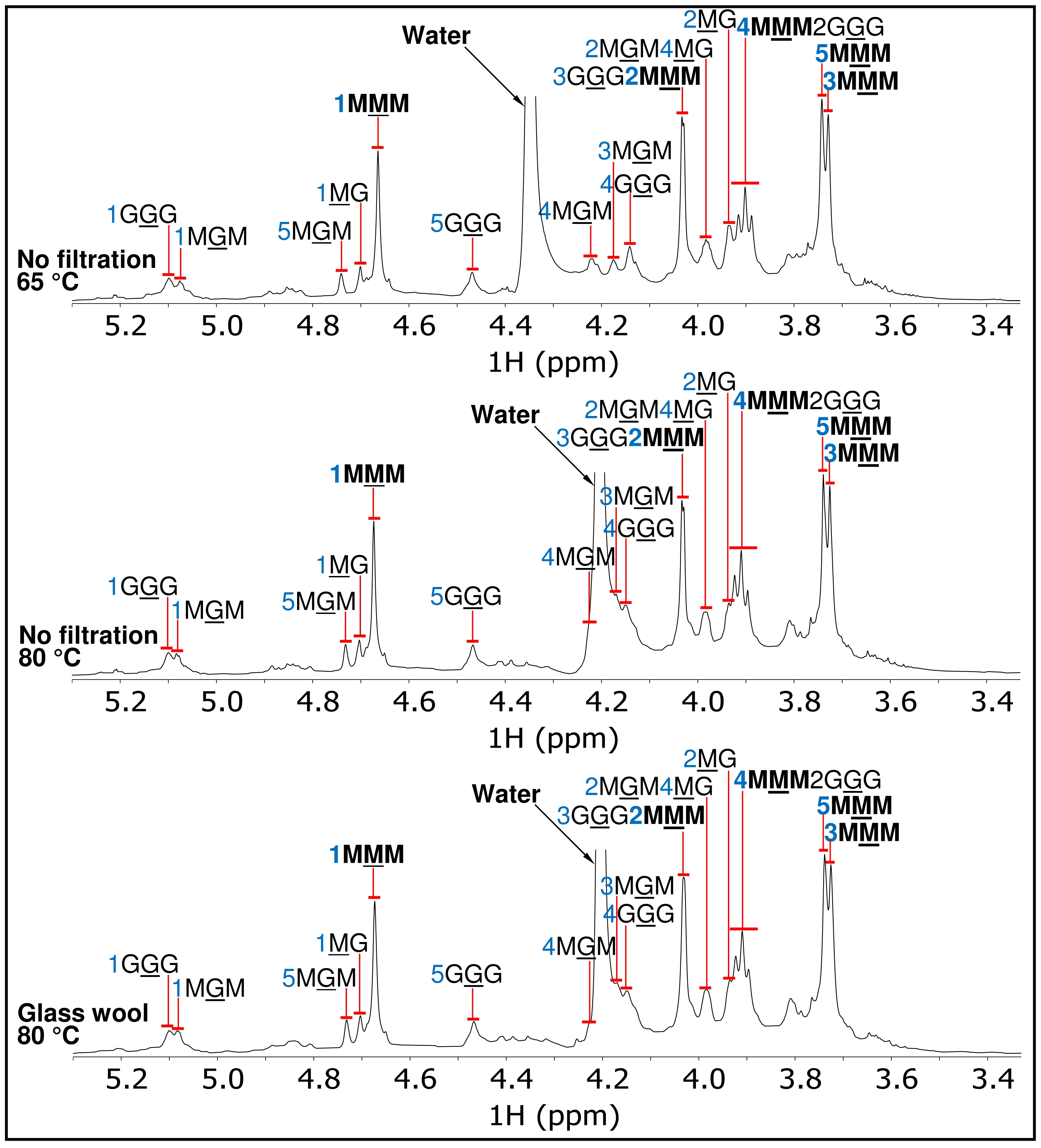

**Fig. S19.** Assignment of the 1D ^1^H NMR spectra of SKP-G2 in D_2_O collected at 65 °C without filtration (upper), at 80 °C without filtration (middle), and at 80 °C with glass wool filtration (lower). For example, the annotation “1MMM” denotes H-1 of the underlined central M residue in the MMM triad, whereas “4MMM2GGG” indicates overlapping signals of “4MMM” and “2GGG”. Annotations for the dominant MMM triad are highlighted in bold.

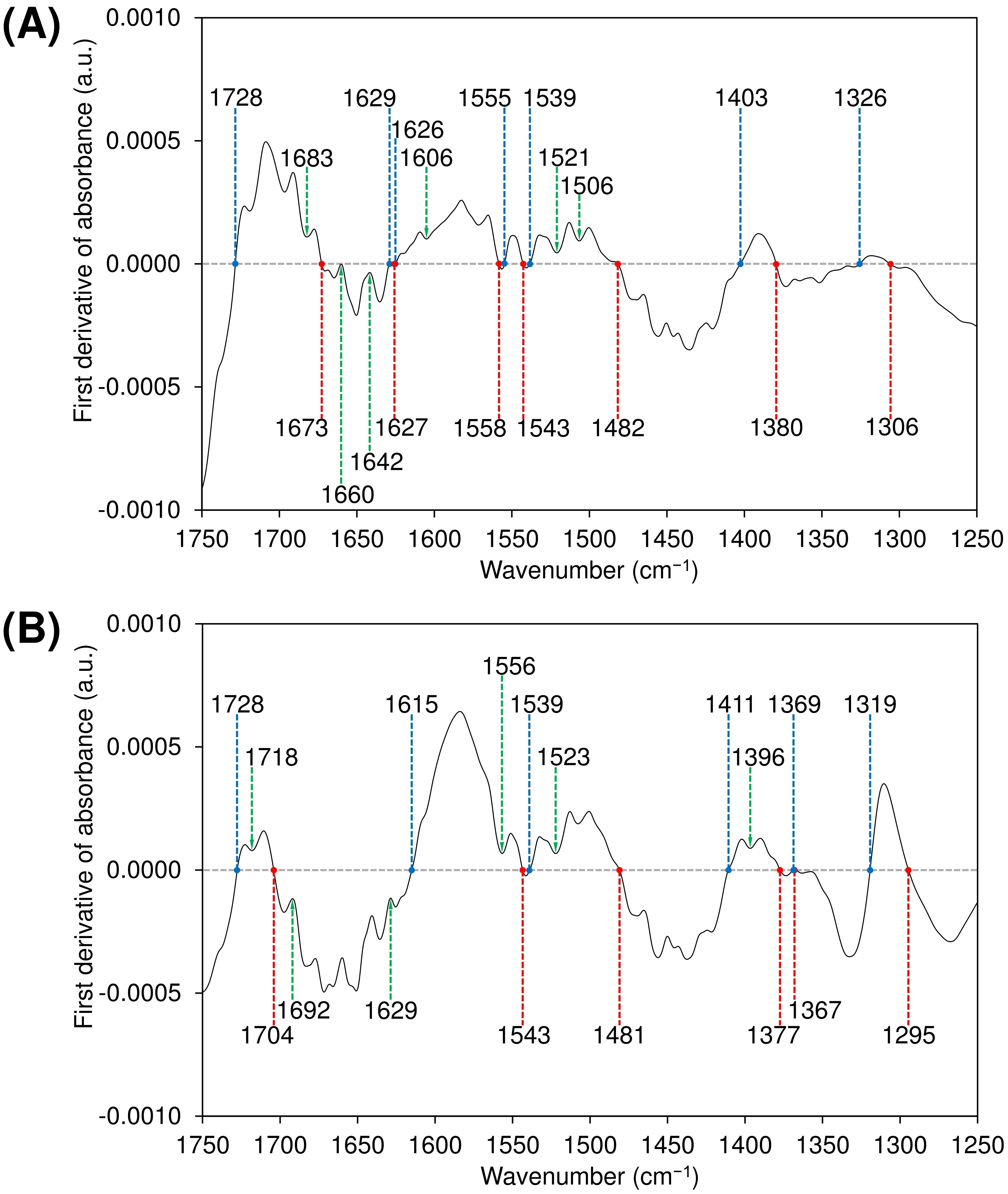

**Fig. S20.** First-derivative ATR-FTIR spectra of (A) SKP-G1 and (B) SKP-G2 in the 1250–1750 cm^–1^ region. Colored lines indicate key spectral features. The blue line indicates a zero crossing from positive to negative with increasing wavenumber, whereas the red line indicates a zero crossing in the opposite direction, from negative to positive. The green line marks the local extremum closest to zero.

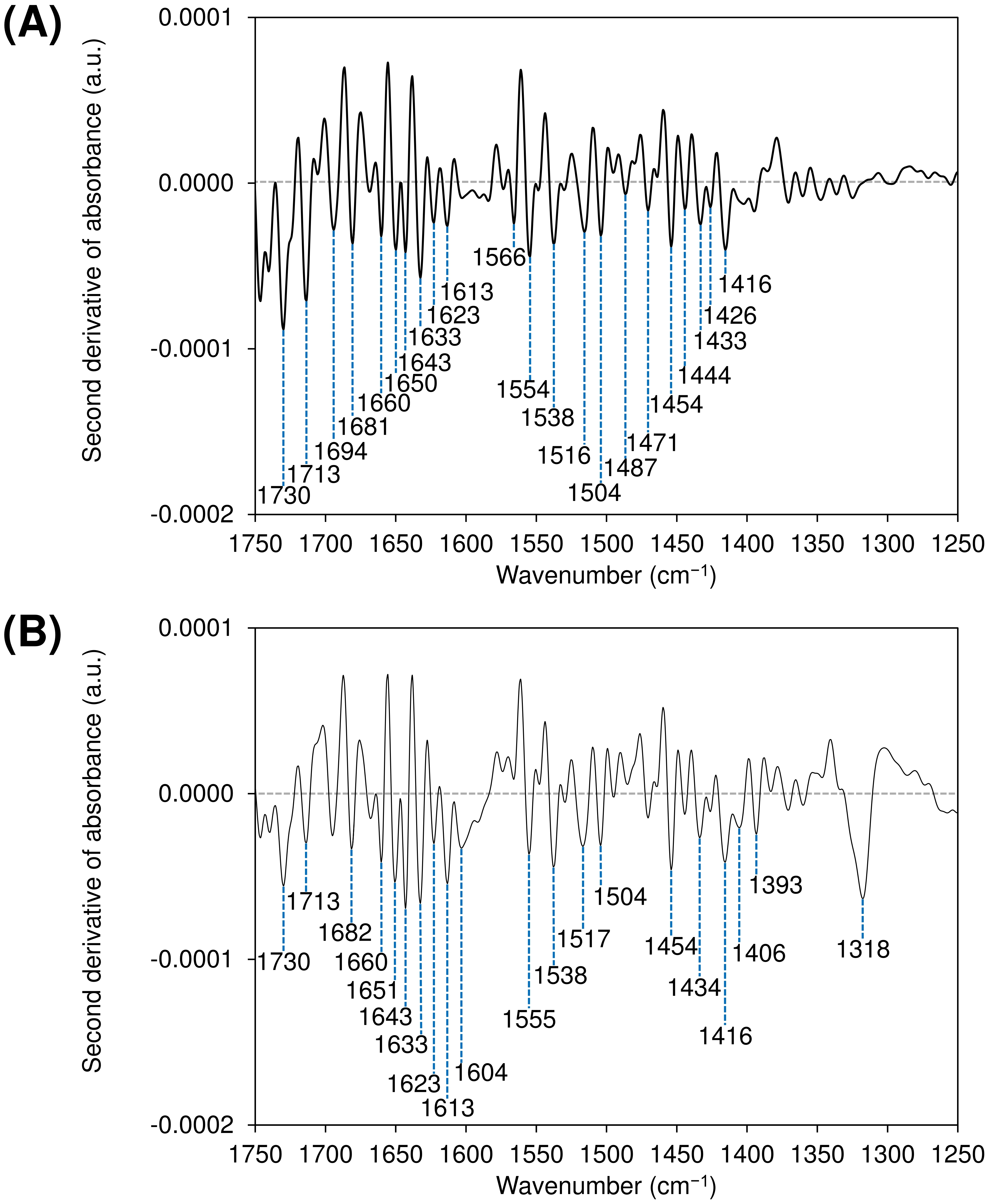

**Fig. S21.** Second-derivative ATR-FTIR spectra of (A) SKP-G1 and (B) SKP-G2 in the 1250–1750 cm⁻¹ region. The blue vertical lines indicate the wavenumbers of selected negative second-derivative bands.

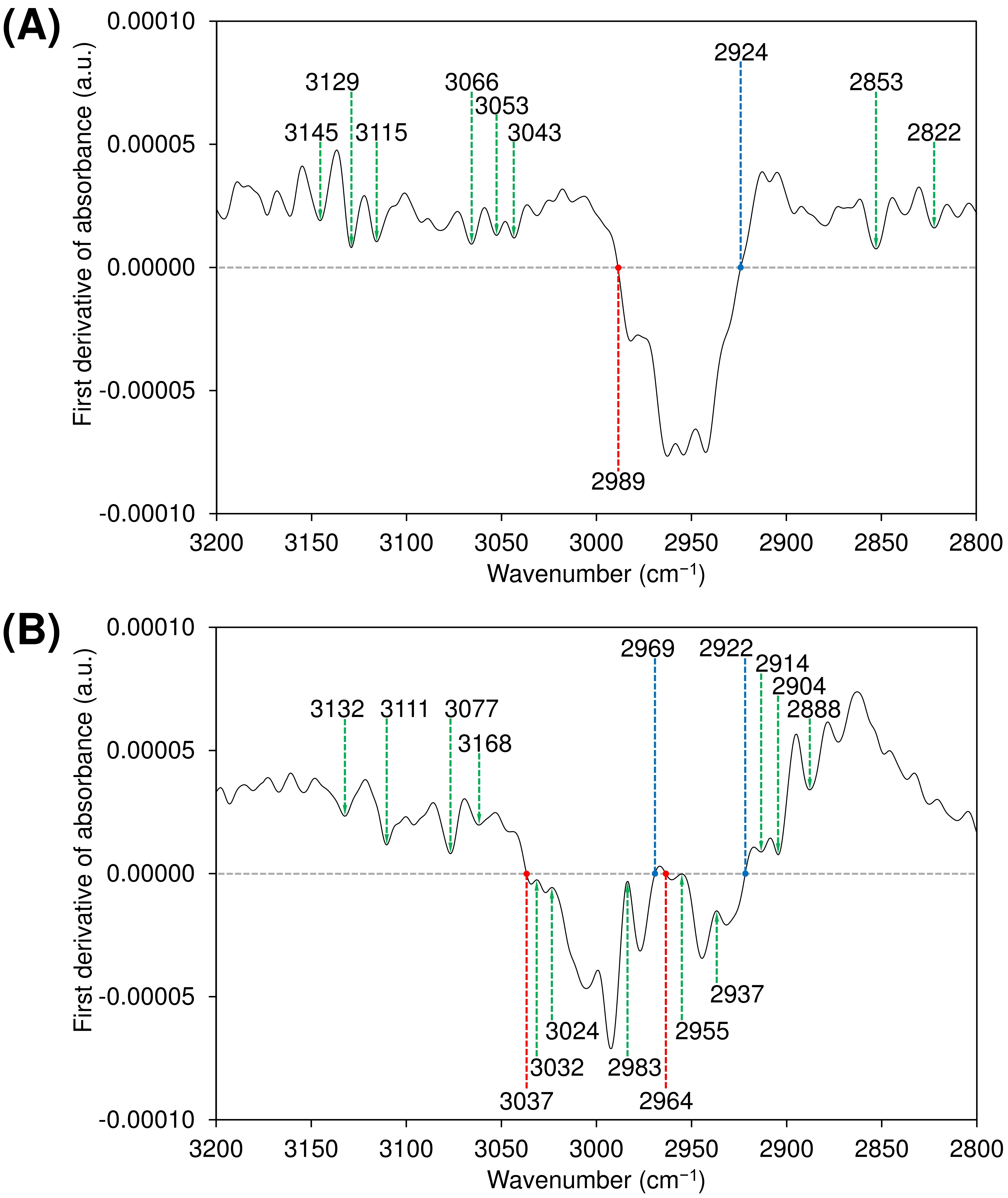

**Fig. S22.** First-derivative ATR-FTIR spectra of (A) SKP-G1 and (B) SKP-G2 in the 2800–3200 cm⁻¹ region. Colored lines indicate key spectral features. The blue line indicates a zero crossing from positive to negative with increasing wavenumber, whereas the red line indicates a zero crossing in the opposite direction, from negative to positive. The green line marks the local extremum closest to zero.

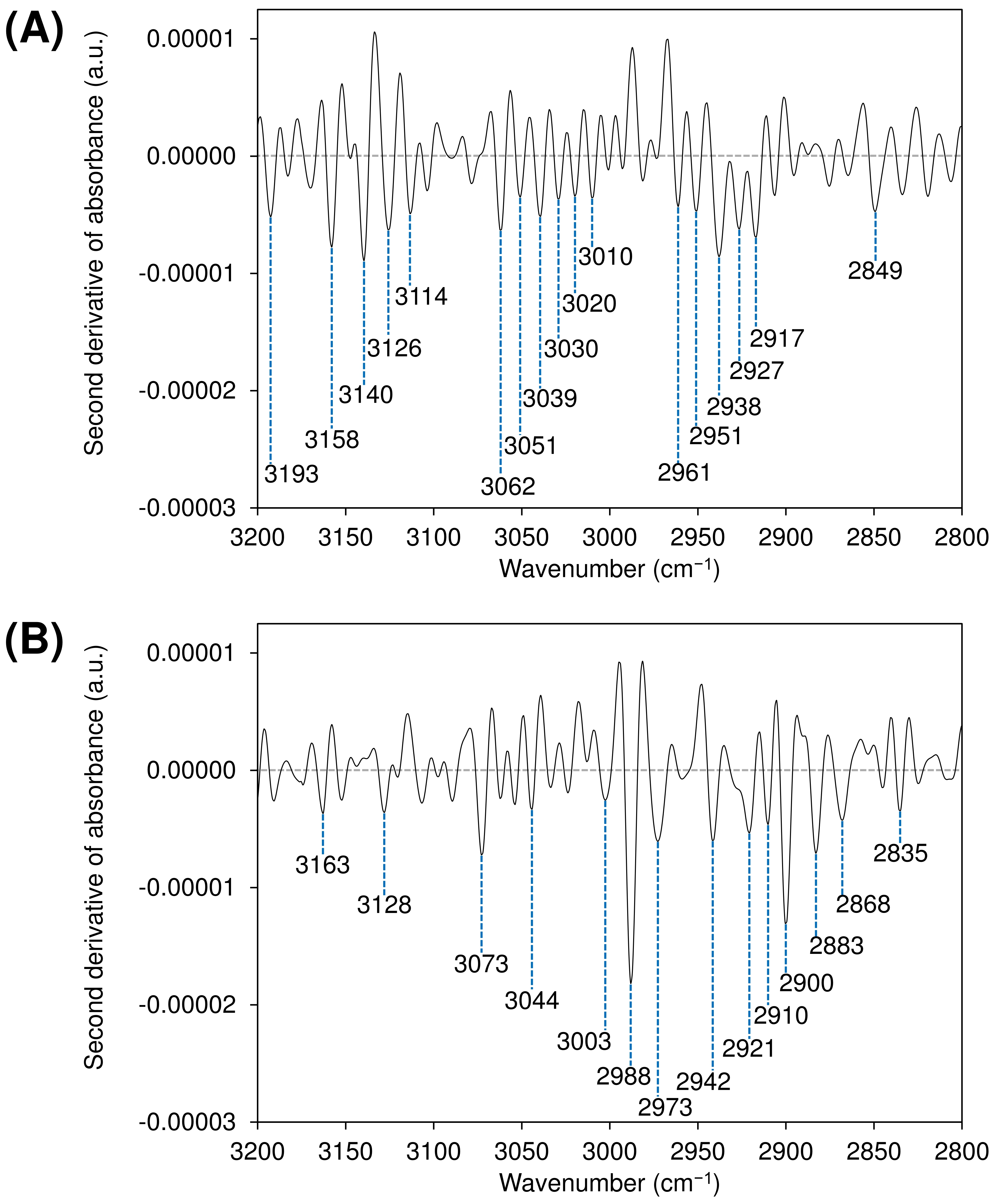

**Fig. S23.** Second-derivative ATR-FTIR spectra of (A) SKP-G1 and (B) SKP-G2 in the 2800–3200 cm^–1^ region. The blue vertical lines indicate the wavenumbers of selected negative second-derivative bands.

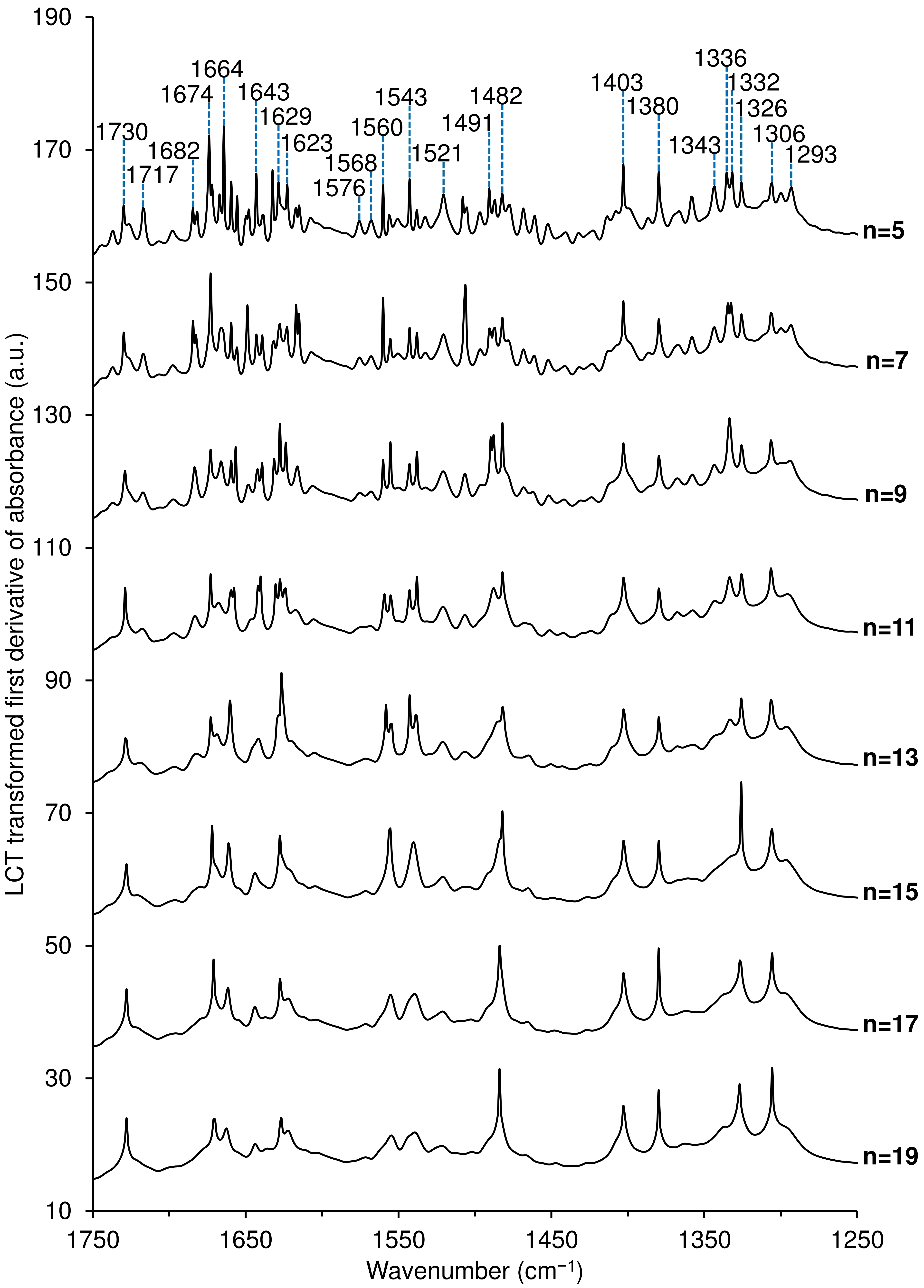

**Fig. S24.** LCT-transformed first-derivative ATR-FTIR spectra of SKP-G1 in the 1250–1750 cm^–1^ region, with the lowercase letter n denoting the number of Savitzky-Golay smoothing points, increasing from 5 to 19.

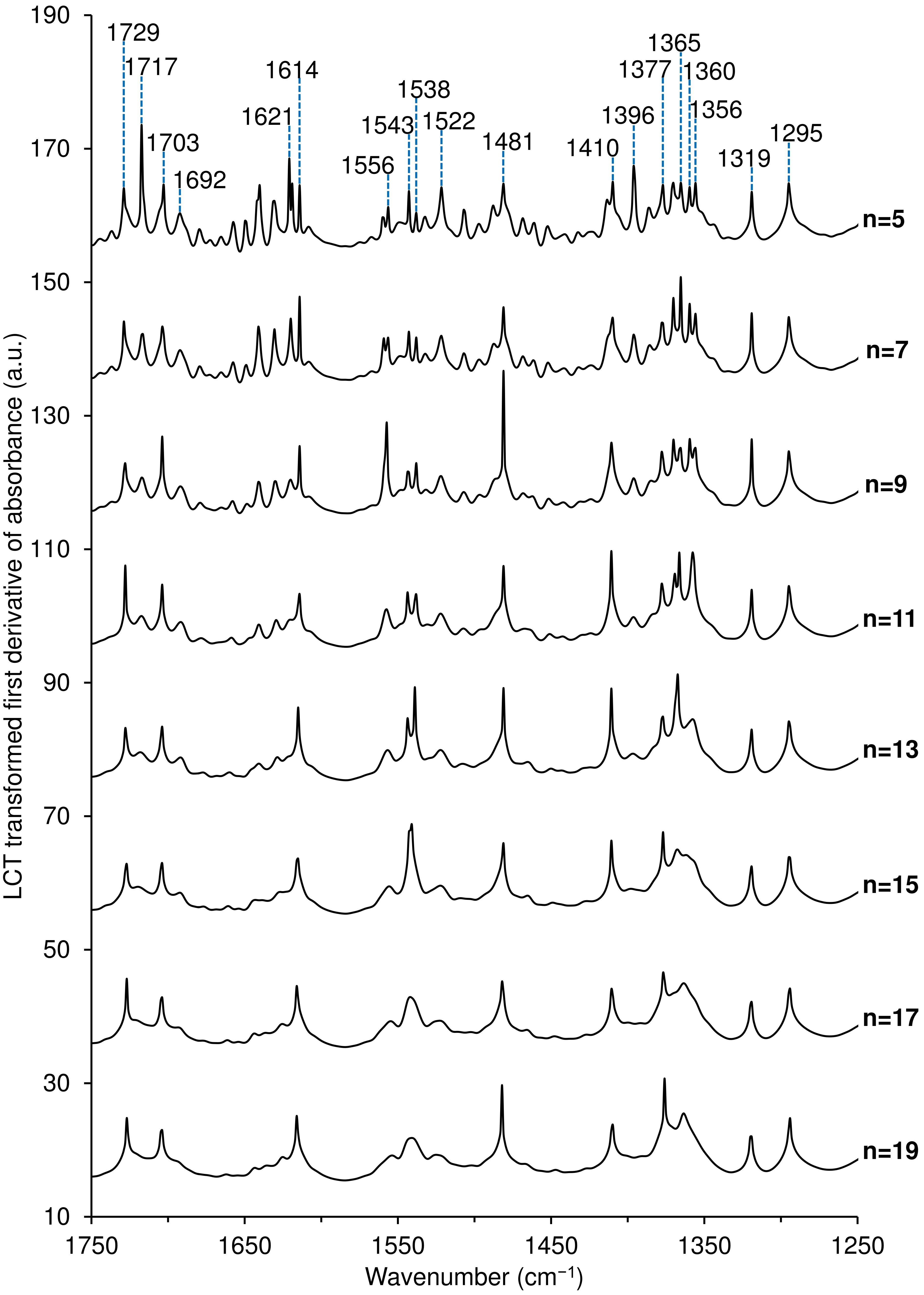

**Fig. S25.** LCT-transformed first-derivative ATR-FTIR spectra of SKP-G2 in the 1250–1750 cm^–1^ region, with the lowercase letter n denoting the number of Savitzky-Golay smoothing points, increasing from 5 to 19.

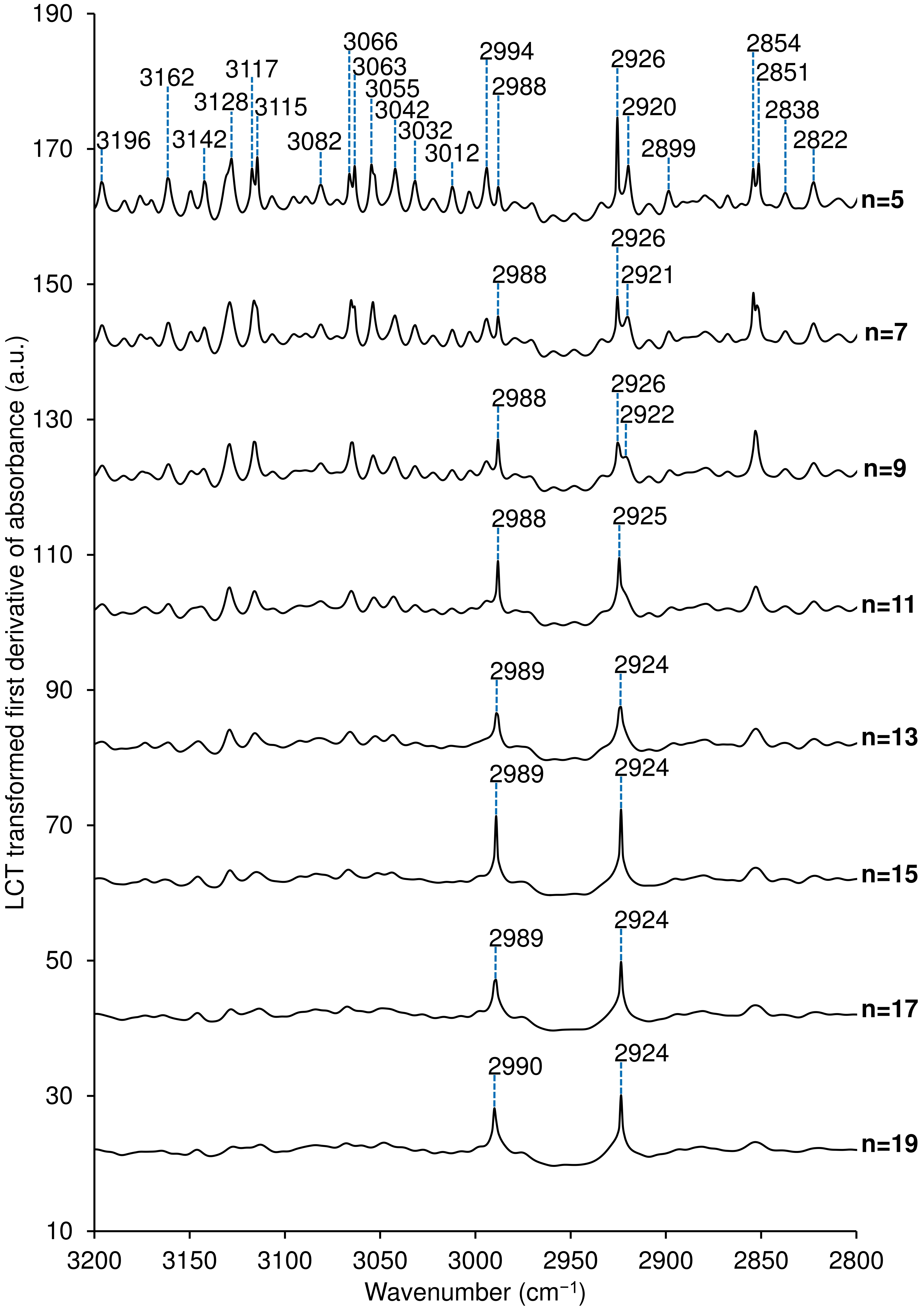

**Fig. S26.** LCT-transformed first-derivative ATR-FTIR spectra of SKP-G1 in the 2800–3200 cm^–1^ region, with the lowercase letter n denoting the number of Savitzky-Golay smoothing points, increasing from 5 to 19.

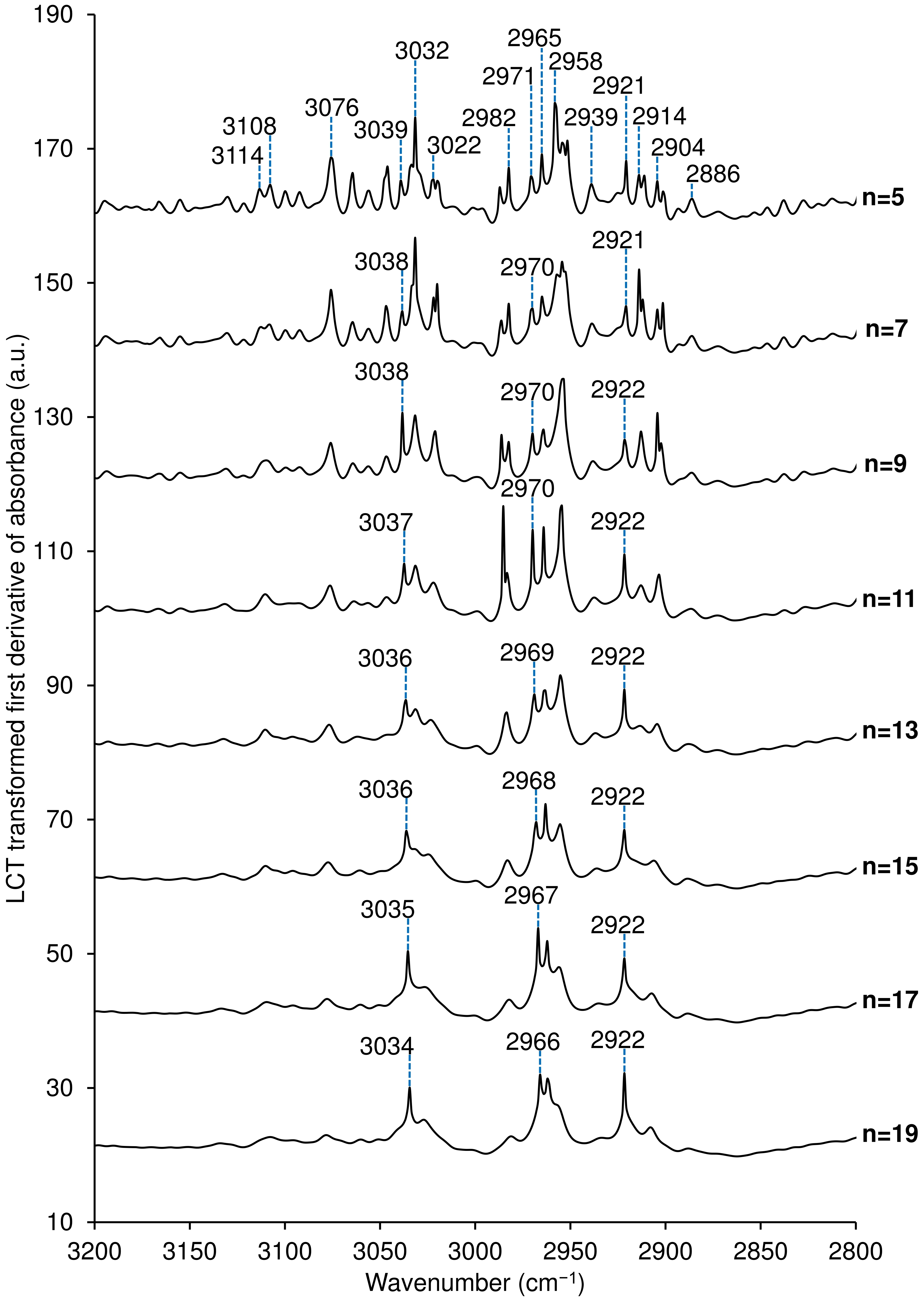

**Fig. S27.** LCT-transformed first-derivative ATR-FTIR spectra of SKP-G2 in the 2800–3200 cm^–1^ region, with the lowercase letter n denoting the number of Savitzky-Golay smoothing points, increasing from 5 to 19.

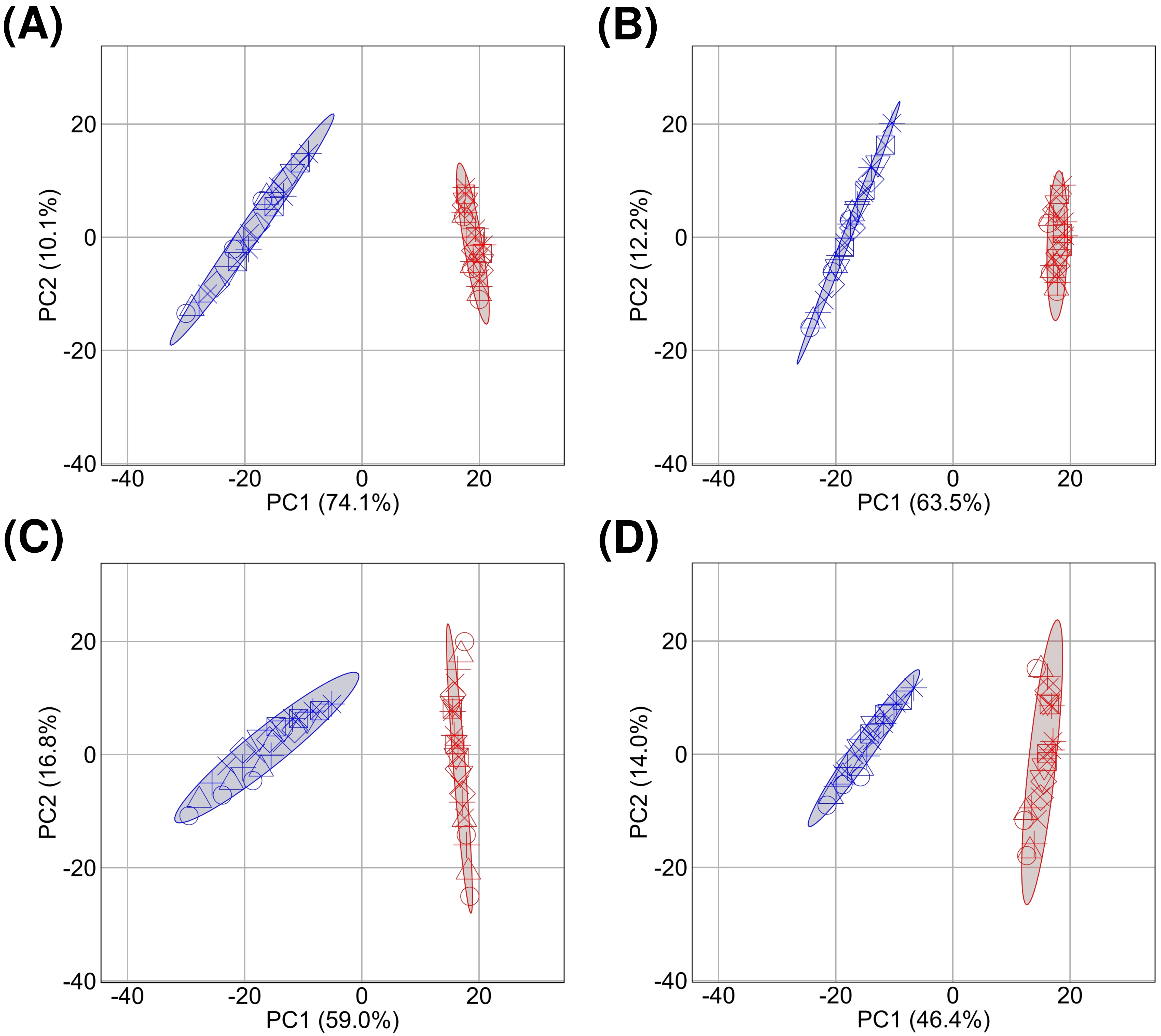

**Fig. S28.** PCA score plots for SKP-G1 (blue) and SKP-G2 (red) based on (A) first-derivative ATR-FTIR spectra, (B) LCT-transformed first-derivative ATR-FTIR spectra, (C) second-derivative ATR-FTIR spectra, and (D) LCT-transformed second-derivative ATR-FTIR spectra. Each point is plotted using one of eight marker shapes representing Savitzky-Golay smoothing windows ranging from 5 to 19 points: circle (○), upright triangle (△), plus sign (+), diagonal cross (×), diamond (◇), inverted triangle (▽), square with diagonal cross (☒), and eight-spoked asterisk (✳︎), respectively. The 95% data ellipses are calculated from the pooled PCA scores across all eight smoothing windows for each fraction.

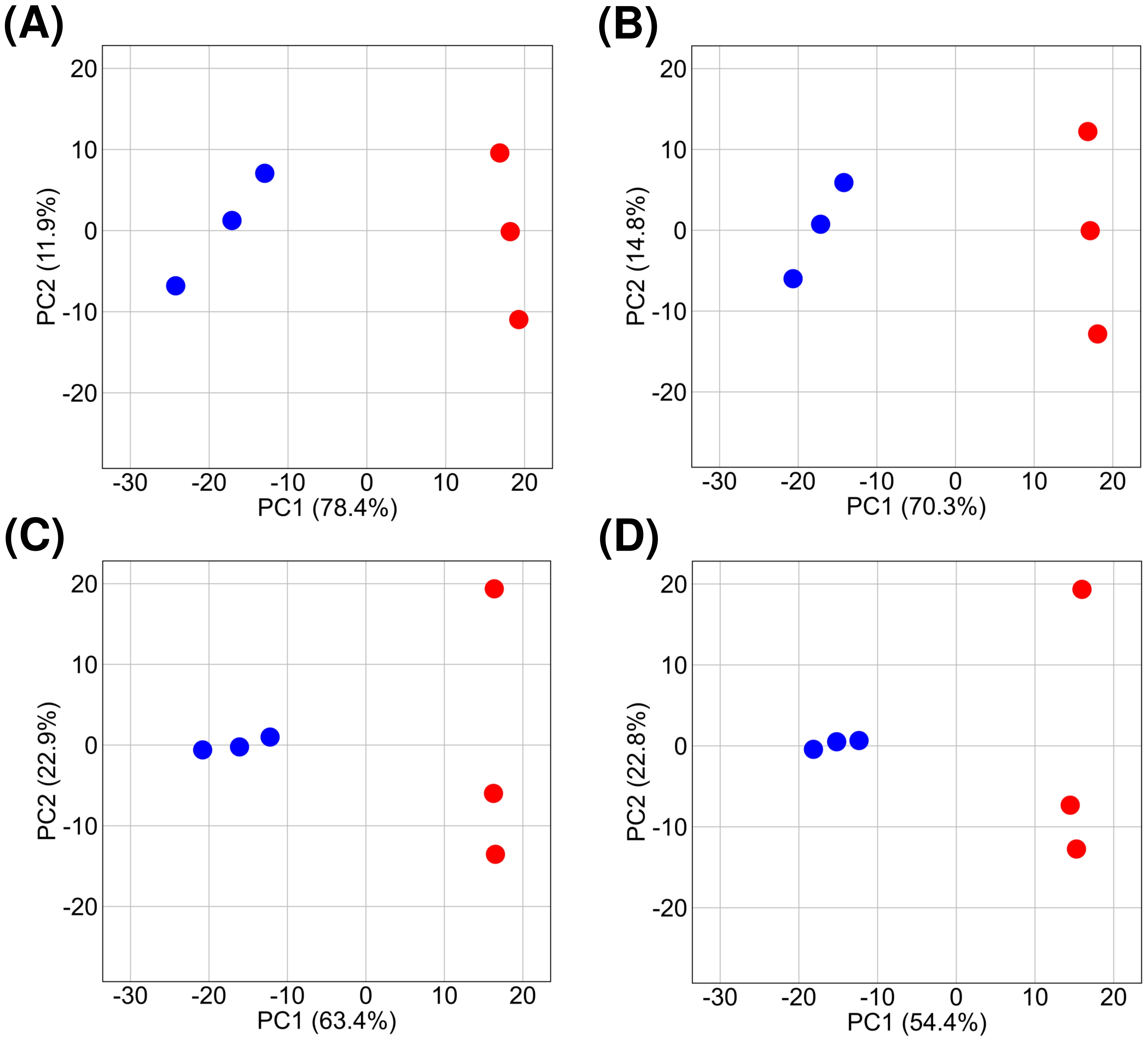

**Fig. S29.** PCA score plots of SKP-G1 (blue) and SKP-G2 (red) based on (A) first-derivative ATR-FTIR spectra, (B) LCT-transformed first-derivative ATR-FTIR spectra, (C) second-derivative ATR-FTIR spectra, and (D) LCT-transformed second-derivative ATR-FTIR spectra. All derivative analyses were performed using a 5-point Savitzky-Golay smoothing window. Three independent experiments were conducted for each sample, resulting in three points per sample in each plot.

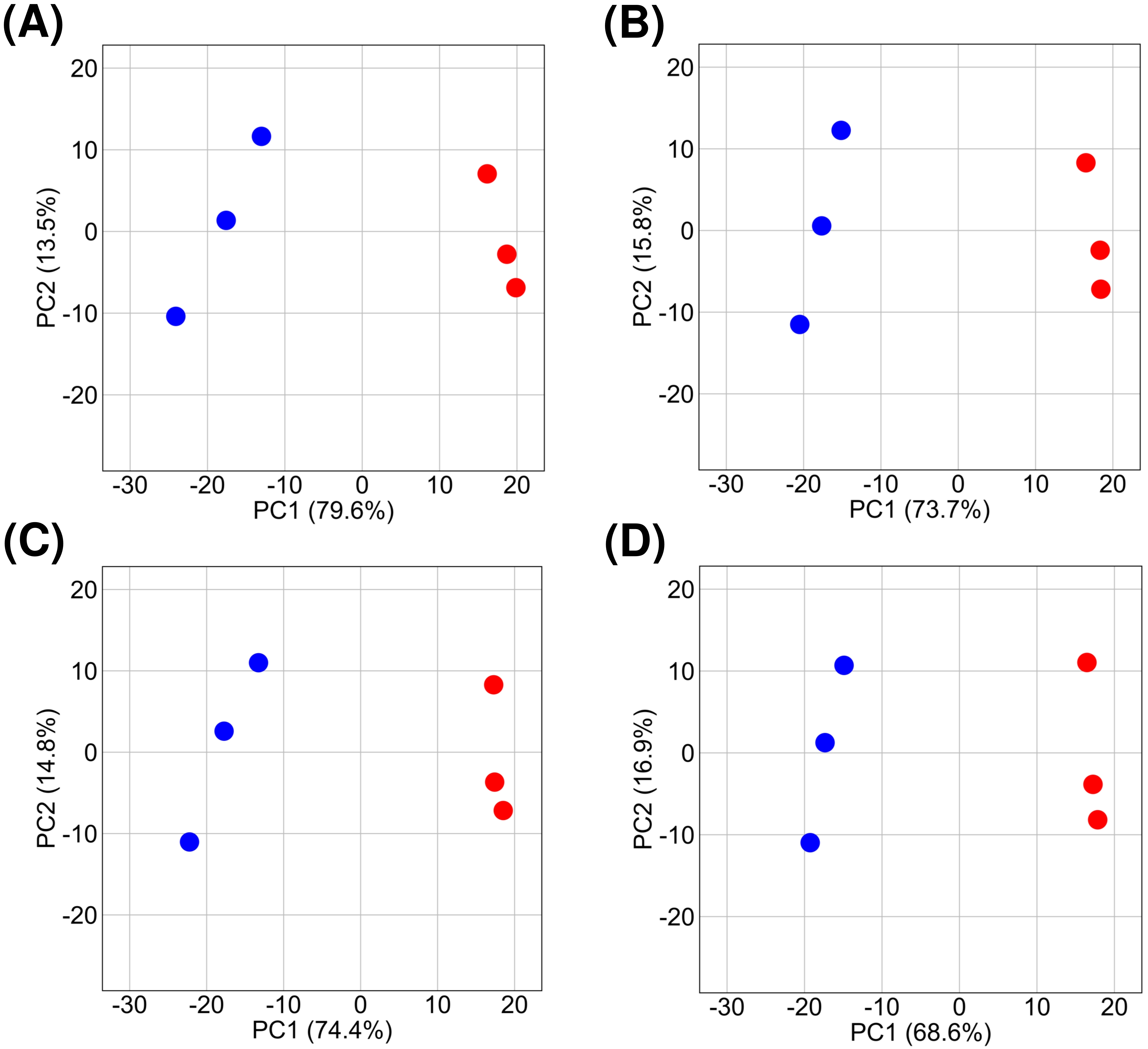

**Fig. S30.** PCA score plots of SKP-G1 (blue) and SKP-G2 (red) based on (A) first-derivative ATR-FTIR spectra, (B) LCT-transformed first-derivative ATR-FTIR spectra, (C) second-derivative ATR-FTIR spectra, and (D) LCT-transformed second-derivative ATR-FTIR spectra. All derivative analyses were performed using a 19-point Savitzky-Golay smoothing window. Three independent experiments were conducted for each sample, resulting in three points per sample in each plot.

**Table S1.** Weights and yields of AIR prepared from dry sugar kelp sample

| Tube No. | Dry weights (g) | | Yield (%, dry mass basis) |
| --- | --- | --- | --- |
|  | Starting material | AIR |  |
| #1 | 3.100 | 1.935 | 62.4 |
| #2 | 3.185 | 1.942 | 61.0 |
| #3 | 3.132 | 1.895 | 60.5 |
| #4 | 3.262 | 2.017 | 61.8 |
| #5 | 3.377 | 2.046 | 60.6 |
| #6 | 3.245 | 1.975 | 60.9 |
| #7 | 3.317 | 2.086 | 62.9 |
| #8 | 3.289 | 2.030 | 61.7 |
| #9 | 3.214 | 1.942 | 60.4 |
| #10 | 3.496 | 2.101 | 60.1 |
| #11 | 3.300 | 2.038 | 61.8 |
| #12 | 3.427 | 2.075 | 60.5 |
| #13 | 3.234 | 1.946 | 60.2 |
| #14 | 3.334 | 2.045 | 61.3 |
| #15 | 3.403 | 2.055 | 60.4 |
| #16 | 3.548 | 2.159 | 60.9 |
| #17 | 3.509 | 2.167 | 61.8 |
| #18 | 3.215 | 1.938 | 60.3 |
| #19 | 3.253 | 2.003 | 61.6 |
| Mean |  |  | 61.1 |
| Standard deviation |  |  | 0.8 |

Note: AIR samples were prepared in nineteen 50 mL tubes, numbered #1–#19, as detailed in Section 2.2 of the main text.

**Table S2.** Relative peak areas (area %) of fucose, GulA, and ManA in SKP-G1 and SKP-G2

| Monosaccharides | SKP-G1 | | |  | SKP-G2 | | |
| --- | --- | --- | --- | --- | --- | --- | --- |
|  | R1 | R2 | Mean |  | R1 | R2 | Mean |
| Fuc | 3.12 | 2.31 | 2.71 |  | 14.83 | 13.72 | 14.27 |
| GulA | 46.48 | 58.04 | 52.26 |  | 36.49 | 26.42 | 31.45 |
| ManA | 50.40 | 39.65 | 45.02 |  | 48.69 | 59.86 | 54.27 |

Note: R1 and R2 represent two independent experiments conducted for each sample, from which the mean values were calculated.

**Table S3.** Estimated monosaccharide compositions (mol%) of alginates quantified by GC-MS/FID analysis of acetate derivatives of monosaccharide alditols and 1,6-anhydrogulose generated from depolymerized SKP-G1 and SKP-G2

| Monomer | SKP-G1 | | |  | SKP-G2 | | | Acetate derivatives |
| --- | --- | --- | --- | --- | --- | --- | --- | --- |
|  | R1 | R2 | Mean |  | R1 | R2 | Mean |  |
| Ara | 0.5 | 0.5 | 0.5 |  | 0.5 | 0.6 | 0.5 | 1-deutero-arabitol |
| Fuc | 1.3 | 1.2 | 1.2 |  | 7.0 | 8.0 | 7.5 | 1-deutero-fucitol |
| Rha | t.r. | t.r. | t.r. |  | t.r. | t.r. | t.r. | 1-deutero-rhamnitol |
| Xyl | 0.6 | 0.6 | 0.6 |  | 0.8 | 1.0 | 0.9 | 1-deutero-xylitol |
| Gal | t.r. | t.r. | t.r. |  | 1.4 | 1.7 | 1.6 | 1-deutero-galactitol |
| Glc | 1.2 | 1.3 | 1.2 |  | 1.8 | 2.6 | 2.2 | 1-deutero-sorbitol |
| Man | 1.2 | 1.2 | 1.2 |  | 2.8 | 3.0 | 2.9 | 1-deutero-mannitol |
| GalA | 0.7 | 0.7 | 0.7 |  | 0.8 | 0.8 | 0.8 | 1,6,6′-trideutero-galactitol |
| GlcA | t.r. | t.r. | t.r. |  | t.r. | t.r. | t.r. | 1,6,6′-trideutero-sorbitol |
| GulA | 68.0 | 68.0 | 68.0 |  | 44.3 | 44.4 | 44.4 | 1,1′,6-trideutero-sorbitol and 1,6-anhydro-Gul*p* |
| ManA | 26.2 | 26.2 | 26.2 |  | 40.3 | 37.4 | 38.9 | 1,6,6′-trideutero-mannitol |

Note: R1 and R2 represent two independent experiments conducted for each sample, from which the mean values were calculated. 1,6,6′-Trideutero-sorbitol derived from GlcA and 1,1′,6-trideutero-sorbitol derived from GulA are both sorbitols that differ only in the positions of deuteration; therefore, their acetate derivatives coelute on GC and cannot be distinguished by FID. Because only a trace amount of GlcA was detected by HPAEC-PAD, for estimation purposes, the peak of peracetylated sorbitol was assigned exclusively to 1,1′,6-trideutero-sorbitol derived from GulA. t.r. denotes a trace amount (mol% < 0.5). Relative response factors of 0.745, 0.890, 0.835, and 0.535 were calculated using the effective carbon number concept for the acetate derivatives of pentitol, hexitol, 6-deoxyhexitol, and 1,6-anhydro-Gul*p*, respectively [1, 2].

**Table S4.** Results of methylation-GC-MS analysis of SKP-G1 and SKP-G2 subjected to mild methanolysis (0.5 M HCl in methanol, 80 °C, 20 min), followed by NaBD_4_ reduction

| Linkages | SKP-G1 | | |  | SKP-G2 | | |
| --- | --- | --- | --- | --- | --- | --- | --- |
|  | R1 | R2 | Mean |  | R1 | R2 | Mean |
| t-Fuc*p* | 0.5 | 1.5 | 1.0 |  | 1.6 | 3.3 | 2.4 |
| t-Gal*p* | t.r. | t.r. | t.r. |  | 0.6 | 0.8 | 0.7 |
| 4-Glc*p* | 1.7 | 0.5 | 1.1 |  | 8.1 | 1.7 | 4.9 |
| 2-Man*p* | 0.8 | 1.5 | 1.2 |  | 1.9 | 2.9 | 2.4 |
| 3-Glc*p*A | t.r. | t.r. | t.r. |  | 0.8 | 1.3 | 1.0 |
| 4-Glc*p*A | 0.5 | 0.6 | 0.6 |  | 1.6 | 1.8 | 1.7 |
| t-Gul*p*A | 1.7 | 4.9 | 3.3 |  | 0.7 | 1.5 | 1.1 |
| 4-Gul*p*A | 57.2 | 50.3 | 53.7 |  | 34.5 | 34.7 | 34.6 |
| t-Man*p*A | 3.7 | 5.1 | 4.4 |  | 3.4 | 5.3 | 4.3 |
| 4-Man*p*A | 33.3 | 34.6 | 33.9 |  | 46.8 | 46.8 | 46.8 |

Note: R1 and R2 represent two independent experiments conducted for each sample, from which the mean values were calculated. t.r. denotes a trace amount (mol% < 0.5). Before linkage analysis (methylation-GC-MS analysis), the samples were subjected to mild methanolysis (0.5 M HCl in methanol, 80 °C, 20 min), followed by NaBD_4_ reduction to convert uronic acids to their corresponding 6,6′-dideuterated neutral sugars [3-6]. The relative molar compositions (mol%) of the PMAAs were estimated by dividing the TIC peak area of each PMAA by its molecular mass, in accordance with the published protocol [7].

**Table S5.** Results of HPSEC analysis of **SKP-G1** and **SKP-G2**

| Parameters |  | **SKP-G1** |  |  |  | **SKP-G2** |  |
| --- | --- | --- | --- | --- | --- | --- | --- |
|  | R1 | R2 | Mean |  | R1 | R2 | Mean |
| **Rh(Q)z**  (nm) | 63.255  (±41.376%) | 66.956  (±44.314%) | 65.106 |  | 40.032  (±51.564%) | 72.254  (±32.329%) | 56.143 |
| **Rh(v)z**  (nm) | 16.248  (±0.359%) | 16.530  (±0.590%) | 16.389 |  | 12.908  (±0.456%) | 12.869  (±0.498%) | 12.889 |
| **Mn**  (g/mol) | 2.146×10^4^  (±1.130%) | 2.312×10^4^  (±1.678%) | 2.229×10^4^ |  | 2.432×10^4^  (±1.522%) | 2.332×10^4^  (±1.333%) | 2.382×10^4^ |
| **Mw**  (g/mol) | 3.971×10^4^  (±1.011%) | 4.192×10^4^  (±1.649%) | 4.082×10^4^ |  | 3.973×10^4^  (±1.329%) | 3.721×10^4^  (±1.359%) | 3.847×10^4^ |
| **Mw/Mn** | 1.850  (±1.516%) | 1.813  (±2.353%) | 1.8315 |  | 1.634  (±2.021%) | 1.595  (±1.903%) | 1.615 |
| **Rgz**  (nm) | 25.9  (±5.8%) | 34.2  (±5.5%) | 30.1 |  | 19.0  (±13.8%) | 17.9  (±16.6%) | 18.5 |
| **[η]n**  (mL/g) | 99.904  (±0.184%) | 102.441  (±0.250%) | 101.173 |  | 68.024  (±0.253%) | 69.529  (±0.305%) | 68.777 |
| **[η]w**  (mL/g) | 179.14  (±0.20%) | 180.55  (±0.28%) | 179.85 |  | 106.06  (±0.30%) | 107.56  (±0.36%) | 106.81 |
| **[η]z**  (mL/g) | 367.862  (±0.272%) | 368.623  (±0.414%) | 368.243 |  | 205.657  (±0.409%) | 216.685  (±0.493%) | 211.171 |
| **MHS a** | 0.970  (±0.007%) | 0.979  (±0.010%) | 0.975 |  | 0.925  (±0.014%) | 0.948  (±0.016%) | 0.937 |
| **MHS K**  (mL/g) | 6.550×10^-3^  (±0.075%) | 5.693×10^-3^  (±0.108%) | 6.122×10^-3^ |  | 6.121×10^-3^  (±0.138%) | 5.114×10^-3^  (±0.164%) | 5.618×10^-3^ |

**Note:** The terms in the table are defined as follows: **Rh(Q)z**, the Z‑average hydrodynamic radius determined by DLS, where Q denotes quasi‑elastic light scattering (QELS), an alternative term for DLS; **Rh(v)z**, the Z‑average hydrodynamic radius determined by viscometry; **Mn** and  **Mw**, the number‑average and weight‑average molecular weights, respectively; **Rgz**, the Z‑average radius of gyration; **[η]n** , **[η]w**, and **[η]z**, the number‑average, weight‑average, and Z‑average intrinsic viscosities, respectively; and **MHS**  **a** and **MHS K**, the Mark–Houwink–Sakurada parameters a and K, respectively. The ratio  **Mw/Mn** denotes the polydispersity index. Bracketed percentage values indicate the relative uncertainties in each measurement, as calculated by the ASTRA software.

**Table S6**. Monad, diad, and triad frequencies, M/G ratio, and parameter η for SKP-G1 and SKP-G2, as determined by 1D ^1^H NMR

| Samples | Separate experiments | *F*_G_ | *F*_M_ | *F*_GG_ | *F*_MG_  *F*_GM_ | *F*_MM_ | *F*_GGG_ | *F*_GGM_  *F*_MGG_ | *F*_MGM_ | *F*_MMM_ | *F*_MMG_  *F*_GMM_ | *F*_GMG_ | M/G | η |
| --- | --- | --- | --- | --- | --- | --- | --- | --- | --- | --- | --- | --- | --- | --- |
| **SKP-G1** |  |  |  |  |  |  |  |  |  |  |  |  |  |  |
| NF65 | R1 | 0.73 | 0.27 | 0.58 | 0.15 | 0.11 | 0.52 | 0.06 | 0.09 | [0.00, 0.11] | [0.00, 0.11] | [0.00, 0.15] | 0.37 | 0.79 |
|  | R2 | 0.77 | 0.23 | 0.63 | 0.14 | 0.10 | 0.58 | 0.05 | 0.08 | [0.00, 0.10] | [0.00, 0.10] | [0.00, 0.14] | 0.31 | 0.76 |
| NF80 | R1 | 0.75 | 0.25 | 0.60 | 0.15 | 0.10 | 0.56 | 0.04 | 0.11 | [0.00, 0.10] | [0.00, 0.10] | [0.00, 0.15] | 0.33 | 0.80 |
|  | R2 | 0.75 | 0.25 | 0.61 | 0.15 | 0.10 | 0.56 | 0.05 | 0.10 | [0.00, 0.10] | [0.00, 0.10] | [0.00, 0.15] | 0.33 | 0.79 |
| GW80 | R1 | 0.72 | 0.28 | 0.56 | 0.16 | 0.12 | 0.51 | 0.05 | 0.11 | [0.00, 0.12] | [0.00, 0.12] | [0.00, 0.16] | 0.39 | 0.81 |
|  | R2 | 0.74 | 0.26 | 0.57 | 0.16 | 0.10 | 0.53 | 0.05 | 0.11 | [0.00, 0.10] | [0.00, 0.10] | [0.00, 0.16] | 0.36 | 0.83 |
| **SKP-G2** |  |  |  |  |  |  |  |  |  |  |  |  |  |  |
| NF65 | R1 | 0.31 | 0.69 | 0.22 | 0.09 | 0.60 | 0.22 | 0.00 | 0.09 | [0.51, 0.60] | [0.00, 0.09] | [0.00, 0.09] | 2.24 | 0.44 |
|  | R2 | 0.30 | 0.70 | 0.20 | 0.10 | 0.60 | 0.20 | 0.00 | 0.10 | [0.50, 0.60] | [0.00, 0.10] | [0.00, 0.10] | 2.31 | 0.46 |
| NF80 | R1 | 0.29 | 0.71 | 0.19 | 0.10 | 0.61 | 0.19 | 0.00 | 0.10 | [0.51, 0.61] | [0.00, 0.10] | [0.00, 0.10] | 2.41 | 0.48 |
|  | R2 | 0.28 | 0.72 | 0.19 | 0.09 | 0.63 | 0.19 | 0.00 | 0.09 | [0.54, 0.63] | [0.00, 0.09] | [0.00, 0.09] | 2.60 | 0.45 |
| GW80 | R1 | 0.29 | 0.71 | 0.19 | 0.10 | 0.61 | 0.19 | 0.00 | 0.10 | [0.52, 0.61] | [0.00, 0.10] | [0.00, 0.10] | 2.46 | 0.48 |
|  | R2 | 0.28 | 0.72 | 0.19 | 0.09 | 0.62 | 0.19 | 0.00 | 0.09 | [0.53, 0.62] | [0.00, 0.09] | [0.00, 0.09] | 2.54 | 0.47 |

**Note:** R1 and R2 represent values obtained from two independent experiments conducted for each sample. [a, b] denotes a closed interval on the real number line, including all values $x$ such that $a\leq x\leq b$. The M-to-G ratio (M/G) was calculated as ${F_{M}}/{F_{G}}$, and parameter η was calculated as ${F_{\mathrm{GM}}}/\left( F_{G}\times F_{M} \right)$[8]. “NF65,” “NF80,” and “GW80” denote spectra collected at 65 °C without filtration, at 80 °C without filtration, and at 80 °C with glass wool filtration, respectively.

**Table S7.** Summary of ^1^H NMR chemical shifts (δ, ppm) at 65 °C and 80 °C for detected minor reducing end residues of alginates

| Residue | Assignment | T | δ(H-1) | δ(H-2) | δ(H-3) | δ(H-4) | δ(H-5) |
| --- | --- | --- | --- | --- | --- | --- | --- |
| Gα | →4)-α-l-Gul*p*A | 65 °C | 5.21 | 3.92 | 4.33 | – | – |
|  |  | 80 °C | 5.21 | 3.92 | 4.33 | – | – |
| Gβ | →4)-β-l-Gul*p*A | 65 °C | 4.89 | 3.62 | 4.16 | – | – |
|  |  | 80 °C | 4.89 | 3.62 | 4.15 | – | – |
| Mβ | →4)-β-d-Man*p*A | 65 °C | 4.86 | 3.71 | – | – | – |
|  |  | 80 °C | 4.86 | 3.70 | – | – | – |

Note: The symbol “–” indicates that chemical shifts were not obtained because the signals from the corresponding protons in the minor residues were too weak to detect. Chemical shifts were referenced to TSP-d_4_ at 0 ppm in D_2_O.

**R script for making PCA score plot (Fig. 14)**

\# Load required libraries

library(dplyr)

library(forcats)

library(ggplot2)

\# Insert the name of your file here

data <- read.csv("PCAMike2ndderLCT-threePs.csv")

\# Perform principal component analysis (PCA), scale = TRUE will subtract mean and divide by standard deviation

pca <- prcomp(data[, -1], scale = TRUE)

\# Create a data frame with PCA results

pca_data <- as.data.frame(pca$x[, 1:2])

pca_data$Label <- data[, 1]

\# Assign groups (Rows 1-9 = Group A, Rows 10-18 = Group B)

pca_data$Group <- ifelse(as.numeric(rownames(pca_data)) %in% 1:9, "G1", "G2")

\# Assign shared shapes to specific rows

pca_data$Shape <- as.character(pca_data$Label) # Start with unique labels

pca_data$Shape[as.numeric(rownames(pca_data)) %in% c(1:3, 10:12)] <- "p5"

pca_data$Shape[as.numeric(rownames(pca_data)) %in% c(4:6, 13:15)] <- "p11"

pca_data$Shape[as.numeric(rownames(pca_data)) %in% c(7:9, 16:18)] <- "p19"

\# Define custom shapes, assigning a diamond shape (5) to "p19"

shape_values <- setNames(c(1, 2, 5), c("p5", "p11", "p19"))

\# Extract variance explained by each principal component

variance_explained <- round(pca$sdev^2 / sum(pca$sdev^2) * 100, 2)

variance_explained <- sprintf("%.1f", variance_explained)

\# Flip the y-axis

pca_data$PC2 <- -pca_data$PC2

\# Create PCA plot using ggplot2

pca_plot <- ggplot(pca_data, aes(x = PC1, y = PC2, color = Group, shape = Shape)) +

geom_point(size = 10) +

stat_ellipse(aes(group = Group, fill = Group), alpha = 0.2, geom = "polygon", level = 0.95) + # Grouped ellipses

scale_color_manual(values = c("G1" = "blue", "G2" = "red")) + # Set custom colours

scale_fill_manual(values = c("G1" = "blue", "G2" = "red"), guide = "none") + # Match fill colours

scale_shape_manual(values = shape_values) + # Assign custom shapes

labs(

x = paste("PC1 (", variance_explained[1], "%)", sep = ""),

y = paste("PC2 (", variance_explained[2], "%)", sep = ""),

color = "Group",

shape = "Sample"

) +

xlim(-41, 31) + # Set x-axis range

ylim(-37, 30.5) + # Set y-axis range

theme_minimal() +

theme(

legend.key.size = unit(0.7, "cm"),

legend.text = element_text(size = 12, face = "italic"),

legend.title = element_text(size = 14),

axis.text = element_text(color = "black", size = 28),

axis.title = element_text(size = 28),

panel.grid.major = element_line(color = "gray", size = 0.5),

panel.grid.minor = element_blank(),

panel.border = element_rect(color = "black", fill = NA, size = 1)

)

\# Display the plot

print(pca_plot)

\# Save PCA plot as PNG

ggsave("PCAMike2ndderLCT-threePs.png", plot = pca_plot, width = 10, height = 8, units = "in", dpi = 900)

**R script for making PCA score plot (Fig. S28)**

\# Load required libraries

library(dplyr)

library(forcats)

library(ggplot2)

\# Insert the name of your file here

data <- read.csv("PCAMike2ndderLCT.csv")

\# Perform principal component analysis (PCA), scale = TRUE will subtract mean and divide by standard deviation

pca <- prcomp(data[, -1], scale = TRUE)

\# Create a data frame with PCA results

pca_data <- as.data.frame(pca$x[, 1:2])

pca_data$Label <- data[, 1]

\# Assign colours to groups (Rows 1-24 = Group A, Rows 25-48 = Group B)

pca_data$Group <- ifelse(rownames(pca_data) %in% 1:24, "G1", "G2")

\# Assign shared shapes to specific rows

pca_data$Shape <- as.character(pca_data$Label) # Start with unique labels

pca_data$Shape[as.numeric(rownames(pca_data)) %in% c(1:3, 25:27)] <- "p5"

pca_data$Shape[as.numeric(rownames(pca_data)) %in% c(4:6, 28:30)] <- "p7"

pca_data$Shape[as.numeric(rownames(pca_data)) %in% c(7:9, 31:33)] <- "p9"

pca_data$Shape[as.numeric(rownames(pca_data)) %in% c(10:12, 34:36)] <- "p11"

pca_data$Shape[as.numeric(rownames(pca_data)) %in% c(13:15, 37:39)] <- "p13"

pca_data$Shape[as.numeric(rownames(pca_data)) %in% c(16:18, 40:42)] <- "p15"

pca_data$Shape[as.numeric(rownames(pca_data)) %in% c(19:21, 43:45)] <- "p17"

pca_data$Shape[as.numeric(rownames(pca_data)) %in% c(22:24, 46:48)] <- "p19"

\# Define custom shapes

unique_shapes <- unique(pca_data$Shape)

shape_values <- setNames(1:length(unique_shapes), unique_shapes) # Assign unique shapes

\# Extract variance explained by each principal component

variance_explained <- round(pca$sdev^2 / sum(pca$sdev^2) * 100, 2)

variance_explained <- sprintf("%.1f", variance_explained)

\# Flip the y-axis

pca_data$PC2 <- -pca_data$PC2

\# Create PCA plot using ggplot2

pca_plot <- ggplot(pca_data, aes(x = PC1, y = PC2, color = Group, shape = Shape)) +

geom_point(size = 10) +

stat_ellipse(aes(group = Group, fill = Group), alpha = 0.2, geom = "polygon", level = 0.95) + # Grouped ellipses

scale_color_manual(values = c("G1" = "blue", "G2" = "red")) + # Set custom colours

scale_fill_manual(values = c("G1" = "blue", "G2" = "red"), guide = "none") + # Match fill colours

scale_shape_manual(values = shape_values) + # Assign custom shapes

labs(

x = paste("PC1 (", variance_explained[1], "%)", sep = ""),

y = paste("PC2 (", variance_explained[2], "%)", sep = ""),

color = "Group",

shape = "Sample"

) +

xlim(-41, 31) + # Set x-axis range

ylim(-37, 30.5) + # Set y-axis range

theme_minimal() +

theme(

legend.key.size = unit(0.7, "cm"),

legend.text = element_text(size = 12, face = "italic"),

legend.title = element_text(size = 14),

axis.text = element_text(color = "black", size = 28),

axis.title = element_text(size = 28),

panel.grid.major = element_line(color = "gray", size = 0.5),

panel.grid.minor = element_blank(),

panel.border = element_rect(color = "black", fill = NA, size = 1)

)

\# Display the plot

print(pca_plot)

\# Save PCA plot as PNG

ggsave("PCAMike2ndderLCT.png", plot = pca_plot, width = 10, height = 8, units = "in", dpi = 900)

**R script for making PCA score plot (Figs. S29 and S30)**

\# Load required libraries

library(dplyr) # For data manipulation

library(forcats) # For further data manipulation

library(ggplot2) # For plotting

\# Insert the name of your file here

data <- read.csv("PCAMike2ndderLCT-p5.csv")

\# For Fig. S30, use the corresponding p19 CSV file and change the output PNG filename.

\# Perform principal component analysis (PCA), scale=true will subtract mean and divide by SDev

pca <- prcomp(data[, -1], scale = TRUE)

\# Create a data frame with PCA results

pca_data <- as.data.frame(pca$x[, 1:2])

pca_data$Label <- as.factor(data[, 1]) # Ensure Label is a factor for colour mapping

\# Display summary of PCA results

summary(pca)

biplot(pca, scale = 0)

\# Extract PC scores

str(pca)

pca$x

data2 <- cbind(data, pca$x[, 1:2])

head(data2)

\# Extract variance explained by each principal component

variance_explained <- round(pca$sdev^2 / sum(pca$sdev^2) * 100, 2)

\# Format variance explained to show one decimal place

variance_explained <- sprintf("%.1f", variance_explained)

\# Create PCA plot using ggplot2

pca_plot <- ggplot(pca_data, aes(x = PC1, y = PC2, color = Label, fill = Label)) +

geom_point(size = 10, shape = 16) +	# Adjust size and shape of dots

#stat_ellipse(alpha = 0.2) + # Add ellipses

labs(

x = paste("PC1 (", variance_explained[1], "%)", sep = ""), # X-axis label

y = paste("PC2 (", variance_explained[2], "%)", sep = ""), # Y-axis label

color = expression("Fraction"), # Set the legend title for the colour aesthetic

fill = expression("Fraction")	# Set the legend title for the fill aesthetic

) +

guides(

color = guide_legend(override.aes = list(shape = 16, size = 4)), #legend label shape and size

fill = guide_legend(override.aes = list(shape = 16, size = 4)) #legend label shape and size

) +

xlim(-31, 21) + # Set x-axis range

ylim(-27, 20.5) + # Set y-axis range

theme_minimal() +

theme(

legend.key.size = unit(0.7, "cm"), # Adjust the size of legend keys

legend.text = element_text(size = 24, face = "italic"), # Adjust the size of legend text, and make it italic

legend.title = element_text(size = 24), # Adjust the size of legend title

plot.background = element_rect(fill = "white"), # Set plot background color

plot.title = element_text(size = 16, hjust = 0.5), # Adjust plot title size and center it

axis.text = element_text(color="black", size = 28), # Adjust the font size for axis tick labels

axis.title = element_text(size = 28), # Adjust the font size for axis labels

panel.grid.major = element_line(color = "gray", size = 0.5), # Add major gridlines

panel.grid.minor = element_blank(), # Remove minor gridlines

panel.border = element_rect(color = "black", fill = NA, size = 1) # Add black border

) +

ggtitle("") # Add a title to the plot

\# Set the colour palette to red and blue

pca_plot <- pca_plot +

scale_color_manual(values = c("blue", "red")) + # Set colours for points

scale_fill_manual(values = c("blue", "red")) # Set colours for ellipses and legend

\# Display the plot

print(pca_plot)

\# Set output file name and save PCA plot as PNG

ggsave("PCAMike2ndderLCT-p5.png", plot = pca_plot, width = 10, height = 8, units = "in", dpi = 900)
